# Differential Kinetics of SARS-CoV-2 Proteases Revealed by a Dual-Color, BRET-based Protease Biosensor, DuProSense

**DOI:** 10.1101/2024.09.26.615113

**Authors:** Asma Fatima, Anupriya M Geethakumari, S M Nasir Uddin, Kabir H Biswas

## Abstract

While SARS-CoV-2 M^pro^ and PL^pro^ proteases are known to cleave polyproteins pp1a and pp1ab at multiple sites, these have not been comprehensively characterized in living cells. Here we engineered a two-color Bioluminescence Resonance Energy Transfer (BRET)-based, dual protease (DuProSense) biosensor platform relying on a proximity-dependent energy transfer from a luciferase donor to two spectrally separated fluorescent protein acceptors enabling simultaneous monitoring of processing of two cleavage sites in a single assay with high specificity. DuProSense revealed a similar M^pro^ and PL^pro^ cleavage kinetics for their N-terminal autocleavage sites. Importantly, systematic characterization of various M^pro^ and PL^pro^ cleavage sites using DuProSense revealed significant differences in cleavage rates and nirmatrelvir potency of M^pro^ cleavage sites but no correlation between the cleavage rates and nirmatrelvir IC_50_ values. Overall, our results provide deeper insights into the proteolytic processing of SARS-CoV-2 polyproteins and the dual color BRET platform will find wider applications in the future.

**Highlights:**

- Engineered a two-color BRET-based, dual protease biosensor (DuProSense)
- DuProSense biosensor enabled simultaneous and specific monitoring of M^pro^ and PL^pro^ activities
- DuProSense platform revealed differential cleavage kinetics of M^pro^ cleavage sites in live cells
- DuProSense platform revealed M^pro^ cleavage site-dependent nirmatrelvir potency in live cells

## Introduction

COVID-19, caused by SARS-CoV-2, has affected human health and the economy globally causing more than 7 million deaths and 750 million infections^1^. While the availability of vaccines has reduced the disease burden, recurrent emergence of SARS-CoV-2 variants with increased infection potential, disease severity, and resistance to antibody-mediated neutralization may make them ineffective.^2–4^ Therefore, the pharmacological targeting of SARS-CoV-2 proteins that are required for viral replication appears to be an attractive target for anti-COVID-19 therapy. Upon internalization of SARS-CoV-2, the viral genome is translated into two polyproteins, pp1a and pp1ab, consisting of various non-structural proteins (NSPs), in addition to other proteins. The SARS-CoV-2 infection cycle requires the cleavage of these polyproteins into functional proteins through the activity of two highly conserved viral cysteine proteases, main protease (M^pro^ or 3CL^pro^ or non-structural protein, NSP5) and papain-like protease (PL^pro^ or NSP3).^5–8^ The active homodimer M^pro^ cleaves the polyproteins at a total of 11 sites (present between NSP4 to NSP11 in pp1a and NSP4 to NSP16 in pp1ab) while PL^pro^ cleaves the polyproteins at a total of 3 sites (present between NSP1 to NSP4 in both pp1a and pp1ab) (Fig. 1A,B).^10^ Both M^pro^ and PL^pro^ show high specificity about their cleavage sites.^9,10^ For these reasons, the proteases have been suggested to be excellent targets for developing pharmacological agents for anti-SARS-CoV-2 therapy. Indeed, Paxlovid®, an FDA-approved anti-SARS-CoV-2 drug inhibits the activity of M^pro^.^7^ In addition, knowledge of the cleavage kinetics of proteases for all the cleavage sites is also important in drug design.

**Fig. 1.**
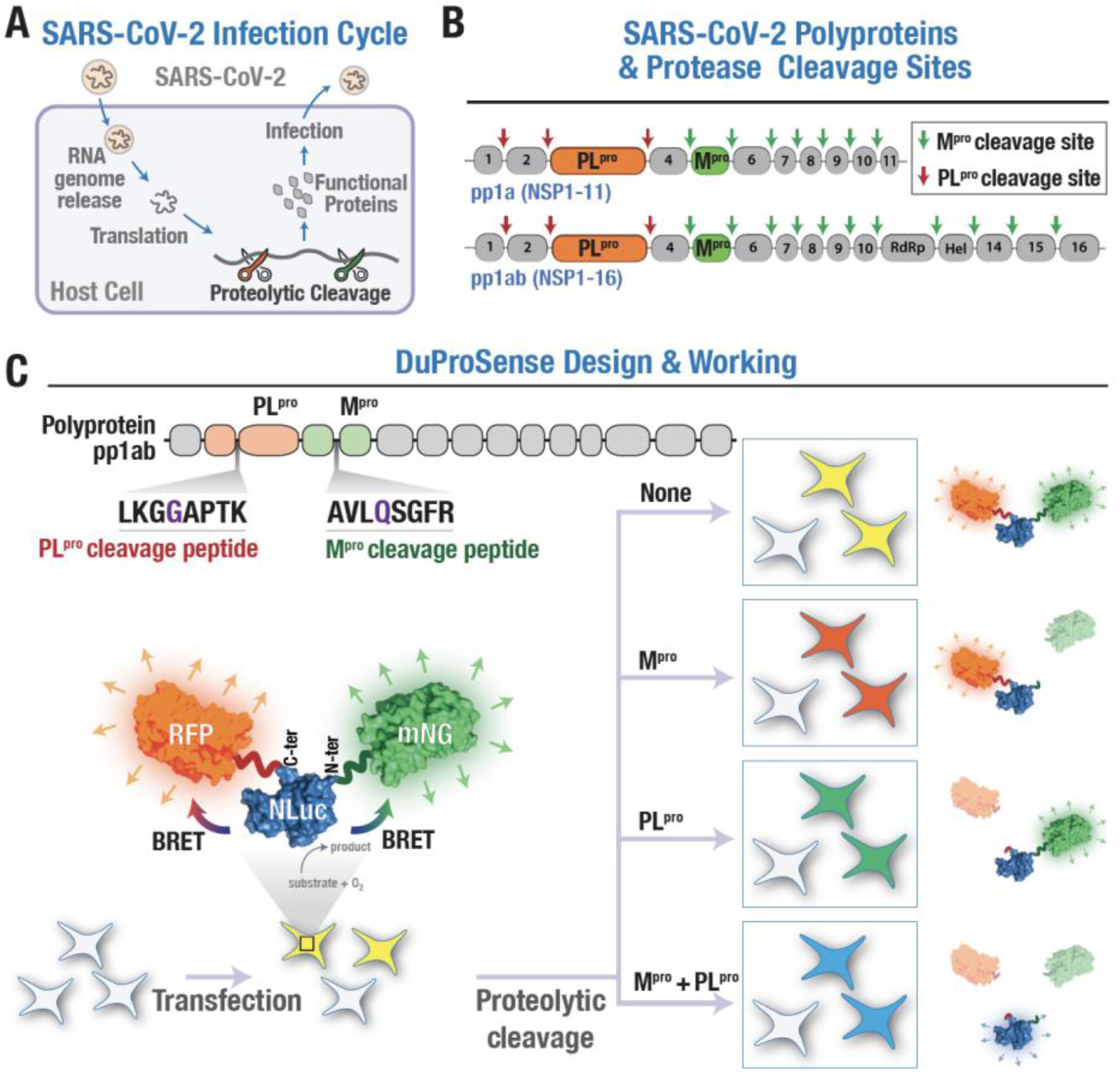
SARS-CoV-2 infection cycle, protease cleavage sites and BRET-based, dual protease biosensor (DuProSense biosensor). (A) Schematic showing the SARS-CoV-2 infection cycle in mammalian cells wherein the virus enters the cell, and releases its RNA genome, which is then translated into various proteins, including polyproteins. The polyproteins are then processed by the two viral proteases, M^pro^ and PL^pro^, into active proteins that enable the continuation of the infection cycle. (B) Schematic showing polyproteins, pp1a and pp1ab, and the cleavage sites M^pro^ (green arrows) and PL^pro^ (red arrows). (C) Schematic representation of the design and working of a genetically encoded, BRET-based SARS-CoV-2 DuProSense biosensor. Close positioning of mNG and RFP with NLuc will result in a significant resonance energy transfer in the absence of the SARS-CoV-2 M^pro^ and PL^pro^ activity. M^pro^-mediated cleavage of the biosensor will result in the separation of NLuc and mNG leading to a decrease in the resonance energy transfer between NLuc and mNG that can be detected as a reduction of the green fluorescence. PL^pro^-mediated cleavage of the biosensor will result in the separation of NLuc and RFP leading to a reduction in the resonance energy transfer between NLuc and RFP that can be detected as a decrease in the red fluorescence. Cleavage of the biosensor by both M^pro^ and PL^pro^ will lead to a reduction in both green and red fluorescence.

Previously, cleavage kinetics of M^pro^ and PL^pro^ have been reported using in vitro assays.^11–14^ These mechanistic studies provided details of enzyme-substrate interactions under controlled conditions. However, the SARS-CoV-2 replication may be influenced by the dynamic and complex cellular environment with several factors affecting M^pro^ and PL^pro^ activity such as cell signaling, regulation, subcellular localization, intermolecular interactions, and dynamical substrate availability that may be caused by the presence of host cell substrates.^15–17^ Therefore, to comprehensively understand the complexities of viral protease kinetics within host cells, we aimed to determine the cleavage kinetics of all cleavage sites present in the SARS-CoV-2 viral polyprotein by M^pro^ and PL^pro^ in live cells.

Previously, several assays have been developed to study these proteases in live cells, including those that utilize Bioluminescence Resonance Energy Transfer (BRET), a phenomenon of proximity-dependant non-radiative, Förster resonance energy transfer between a bioluminescent luciferase donor protein and a fluorescent acceptor protein.^18^ Besides physical proximity, the efficiency of BRET depends on the spectral overlap of the donor emission spectra and acceptor excitation spectra and the relative orientation of the donor and the acceptor moieties.^18^ It is a naturally occurring phenomenon and could be observed, for instance, between the jellyfish, *Aequorea*, photoprotein aequorin, which emits blue light and is transferred to the green fluorescent protein (GFP) leading to the production of green light.^18^ BRET was initially utilized for studying the interaction between cyanobacterial circadian proteins KaiA and KaiB leading to the observation of KaiB homodimerization.^19^ Following this initial report, BRET has been utilized in developing a large variety of biological and biomedical applications.^20–24^ This is, in part, due to several advantages of BRET over the widely used, competing technology, Fluorescence Resonance Energy Transfer (FRET), wherein resonance energy transfer occurs between two fluorescent proteins acting as an energy donor and an energy acceptor.^21^ Some of these advantages include no requirement for an external light source for donor fluorescent protein excitation and low non-specific emission in the acceptor channel that is typically observed in FRET experiments, either due to bleed through from optical filters or direct excitation of the acceptor fluorescent protein.^28^ Some key examples of assays wherein BRET has been utilized successfully include monitoring protein-protein interaction^23^ such as the formation of multiprotein complexes by G-protein coupled receptors (GPCRs)^25^, detecting protein conformational changes^26–29^, engineering biosensors for detecting biomolecules such as cAMP^30^ and cGMP^31,32^, detecting receptor-ligand binding^33,34^, optogenetics using the LOV domain protein^35^, or monitoring the activity of proteolytic enzymes^36–38^.

These successful applications of BRET have been largely due to significant advances in the engineering of brighter and smaller luciferases such as mutants of *Renilla* luciferase^39,40^, NanoLuc (NLuc)^36,41^ and more recently, artificial luciferases such as picALuc^42^ and its mutants^43^, luciferase and their substrates with altered emission spectra^44,45^, spectrally shifted and/or brighter fluorescent acceptor proteins such as GFP^2^ or mNeonGreen (mNG)^22,28,31^. In addition to these, recently a combination of BRET and FRET has been utilized to develop several applications including biosensors such as those for ERK^44^, cAMP^46^, Zn^2+47^, development of optical encryption keys using dendrimer DNA-based nanoscaffold^48^ as well as for visualizing tissue inflammation^49^. While these have undoubtedly increased the applications of BRET, they are limited to the utilization of a single acceptor fluorescent protein and therefore, BRET assays have been limited to reporting a single molecular event at a time.

In this study, we intend to increase the applicability of BRET through the utilization of two, spectrally distinct fluorescent acceptor proteins for simultaneous monitoring of two types of molecular events through determining resonance energy transfer to the acceptor fluorescent proteins in a single biosensor construct. For this, we utilize the previously described BRET donor-acceptor pair consisting of NLuc (luciferase) and mNG (green fluorescent protein) for monitoring BRET in the green channel and screen several red fluorescent proteins as the second acceptor for monitoring BRET in the red channel. In summary, we have designed a BRET-based dual protease biosensor (DuProSense biosensor) by including cognate cleavage sites of M^pro^ and PL^pro^ between mNG and NLuc (green channel) and NLuc and an RFP (red channel), respectively. Based on the results obtained from fluorescence, bioluminescence, and BRET measurements with constructed biosensors using various RFPs, including long Stoke-shifted RFPs (mBeRFP, LSS-mKate2, CyOFP1, and mScarlet), we selected mScarlet as the most suitable second acceptor fluorescent protein. Importantly, we show the specificity and sensitivity of the mNG and mScarlet-based DuProSense biosensor through mutagenesis and known pharmacological inhibitors. The DuProSense biosensor has been used to determine all the viral polyprotein cleavage site kinetics of M^pro^ and PL^pro^ in live cells. Moreover, we characterized all SARS-CoV-2 M^pro^ substrate cleavage sites based on its inhibition by an FDA-approved inhibitor, nirmatrelvir. Thus, the engineering of a BRET-based DuProSense biosensor will be of particular interest because of its capability to enable simultaneous monitoring of the activity of the proteases and therefore, the activity of pharmacological agents against the proteases or mutations in the proteases^50^.

## Results

### BRET-based dual protease biosensor (DuProSense biosensor) design

To simultaneously monitor the cleavage activity of M^pro^ and PL^pro^ in living cells, we decided to utilize BRET as the technology. However, BRET has been typically limited to monitoring a single biomolecular event so far and therefore, cannot be directly utilized for simultaneous monitoring of two distinct molecular events. To increase the capability of BRET-based biosensors from detecting a single molecular event to two, we aimed to develop the DuProSense biosensor to simultaneously detect two distinct molecular events, i.e., proteolytic cleavage, using a single BRET-based biosensor (Fig. 1C). One possibility of achieving this would be to include a second BRET acceptor in the biosensor construct that will report on the second molecular, in addition to the first BRET acceptor that reports the first molecular event. For this, we engineered a fusion protein containing the bright, monomeric mNG, which acted as the BRET acceptor in the green channel, on the N-terminal of NLuc and the bright, monomeric mSca, which acted as the BRET acceptor in the red channel, on the C-terminal of NLuc. The emission peak of 594 nm of mSca^51^ is well separated from the emission peak of mNG (517 nm), and therefore will allow unambiguous spectral separation. The key challenges in designing DuProSense is the possibility of a direct fluorescence resonance energy (FRET) from the green channel BRET acceptor, mNG, to the red channel BRET acceptor and a bleed through of mNG signal into mSca. To determine the impact of these issues and to devise ways to mitigate them, we utilized protein constructs containing NLuc^52^ alone and mNG-NLuc^38^ and designed another with NLuc-mSca and compared their bioluminescence spectra with that of mNG-NLuc-mSca DuProSense biosensor construct (Fig. 2A). Bioluminescence spectral measurements of cells expressing NLuc alone showed a single peak (at 467 nm) while mNG-NLuc and NLuc-mSca showed two peaks (467 and 533 nm and 467 and 615 nm, respectively) (Fig. 2A). On the other hand, the mNG-NLuc-mSca DuProSense biosensor construct showed three peaks (467, 533 and 615 nm) (Fig. 2A).

**Fig. 2.**
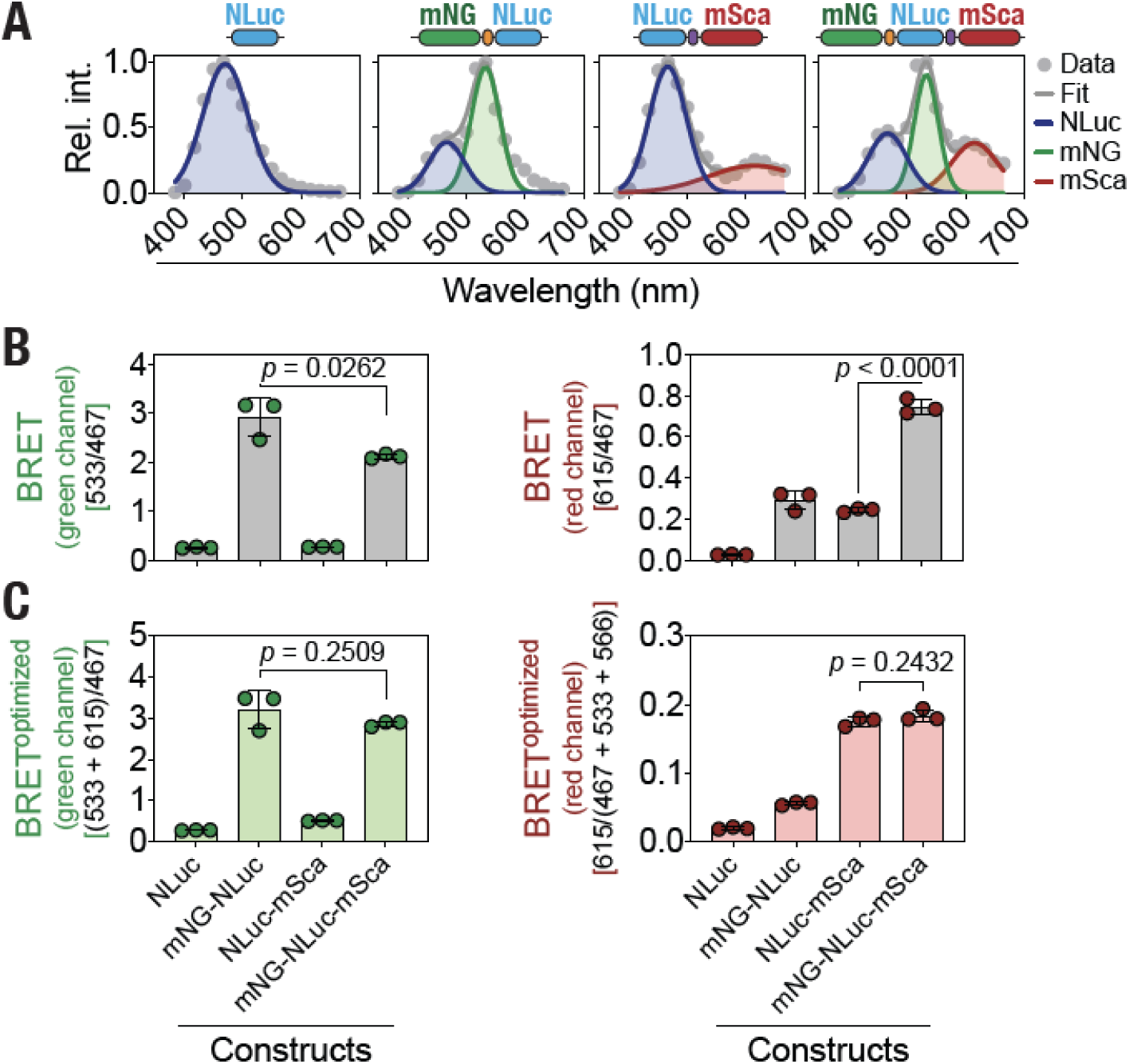
Optimized BRET calculations for a dual color BRET-based biosensor construct. (A) Graphs showing bioluminescence spectra measured from cells expressing NLuc, mNG-NLuc, NLuc-mSca and mNG-NLuc-mSca protein constructs. Spectra were normalized with the peak NLuc emission (*I*_467_) for the NLuc alone and NLuc-mSca constructs and with the peak mNG emission (*I*_533_) for mNG-NLuc and mNG-NLuc-mSca constructs for ease of visualization of various emission peaks. Data were fit to a single Gaussian model for NLuc, a double Gaussian model for mNG-NLuc and NLuc-mSca, and a triple Gaussian model for mNG-NLuc-mSca. (B,C) Graphs showing conventional green (left panel) and red (right panel) channel BRET (B) and optimized green (left panel) and red (right panel) channel BRET (C) values of the indicated protein constructs. Data shown are mean ± S.D. from three independent experiments, with each experiment performed in triplicates.

We then determined BRET in the green and red channels for each of the constructs using conventional method i.e. ratio of peak mNG and NLuc emissions (*I*_533_/*I*_467_) for the green and ratio of peak mSca and NLuc emissions (*I*_615_/*I*_467_) for the red channel. This revealed low green channel BRET values for the NLuc and the NLuc-mSca, constructs that do not contain mNG, and high green channel BRET for the mNG-NLuc and the mNG-NLuc-mSca, constructs that do contain mNG (Fig. 2B; left panel). On the other hand, while red channel BRET values were found to be low for the NLuc alone construct and high for the mNG-NLuc-mSca construct, it was found to be similar for the mNG-NLuc, a construct that do not contain mSca, and the NLuc-mSca, a construct that does contain mSca, (Fig. 2B; right panel), suggestive of a direct FRET between mNG and mSca and/or bleed through from the green to the red channel. We then performed an analysis of the bioluminescence spectra and empirically derived equations optimized for the two channels and determined BRET in the green (ratio of *I*_533+615_ and *I*_467_) and red (ratio of *I*_615_ and *I*_467+533+566_) channels for each of the constructs (see Materials and Methods section). This revealed a very low green channel BRET for NLuc alone and NLuc-mSca constructs but a high green channel BRET for mNG-NLuc and mNG-NLuc-mSca constructs (Fig. 2C; left panel). Similarly, a very low red channel BRET was observed for the NLuc alone and the mNG-NLuc constructs but high red channel BRET for NLuc-mSca and mNG-NLuc-mSca constructs (Fig. 2C; right panel). Importantly, the green channel BRET was similar for the mNG-NLuc and the mNG-NLuc-mSca constructs and the red channel BRET was similar for the NLuc-mSca and the mNG-NLuc-mSca constructs (Fig. 2C; right panel). Thus, we were able to successfully mitigate issues arising due to a direct FRET between mNG and mSca and bleed through of mNG (green) emission into mSca (red) channel using the optimized method for calculating green and red channel BRET values for the DuProSense biosensor construct.

### Spectral characterization of various RFP-containing DuProSense biosensor constructs

Having established a method to unambiguously assign green and red channel BRET in the mSca-containing DuProSense construct, we further explored the possibility of utilizing long Stoke-shifted RFPs to increase the spectral overlap between NLuc and the RFP and thus, increase BRET efficiency in the red channel. For this, we analyzed currently available RFPs in the literature (data available at fpbase.org) and chose to consider mBeRFP, LSS-mKate2, and CyOFP1, in addition to mSca, (Table 1) as the probable second BRET acceptor. First, the long Stoke-shifted RFP, mBeRFP^38^, has been reported to have a brightness of 17.6 but excellent excitation spectral overlap with NLuc emission spectra and largest separation of maximal excitation and emission wavelengths (λ_ex_ and λ_em_ of 446 and 611 nm) (Fig. 3A,B). Second, the long Stoke-shifted RFP, LSS-mKate2^53^, has a lesser brightness (4) but the good excitation spectral overlap with NLuc emission spectra as well as good separation of maximal excitation and emission wavelengths (λ_ex_ and λ_em_ of 460 and 605 nm) (Fig. 3A,B). Third, the long stoke-shifted RFP, CyOFP1^54^, has a relatively higher brightness (30) as well as excellent excitation spectral overlap with NLuc emission spectra (λ_ex_ and λ_em_ of 497 and 589 nm) (Fig. 3A,B). We compared the above mentioned RFPs with mSca^51^, which has the highest fluorescence emission yield (brightness value of 70) but less excitation spectral (λ_ex_ and λ_em_ of 569 and 594 nm) overlap with NLuc emission spectra (Fig. 3A,B).

**Table 1.** Spectral and photophysical properties of selected RFPs used to generate various DuProSense biosensor constructs (based on data available at www.fpbase.org).

| | $\lambda_{\text{ex}}$ (nm) | $\lambda_{\text{em}}$ (nm) | $\text{EC (M}^{-1} \text{ cm}^{-1}) \times 10^3$ | Quantum Yield | Brightness | Maturation (min) |
| --- | --- | --- | --- | --- | --- | --- |
| **mBeRFP** | 446 | 611 | 65 | 0.3 | 18 | 60 |
| **LSS-mKate2** | 460 | 605 | 26 | 0.2 | 4 | 150 |
| **CyOFP1** | 497 | 589 | 40 | 0.8 | 30 | 15 |
| **mSca** | 569 | 594 | 100 | 0.7 | 70 | 174 |

**Fig. 3.**
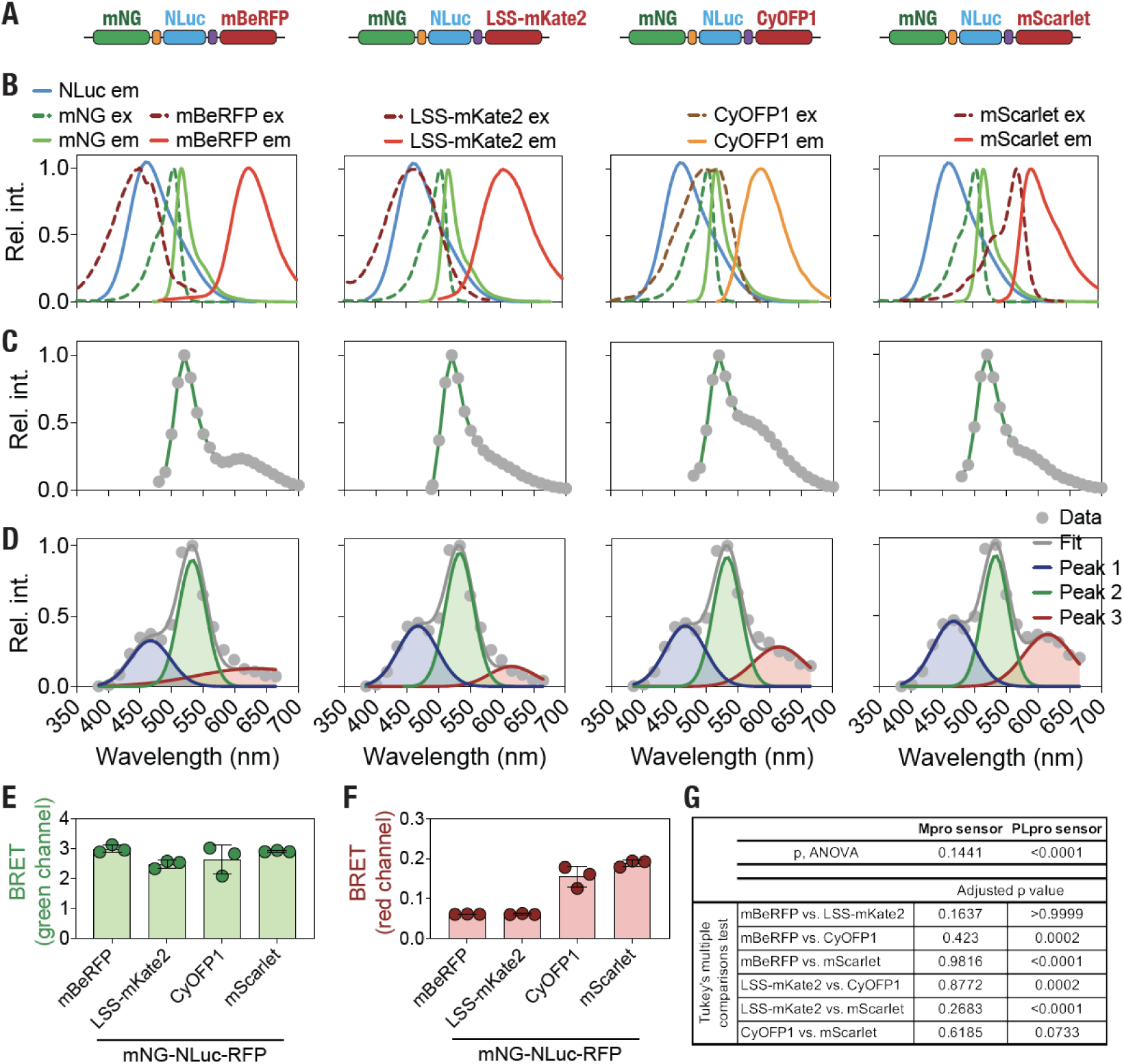
Spectral characterization of different RFP-containing DuProSense biosensor constructs. (A) Schematic showing DuProSense biosensor containing the indicated red fluorescent protein (RFP), in addition to mNG and NLuc. (B) Graph showing NLuc bioluminescence spectra along with the emission and excitation of mNG and indicated RFP. NLuc bioluminescence spectra were experimentally determined while the emission and excitation spectra of the indicated fluorescent proteins were obtained from the FPbase database (www.fpbase.org). (C, D) Graph showing fluorescence (C) and bioluminescence (D) emission spectra of lysates prepared from HEK293T cells expressing DuProSense biosensor constructs containing mBeRFP, LSS-mKate2, CyOFP1 and mSca RFPs. Fluorescence spectra was acquired at an excitation wavelength of 440 nm. Bioluminescence spectra data were fit to a triple Gaussian model. (E,F) Graph showing green (E) and red (F) channel BRET measured from living HEK293T cells expressing DuProSense biosensors constructs containing the indicated RFP after 48 h of transfection. Data shown are mean ± S.D. obtained from three independent experiments, with each experiment performed in triplicates. (G) Table showing results of a one-way ANOVA with Tukey’s multiple comparison test including adjusted *p*-values of the green and red channel BRET obtained from lysates prepared from cells expressing the indicated RFP-containing DuProSense biosensor.

To spectrally characterize mBeRFP, LSS-mKate2, CyOFP1 and mSca containing DuProSense biosensors, we generated plasmid DNA constructs for expressing the biosensors and transfected human embryonic kidney HEK293T cells with the plasmid DNA individually. We then prepared lysates from the transfected cells and measured both fluorescence and bioluminescence spectra (Fig. 3C,D). Fluorescence spectra acquired using 440 nm excitation wavelength showed a peak at ∼533 nm, corresponding to mNG, for all constructs (Fig. 3C). Additionally, a second peak corresponding to the RFPs could also be observed (Fig. 3C), confirming the presence of the RFPs in the DuProSense biosensor constructs. We note that in addition to the possibility of direct excitation of various RFPs, this may also indicate FRET-based energy transfer from mNG to the RFPs. Following this, we measured bioluminescence spectra of each DuProSense biosensor construct after the addition of NLuc substrate. This revealed emission peaks corresponding to NLuc, mNG and the RFP in each of the DuProSense biosensor constructs, although the amplitudes of RFP emission varied between the constructs and were much lower than that of mNG (Fig. 3D). Further, we determined BRET efficiencies in both the green as well as the red channels using the optimized BRET equations described in the previous section (see Materials & Methods for details) (Fig. 3E,F,G). This revealed that BRET efficiencies for NLuc and mNG (green channel biosensor module) were much higher (3.00 ± 0.12, 2.49 ± 0.14, 2.65 ± 0.50 and 2.92 ± 0.03, respectively for mBeRFP, LSS-mKate2, CyOFP1 and mSca-containing DuProSense biosensor constructs; mean ± s.d.; *N* = 3) compared to those between NLuc and RFP (red channel biosensor module) (0.06 ± 0.00, 0.06 ± 0.00, 0.16 ± 0.03 and 0.19 ± 0.01, respectively for mBeRFP, LSS-mKate2, CyOFP1 and mSca-containing DuProSense biosensor constructs; *N* = 3) in each of the DuProSense biosensor constructs. Importantly, while green channel biosensor BRET efficiency remained constant, the red channel biosensor BRET efficiency was found to be high for the DuProSense biosensor constructs containing CyOFP1 and mSca as the RFP acceptor (Fig. 3E,F,G), likely highlighting a role for quantum yield of the RFP (0.8 and 0.7 for CyOFP1 and mSca, respectively, in comparison to 0.3 and 0.2 for mBeRFP and LSS-mKate2, respectively; Pearson *r* = 0.93; Table 1) as the major determinant of BRET efficiency.

### mSca-containing DuProSense biosensor enables simultaneous detection of SARS-CoV-2 M^pro^ and PL^pro^ activity in a highly specific manner

Following spectral characterization of various RFP-containing DuProSense biosensor constructs, we aimed to simultaneously detect the cleavage activity of SARS-CoV-2 M^pro^ and PL^pro^ proteases in live cells using DuProSense biosensor constructs. For this, we included the SARS-CoV-2 M^pro^ N-terminal autocleavage sequence (AVLQSGFR) between mNG and NLuc (green channel biosensor) and PL^pro^ N-terminal autocleavage sequence (LKGGAPTK) between NLuc and RFPs (red channel biosensor) in the DuProSense biosensor constructs. We then transfected HEK293T cells with various RFP-containing DuProSense biosensor constructs along with either M^pro^ or PL^pro^ or both and monitored BRET in both the green (M^pro^ biosensor) as well as the red (PL^pro^ biosensor) channels (Fig. 4). Expression of M^pro^ alone with the DuProSense biosensor constructs resulted in ∼75% decrease in BRET efficiency in the green channel (M^pro^ biosensor module) while expression of PL^pro^ alone did not result in such changes (Fig. 4A,B,C,D; left panel). Importantly, the expression of both M^pro^ and PL^pro^ resulted in similar decreases (∼75%) in the BRET efficiency in the green channel (Fig. 4A,B,C,D; left panel). On the other hand, expression of M^pro^ alone did not result in decreases in BRET efficiency in the red channel (PL^pro^ biosensor module) as did expression of PL^pro^ alone while expression of both M^pro^ and PL^pro^ resulted in a large decrease in the BRET efficiency in the red channel (Fig. 4A,B,C,D; right panel). Interestingly, the decrease in BRET efficiency in the red channel was found to be varying with the LSS-mKate2-containing DuProSense biosensor showed a minimal decrease while the mSca-containing DuProSense biosensor showed the maximal decrease.

**Fig. 4.**
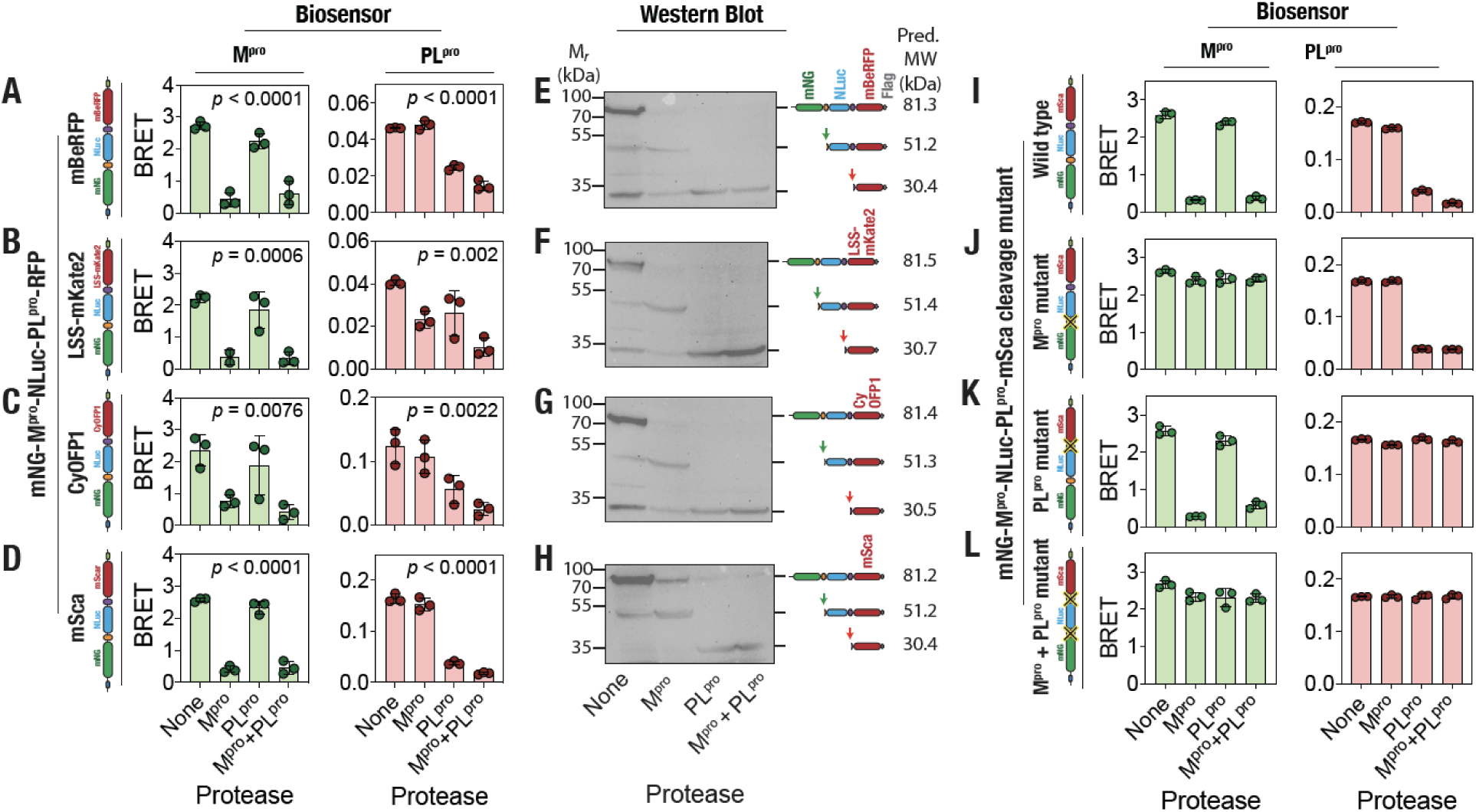
mSca-containing DuProSense biosensor enables robust and specific detection of SARS-CoV-2 M^pro^ and PL^pro^ activity in living cells. (A-D) M^pro^ (green bar graphs; left panels) and PL^pro^ (red bar graphs; right panels) biosensor BRET measured from living HEK293T cells expressing the DuProSense biosensor constructs containing mBeRFP (A), LSS-mKate2 (B), CyOFP1 (C) and mSca (D) in the absence of any protease or in the presence of either M^pro^ or PL^pro^ or both after 48 h of transfection. Note the specific and high decrease in both the M^pro^ (green) and PL^pro^ (red) BRET of the mSca-containing DuProSense biosensor (D). Data shown are mean ± S.D. obtained from three independent experiments, with each experiment performed in triplicates. (E-H) Anti-FLAG-tag western blot images showing proteolytic cleavage of mBeRFP (E), LSS-mKate2 (F), CyOFP1 (G) and mSca (H)-containing DuProSense biosensor constructs in the absence of any protease (first lane) and in the presence of either M^pro^ (second lane) or PL^pro^ (third lane) and both M^pro^ and PL^pro^ (fourth lane). Predicted molecular weights of the DuProSense biosensor constructs and proteolytic cleavage products are indicated on the right side. (I-L) M^pro^ (green bar graphs; left panel) and PL^pro^ (red bar graphs; right panels) biosensor BRET measured from living HEK293T cells expressing the mSca-containing DuProSense biosensor containing either both M^pro^ and PL^pro^ cleavage sites (I), or only PL^pro^ cleavage site (M^pro^ mutant) (J), or only M^pro^ cleavage site (PL^pro^ mutant) (K), or neither M^pro^ nor PL^pro^ cleavage sites (both M^pro^ + PL^pro^ mutant) (L) in the absence of any protease or in the presence of either M^pro^ or PL^pro^ or both in living HEK293T cells after 48 h of transfection. Data shown are mean ± S.D. obtained from three independent experiments, with each experiment performed in triplicates.

In order to confirm if these decreases in BRET efficiencies of the DuProSense biosensor constructs in the presence of either M^pro^ or PL^pro^ or both were due to cleavage of the biosensor proteins, we performed western blot analysis using an anti-FLAG-tag antibody. Each of the four DuProSense biosensor constructs contained a C-terminal 3×FLAG-tag following RFP sequences and therefore, either M^pro^ or PL^pro^-mediated cleavage of the biosensor constructs will result in the appearance of bands with smaller molecular weights. Indeed, expression of M^pro^ in cells along with the four DuProSense biosensor constructs resulted in the loss of the ∼81 kDa band corresponding to the intact DuProSense biosensor construct with a concomitant appearance of a ∼51 kDa band corresponding to the NLuc-RFP fragment (Fig. 4E,F,G,H). Similarly, expression of PL^pro^ resulted in the loss of the band corresponding to the intact DuProSense biosensor constructs with a concomitant appearance of an ∼30 kDa band corresponding to respective RFPs (Fig. 4E,F,G,H). Expression of both M^pro^ and PL^pro^ also resulted in a loss of the band corresponding to the intact DuProSense biosensor construct with the concomitant appearance of a ∼30 kDa band corresponding to respective RFPs. While these results unambiguously revealed a proteolytic cleavage of the four DuProSense biosensor constructs, we noticed significant non-specific cleavage of the mBeRFP, LSS-mKate2, and CyOFP1-containing DuProSense biosensor constructs in the absence of the two SARS-CoV-2 proteases as evidenced by the appearance of a ∼30 kDa band in the blots (Fig. 4E,F,G) while such a band was not observed with the mSca-containing DuProSense biosensor construct. Thus, based on the data presented above i.e., BRET ratio in the green and red channels (high BRET in the red channel), the percentage decrease in BRET efficiency in the presence of protease (maximal decrease in the red channel BRET efficiency in the presence of PL^pro^) and western blot analysis (no non-specific cleavage), we decided to utilize the mSca-containing DuProSense biosensor for further studies.

Following the establishment of mSca-containing DuProSense biosensor as the BRET-based dual protease biosensor of choice, we attempted to determine the specificity of SARS-CoV-2 M^pro^ and PL^pro^-mediated cleavage of the biosensor^5–8,55–57^. For this, we generated mutations in either the M^pro^ cleavage site or the PL^pro^ cleavage site or both through the mutation of the critical Gln (fourth residue in the M^pro^ cleavage site) to Ala (from AVL**Q**SGFR to AVL**A**SGFR) and Gly (fourth residue in the PL^pro^ cleavage site) to Ile (from LKG**G**APTK to LKG**I**APTK). We then expressed the wild type and mutant mSca-containing DuProSense biosensors along with either M^pro^ or PL^pro^ or both proteases. While expression of M^pro^ resulted in a large decrease in the green channel (M^pro^ biosensor module) BRET efficiency of the wild type (Fig. 4I; left panel) as well as the PL^pro^ cleavage site mutant (Fig. 4K; left panel) DuProSense biosensors, such decreases were not seen with M^pro^ cleavage site mutant (M^pro^ as well as both M^pro^ and PL^pro^ cleavage site mutant DuProSense biosensors) (Fig. 4J,L; left panel). Similarly, while expression of PL^pro^ resulted in a large decrease in the red channel (PL^pro^ biosensor module) BRET efficiency of the wild type (Fig. 4I; right panel) as well as the M^pro^ cleavage site mutant (Fig. 4J; right panel) DuProSense biosensors, such decreases were not seen with the PL^pro^ cleavage site mutant (PL^pro^ as well as both M^pro^ and PL^pro^ cleavage site mutant DuProSense biosensors) (Fig. 4K,L; right panel). Importantly, expression of both M^pro^ and PL^pro^ in the cells specifically resulted in a decrease in BRET efficiency of wild type and PL^pro^ cleavage site mutant DuProSense biosensor in the green channel (M^pro^ biosensor module) and wild type and M^pro^ cleavage site mutant DuProSense biosensor in the red channel (PL^pro^ biosensor module). On the other hand, mutation of either the M^pro^, PL^pro^ or both M^pro^ and PL^pro^ cleavage sites resulted in a loss in BRET decreases in the presence of M^pro^ or PL^pro^ and both M^pro^ and PL^pro^ protease, respectively (Fig. 4J-L; Supporting Figure 1). Overall, these results indicate that the response of DuProSense biosensor in the form of a decrease in BRET efficiency in the presence of either M^pro^ or PL^pro^ or both proteases is highly specific.

After determining the specificity of the mSca-containing DuProSense biosensor, we determined the M^pro^ and PL^pro^ plasmid DNA concentration-dependent cleavage of DuProSense biosensor. For this, we transfected the cells with increasing amounts of either M^pro^ or PL^pro^ protease expressing plasmid DNA, along with a constant amount of the DuProSense biosensor plasmid DNA and monitored green and red channel BRET. This revealed an M^pro^ and PL^pro^ plasmid DNA amount-dependent decrease in green and red channel BRET with *EC*_50_ values of 3.8 ± 1.5 and 2.3 ± 1.9 ng/well, respectively (Supporting Figure 2). Moreover, we monitored the cleavage of DuProSense biosensor upon transfection of increasing amounts of both M^pro^ and PL^pro^ plasmid DNA simultaneously. This resulted in a decrease in BRET of both the green (M^pro^ biosensor module) and the red (PL^pro^ biosensor module) simultaneously with *EC*_50_ values of 4.5 ± 1.8 and 1.5 ± 0.6 ng/well, respectively (Supporting Figure 2), values similar to those observed upon expression of M^pro^ and PL^pro^ individually.

Having established the specificity of the DuProSense biosensor, we then monitored the inhibition of SARS-CoV-2 M^pro^ and PL^pro^ proteases in live cells using the mSca-containing DuProSense biosensor. For this, we transfected cells with the mSca-containing DuProSense biosensor along with either M^pro^ or PL^pro^ proteases, treated the cells with a range of concentrations of either GC376 (M^pro^ inhibitor)^58^ or GRL-0617 (PL^pro^ inhibitor)^59^ and measured BRET in both the green channel (M^pro^ biosensor module) as well as the red channel (PL^pro^ biosensor module) (Fig. 5). This revealed a GC376 concentration-dependent increase in the green channel BRET efficiency in cells expressing M^pro^ protease, indicating a concentration-dependent inhibition of the M^pro^ protease (Fig. 5A) whereas red channel BRET efficiency in the same cells remained high (Fig. 5C; Supporting Figure 3). Similarly, while the green channel BRET remained high (Fig. 5B), a GRL-0617 concentration-dependent increase in the red channel BRET efficiency in cells expressing PL^pro^ protease was observed, indicating a concentration-dependent inhibition of PL^pro^ protease in the cells (Fig. 5D; Supporting Figure 3). Further, this revealed an IC_50_ value of 13.9 ± 2.9 μM of GC376 against M^pro^ and 77.2 ± 34.8 μM of GRL-0617 against PL^pro^ (Fig. 5E,F), values similar to those reported previously.^59–65^ Thus, the high specificity of signal changes observed with DuProSense biosensor allowed us to unambiguously monitor the pharmacological inhibition of the two SARS-CoV-2 proteases, M^pro^ and PL^pro^.

**Fig. 5.**
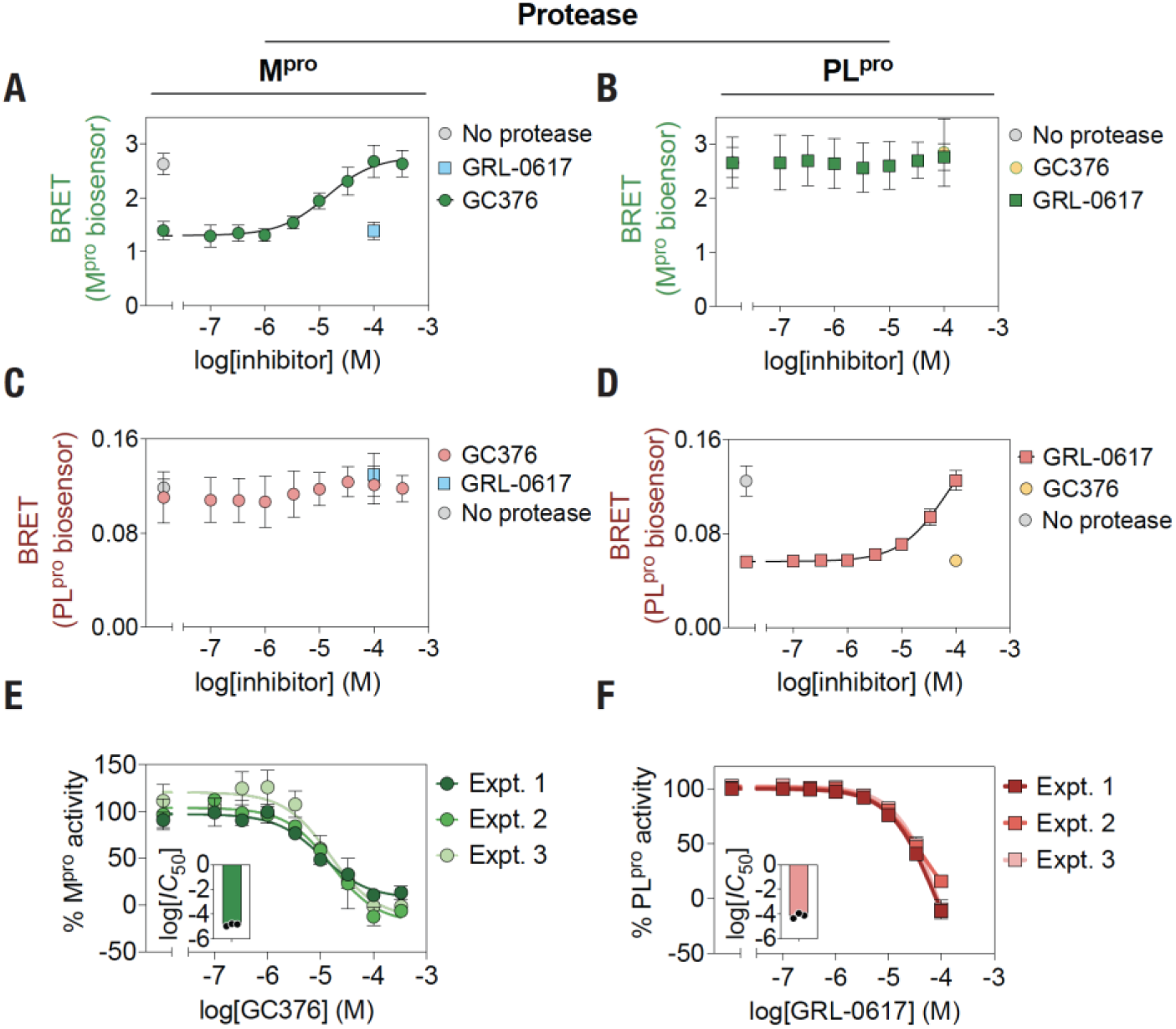
mSca-containing DuProSense biosensor enables highly specific pharmacological inhibition of M^pro^ and PL^pro^ in living cells. (A-D) M^pro^ (A, B) and PL^pro^ (C, D) biosensor BRET measured from living HEK293T cells expressing DuProSense biosensor constructs and M^pro^ or PL^pro^ in the presence of the indicated concentrations of GC376 (A, C) or GRL-0617 (B, D) after 22 h of transfection. (E, F) Graphs showing percentage M^pro^ (E) and PL^pro^ (F) activity in living HEK293T cells expressing DuProSense biosensor constructs and proteases (M^pro^ or PL^pro^) treated with the indicated concentrations of either GC376 (E) or GRL-0617 (F). Insets, graphs showing IC_50_ of GC376 (E) and GRL-0617 (F). Data shown are mean ± S.D. from three independent experiments, with each experiment performed in triplicates.

### Simultaneous monitoring of M^pro^ and PL^pro^ activity temporal dynamics in live cells

Following the utility of the mSca-containing DuProSense biosensor in the simultaneous detection of SARS-CoV-2 M^pro^ and PL^pro^ activity in a highly specific manner, we decided to monitor the temporal dynamics of both M^pro^ and PL^pro^ proteolytic activity in live cells using the biosensor. For this, we transfected HEK293T cells with the biosensor plasmid along with either M^pro^ or PL^pro^ or both protease plasmids and monitored the bioluminescence spectra for 48 h post-transfection (Fig. 6A). Bioluminescence analysis revealed a time-dependent increase in the expression of the biosensor in the initial stages after transfection with values reaching a plateau at later stages in all four conditions (Fig. 6B; Supporting Figure 4). Analysis of bioluminescence spectra of the cells expressing the biosensor in the absence of the protease revealed a time-dependent increase in BRET in the green channel (M^pro^ biosensor module) with BRET values reaching a plateau after about 20 h post-transfection (Fig. 6C; top, left panel), likely suggesting a slower maturation of mNG (BRET acceptor) as compared to NLuc (BRET donor). Similar increases were observed for the red (PL^pro^ biosensor module) channel also, although the rate of increase in red channel BRET was much slower than the green channel BRET (Fig. 6C; bottom, left panel), likely suggesting an even slower maturation of mSca (red channel BRET acceptor), as compared to both mNG and NLuc. Importantly, such increases in the green channel BRET were not observed in cells expressing M^pro^, except for a small increase in the initial period, while the red channel BRET was found to exhibit a similar rate of increase in the same cells (Fig. 6C. On the other hand, while the green channel BRET increased in a manner similar to that of no protease control, the red channel BRET was not found to increase in cells expressing the PL^pro^ protease (Fig. 6C). Cells expressing both M^pro^ and PL^pro^ did not show an increase in either the green, except for a small increase in the initial period, or the red channel BRET (Fig. 6C). The small increase in the green channel BRET likely reflected a slower, compared to the mNG maturation, expression of the functionally active M^pro^, which is known to form a dimer that is required for its proteolytic activity.^66,67^ Conversely, the absence of such increases in the red channel BRET in the case of PL^pro^ expressing cells likely reflected a relatively faster, compared to the slower mSca maturation, expression of PL^pro^ that is functionally active.

**Fig. 6.**
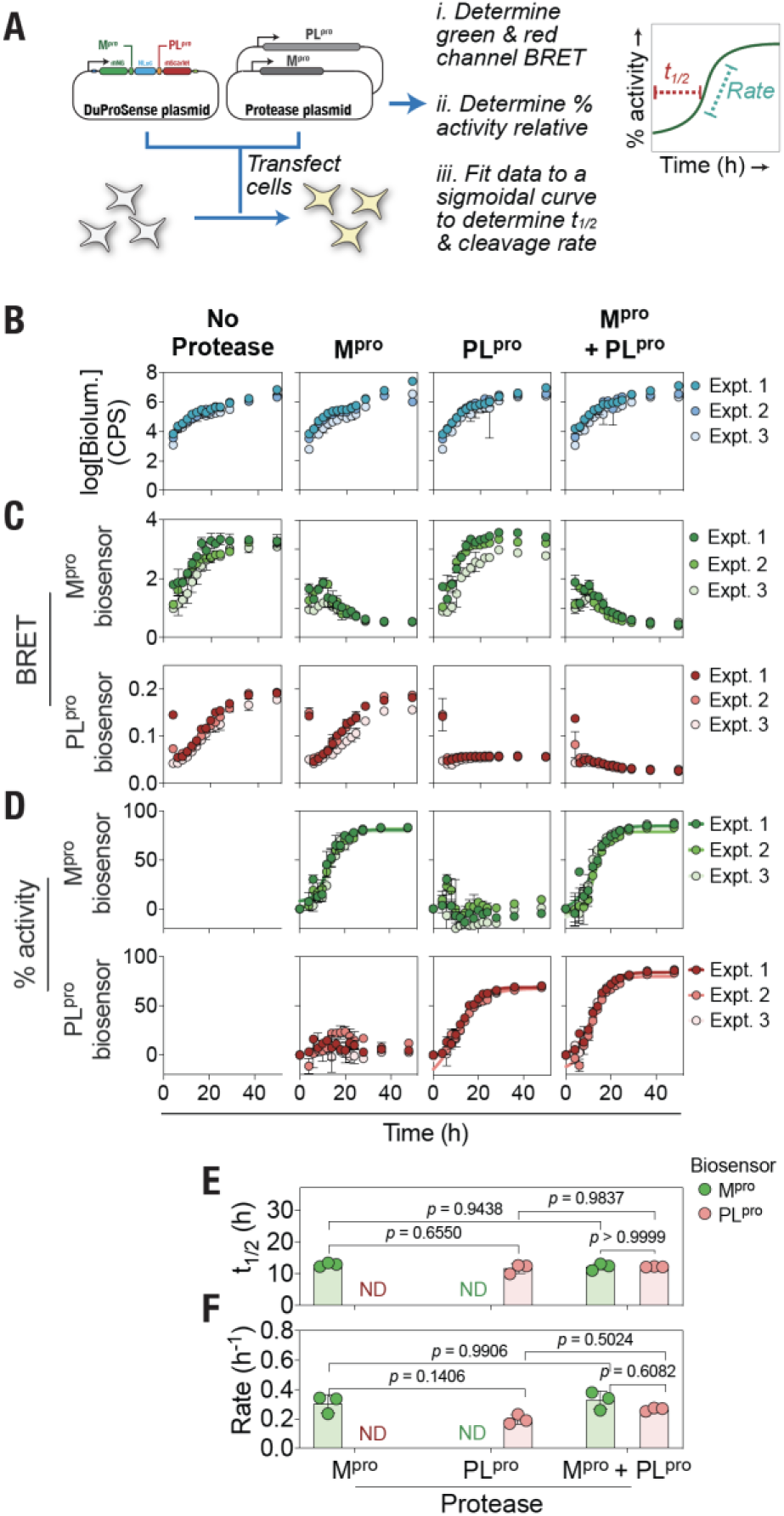
mSca-containing DuProSense biosensor enables simultaneous monitoring of M^pro^ and PL^pro^ cleavage activity in living cells. (A) Schematic showing the experimental protocol in which living HEK293T cells are co-transfected with the DuProSense biosensor and the proteases, followed by monitoring the total bioluminescence, green and red channel BRET over time and data analysis. (B) Graph showing the total bioluminescence measured from living HEK293T cells expressing the mSca-containing DuProSense biosensor in the absence and presence of either M^pro^ and PL^pro^ or both over time. (C) Graph showing the temporal changes in the M^pro^ (top panel) and PL^pro^ biosensor BRET (bottom panel) measured in living HEK293T cells expressing the DuProSense biosensor in the absence or in the presence of the indicated proteases. (D) Graph showing the percentage M^pro^ activity measured by M^pro^ biosensor (green circles) and that of PL^pro^ measured by PL^pro^ biosensor (red circles) in the absence or presence of the proteases. (E, F) Graphs showing the half-time (t_1/2_) (E) and cleavage rate (F) of the M^pro^ and PL^pro^ activity measured from the living HEK293T cells expressing the mSca-containing DuProSense biosensor. Data shown are mean ± S.D. from three independent experiments, with each experiment performed in triplicates.

We utilized the green and red channel BRET values obtained from cells not expressing the proteases and those that express either M^pro^ or PL^pro^ or both to determine the percentage cleavage activity of M^pro^ and PL^pro^ in the protease-expressing cells (Fig. 6D). This analysis revealed a kinetic sigmoidal trend in the cleavage of the M^pro^ biosensor module specifically (maximum activity of 83 ± 0.4%) in the M^pro^ expressing cells and PL^pro^ biosensor module specifically (maximum activity of 70 ± 0.9%) in the PL^pro^ expressing cells. Cells expressing both M^pro^ and PL^pro^ showed a kinetic sigmoidal trend in the cleavage of both M^pro^ and PL^pro^ biosensor modules (maximum activities of 86 ± 2.2 and 86 ± 1.6%; *p* = 0.8728), with the PL^pro^ module showing a higher maximum cleavage activity in cells expressing both M^pro^ and PL^pro^ in comparison to cells expressing only PL^pro^ (86 ± 1.6 vs. 70 ± 0.9%; *p* = 0.0002). We then fitted the data to a kinetic sigmoidal model (see Materials and Methods section for details)to determine the half-life (t_1/2_) and the rate of cleavage of the M^pro^ and PL^pro^ biosensor modules in cells expressing either M^pro^ or PL^pro^ or both (Fig. 6E,F). This revealed that the M^pro^ biosensor module showed a t_1/2_ of 12.9 ± 0.6 and 12.2 ± 1.1 h (*p* = 0.9438) in the presence of only M^pro^ or both M^pro^ and PL^pro^ proteases, respectively, whereas PL^pro^ biosensor module showed a t_1/2_ of 11.7 ± 1.5 and 12.2 ± 0.1 h (*p* = 0.9837) in the presence of only PL^pro^ or both M^pro^ and PL^pro^ proteases, respectively (Fig. 6E). In addition, the kinetic sigmoidal model showed that there was no significant difference in the rate of cleavage of the M^pro^ biosensor module in the presence of only M^pro^ or both proteases (0.31 ± 0.06 vs. 0.33 ± 0.06 h^-1^; *p* = 0.9906) as well as for the PL^pro^ biosensor module in the presence of only PL^pro^ or both proteases (0.19 ± 0.03 vs. 0.27 ± 0.01 h^-1^; *p* = 0.5024) (Fig. 6F). Overall, these results indicate similar cleavage kinetics of both M^pro^ and PL^pro^ biosensor modules by their respective proteases, either individually or together.

### Temporal cleavage dynamics of all pp1a and pp1ab M^pro^ and PL^pro^ cleavage sites in live cells

After establishing the utility of the DuProSense biosensor in simultaneously determining the proteolytic cleavage dynamics of M^pro^ and PL^pro^ in living cells, we aimed to utilize the DuProSense biosensor platform to understand the cleavage kinetics of all SARS-CoV-2 M^pro^ and PL^pro^ cleavage sites in live cells. SARS-CoV-2 pp1a and pp1ab polyproteins contain a total of 11 and 3 cleavage sites for M^pro^ and PL^pro^, respectively (Fig. 7A). While they all been reported to be proteolytically processed by M^pro^ and PL^pro10–14,68^, there are differences in the amino acid sequences of the cleavage site peptides and several of these peptides have been shown to interact differentially with the proteases^16^, thus raising the possibility of a differential cleavage kinetics of the different M^pro^ and PL^pro^ cleavage sites. For this, we designed DuProSense biosensor constructs to monitor the cleavage of each SARS-CoV-2 protease cleavage site compared with the N-terminal auto-cleavage site of their corresponding protease (NSP4-5 for M^pro^ and NSP2-3 for PL^pro^) simultaneously in living cells^14^. Since the N-terminal auto-cleavage sites of both M^pro^ and PL^pro^ have been well characterized^38,69^, we used them as internal controls in the design of each M^pro^ and PL^pro^ cleavage site containing DuProSense biosensors. Specifically, we designed a total of 11 DuProSense biosensor constructs with each containing the M^pro^ N-terminal auto-cleavage site (NSP4-5) between the mNG and NLuc and each of the other cleavage sites of M^pro^ (from NSP4-5 to NSP15-16) between NLuc and mSca. Similarly, we designed two DuProSense biosensor constructs with each containing the PL^pro^ N-terminal auto-cleavage site (NSP2-3) between NLuc and mSca and either NSP1-2 or NSP3-4 cleavage site between the mNG and NLuc (Fig. 7B).

**Fig. 7.**
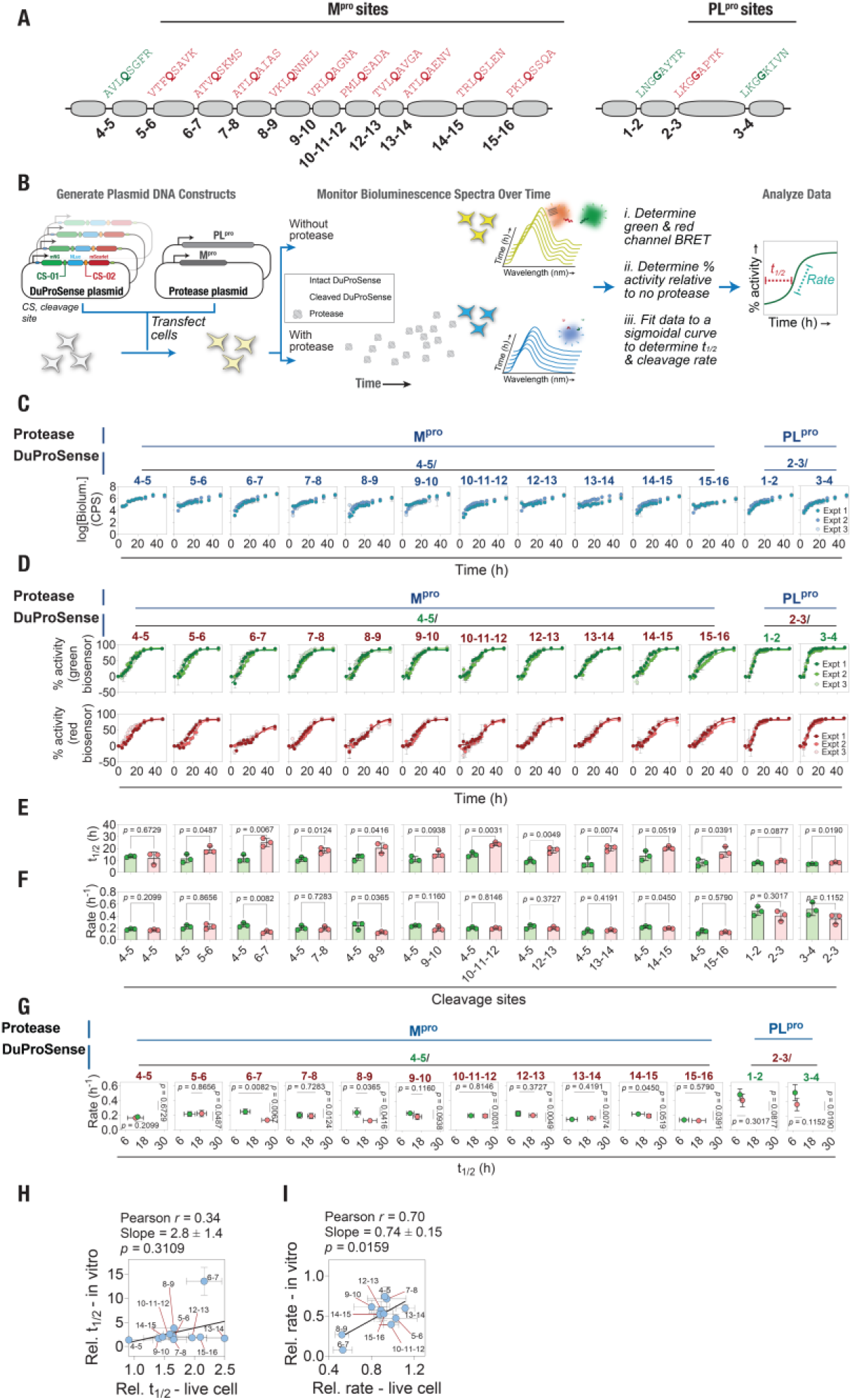
DuProSense biosensor platform reveals differential cleavage kinetics of M^pro^ cleavage sites in living cells. (A) Schematic of viral polyprotein containing M^pro^ and PL^pro^ substrate cleavage sites. (B) Schematic showing the experimental protocol in which living HEK293T cells are co-transfected with the DuProSense biosensor containing M^pro^ and PL^pro^ cleavage sites and the proteases, followed by monitoring the total bioluminescence, green and red channel BRET over time and data analysis. (C) Graph showing the total bioluminescence of living HEK293T cells expressing the DuProSense biosensors containing M^pro^ and PL^pro^ cleavage sites measured at indicated time points after transfection. (D) Graphs showing the percentage protease activity of M^pro^ and PL^pro^ in cells expressing DuProSense biosensors containing M^pro^ and PL^pro^ cleavage sites measured at the indicated time points after transfection. The percentage protease activity from green channel BRET of DuProSense biosensors containing M^pro^ cleavage sites corresponds to NSP4-5 while red channel BRET corresponds to all other M^pro^ cleavage sites. The percentage protease activity from green channel BRET of DuProSense biosensors containing PL^pro^ cleavage sites corresponds to NSP1-2 and NSP3-4 while red channel BRET corresponds to NSP2-3, PL^pro^ N-terminal autocleavage site. (E,F) Graphs showing the half-time (t_1/2_) (E) and rate (F) of cleavage of all substrate sites of M^pro^ and PL^pro^compared to the N-terminal auto-cleavage sites of respective proteases. (G) Graphs comparing the cleavage rate with t_1/2_ of each cleavage site of M^pro^ and PL^pro^. (H,I) Graph showing the correlation of relative t_1/2_ (H) obtained from live cell assay and in vitro assay, and correlation of cleavage rate (I) obtained from live cell assay and in vitro assay. Data shown are mean ± S.D. from three independent experiments, with each experiment performed in triplicates.

To monitor the cleavage kinetics of each M^pro^ and PL^pro^ cleavage site, we transfected HEK293T cells with individual DuProSense biosensor plasmid along with either M^pro^ or PL^pro^ protease expression plasmid and monitored their bioluminescence spectra over time. Expression of the biosensor constructs was confirmed by western blot analysis using the anti-FLAG antibody and lysates prepared from HEK293T cells transfected with the biosensor constructs (Supporting Figure 5). Prior to live cell experiments, we performed FRET measurements with all DuProSense biosensor constructs to determine if the inclusion of different cleavage site peptides altered the conformation of the biosensor constructs. An equal volume of cell lysates prepared from cells expressing the DuProSense biosensor constructs were used to measure FRET. The FRET (measured as a ratio of 590 to 520 nm) between mNG and mSca in each DuProSense biosensor construct showed similar values for the DuProSense biosensor containing PL^pro^ cleavage sites while there were differences in FRET values between DuProSense biosensor containing M^pro^ cleavage sites with NSP15-16 exhibiting the highest FRET value (Supporting Figure 6). These differences in FRET may indicate conformational variations amongst the DuProSense biosensor constructs. We then monitored bioluminescence from cells expressing various DuProSense biosensor constructs either in the absence or in the presence of proteases over time (Supporting Figure 7). A time-dependent increase in total bioluminescence (Fig. 7C) as well as BRET in the green and red biosensor modules (Supporting Figure 8) in the cells expressing various DuProSense biosensor constructs in the absence of the proteases revealed the expression kinetics of the DuProSense biosensor constructs and maturation of the fluorescent proteins, as seen with the M^pro^ and PL^pro^ DuProSense biosensor (Fig. 6). Importantly, the green and red channel BRET values, as determined using the optimized BRET equations developed here, after 48 h of transfection showed no significant difference amongst the DuProSense biosensors (Supporting Figure 9), suggesting that any differences arising from differences in the conformation or FRET in the biosensor constructs or spectral bleed through from the green to the red channels have been addressed highlighting the robustness of the technological platform developed here.

**Fig. 8.**
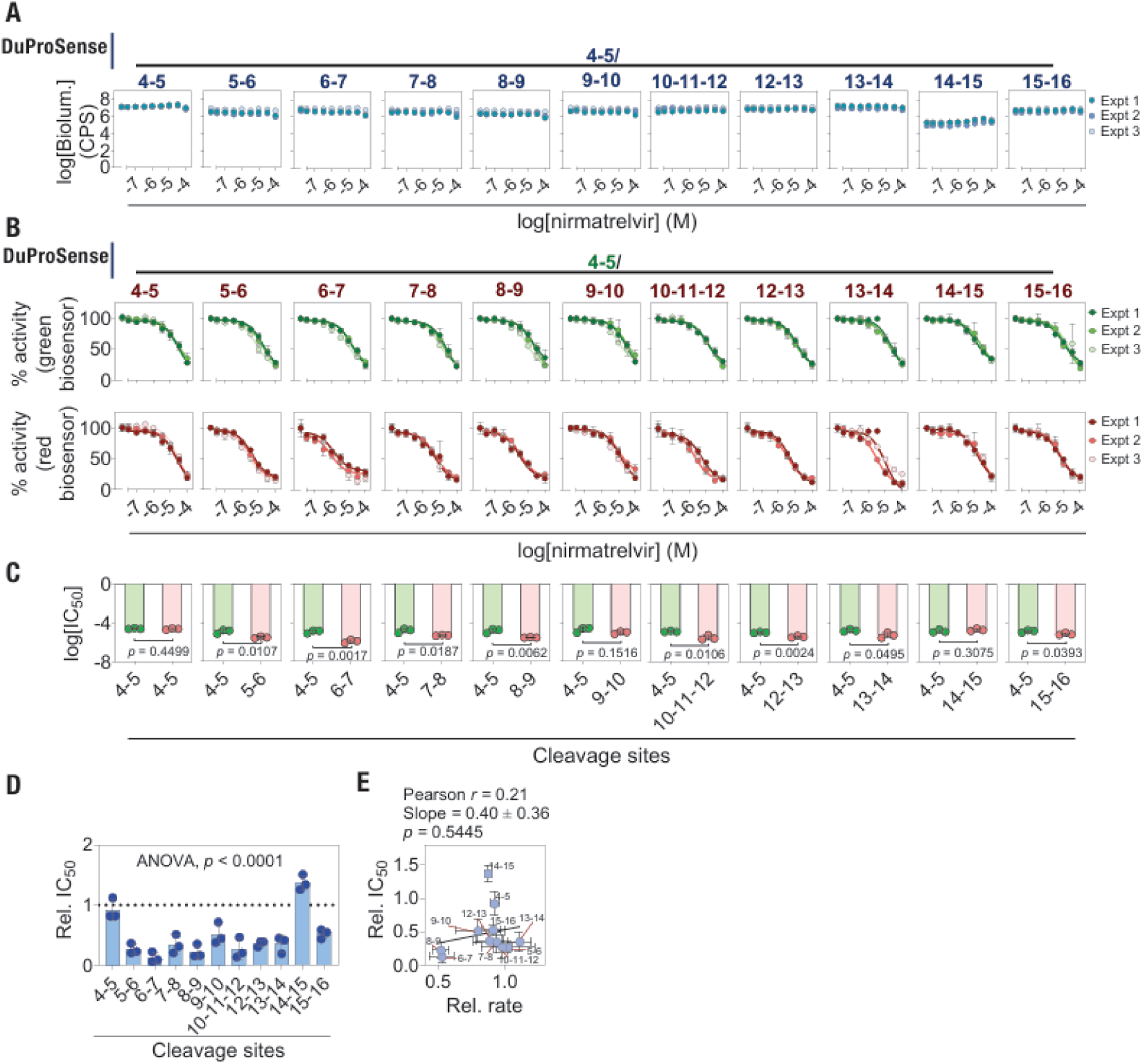
DuProSense biosensor platform reveals cleavage site-specific M^pro^ inhibitory potency of nirmatrelvir in living cells. (A) Graph showing total bioluminescence of cells expressing the DuProSense biosensors containing various M^pro^ cleavage sites in the presence of the indicated concentrations of nirmatrelvir measured after 22 h of transfection. (B) Graphs showing the dose-dependent percentage inhibition of cleavage of the N-terminal auto-cleavage site (NSP4-5) of M^pro^ (green channel; upper panel) and that of other M^pro^ substrate sites (red channel; lower panel). (C) Graph showing the comparison of IC_50_ values of nirmatrelvir obtained from the green and red channels of each DuProSense biosensor construct in expressed in living cells. (D) Graph showing relative IC_50_ obtained for the indicated M^pro^ cleavage sites and the NSP4-5 site obtained from live cell dose-response curve experiments with nirmatrelvir. (E) Graph showing the correlation between relative rate and relative IC_50_ in the presence of nirmatrelvir for corresponding cleavage sites. Data shown are mean ± S.D. from three independent experiments, with each experiment performed in triplicates.

**Fig. 9.**
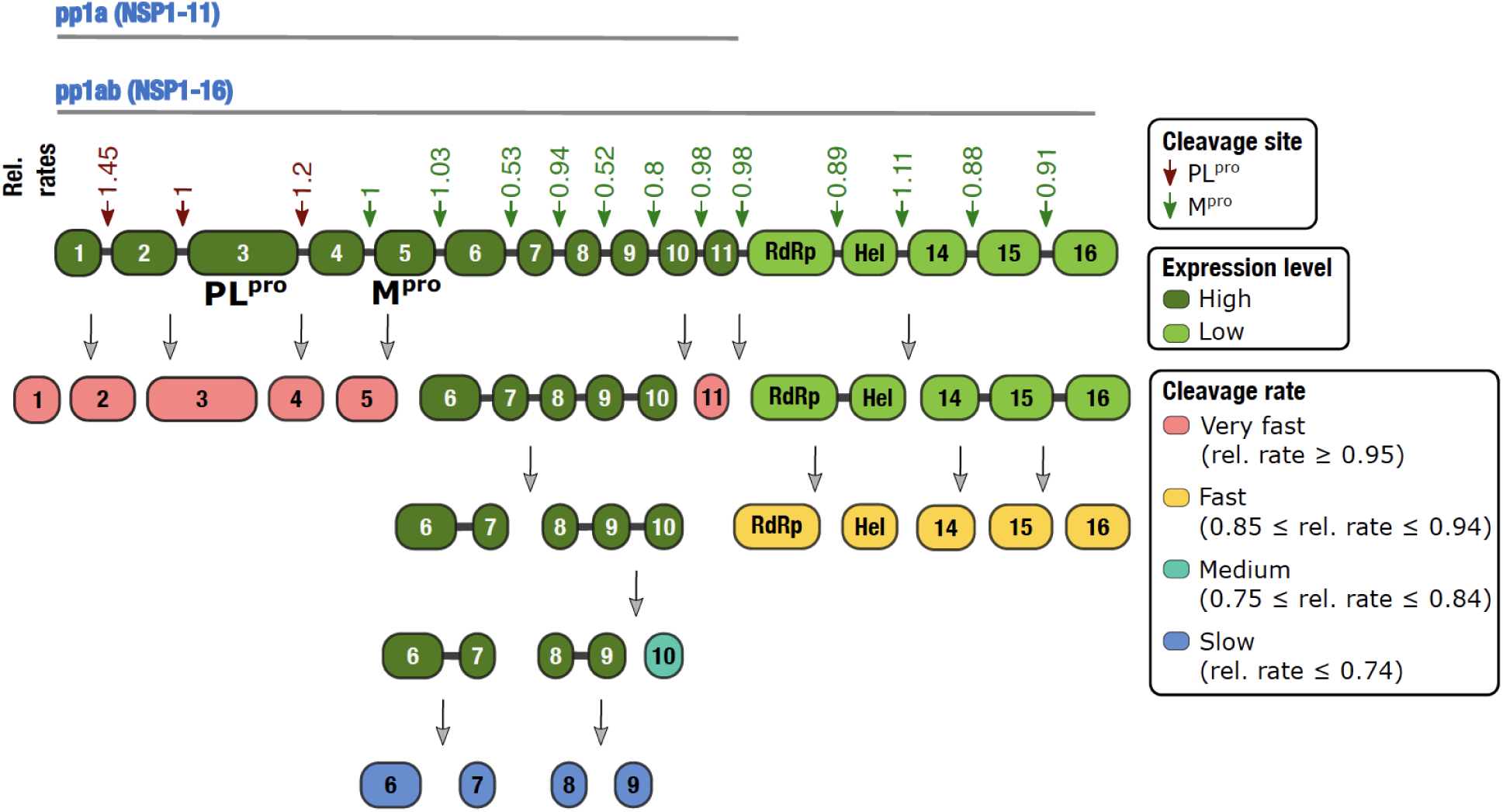
SARS-CoV-2 pp1a and pp1ab polyprotein cleavage kinetics. Schematic showing SARS-CoV-2 polyprotein organization, with PL^pro^ and M^pro^ cleavage sites indicated with red and cyan arrows, respectively, along with their relative rates of cleavage (PL^pro^ cleavage sites in red, relative to NSP2-3; M^pro^ cleavage sites in green, relative to NSP4-5). Proteins from the pp1a are colored dark green, indicating a higher expression level, while those from pp1ab are colored light green, indicating a lower expression level. Cleavage sites are classified as very fast, fast, medium and slow based on their relative rates of proteolytic processing and release of individual proteins is highlighted using a color change based on their cleavage rate classification. Thus, NSP1, NSP2, NSP3, NSP4, NSP5, and NSP11 are released from the polyprotein. These are followed by NSP12 (RdRp), NSP13 (helicase), NSP14, NSP15, and NSP16 are released from their intermediate polyprotein. Subsequently, NSP10 will released from the intermediate polyprotein containing NSP8, 9 and 10. Finally, NSP6, NSP7, NSP8 and NSP9 will be release from the intermediate polyproteins containing NSP6 and 7 and NSP8 and 9, respectively.

Importantly, expression of M^pro^ and PL^pro^ in the cells resulted in a time dependent decrease in the green and the red channel BRET values of DuProSense biosensors containing their respective cleavage sites (Supporting Figure 7). Next, utilizing the BRET values of the green and red channels in the absence and presence of the proteases, we determined the time-dependent activity of the two proteases (percentage) for each of their cleavage sites individually (Fig. 7D; Supporting Figure 7). We utilized the DuProSense biosensor containing NSP4-5 cleavage site in both the green as well as the red channel biosensor modules (i.e. between mNG and NLuc, and NLuc and mSca) to determine whether the position of the cleavage site in the DuProSense biosensor construct or the difference in the maturation of mNG and mSca have any impact on the monitoring of cleavage kinetics of the sites using the DuProSense biosensors. In accordance with the previous observation, the cleavage of both the green and the red biosensor modules occurred in a kinetic sigmoidal fashion in the presence of M^pro^ with a maximum activity of 88 ± 0.5% and 84 ± 0.9% for green and red biosensor modules, respectively (Fig. 7D). Also, fitting the data to a kinetic sigmoidal model revealed no significant difference in the t_1/2_ (13.3 ± 0.6 and 12.0 ± 5.0 h; *p* = 0.6729) and rate of cleavage (0.18 ± 0.01 and 0.16 ± 0.01 h^-1^; *p* = 0.2099) of green and red biosensor modules in the DuProSense biosensor containing NSP4-5 cleavage sites (Fig. 7E,F). Together, these results confirmed that neither the position nor the fluorescent protein has any apparent impact on the kinetics of the protease cleavage sites.

We then analyzed the cleavage kinetics of each DuProSense biosensor containing either M^pro^ or PL^pro^ cleavage sites. This analysis revealed a similar, ∼90% in the green and ∼80% in the red channel, maximum cleavage activity of M^pro^ with various M^pro^ site DuProSense biosensors, except for the biosensor containing NSP4-5/6-7 sites, which showed a significantly reduced extent of cleavage (87 ± 1.9 vs. 58 ± 3.2%; *p* = 0.0002) (Fig. 7D; Supporting Table 2). Similar maximum PL^pro^ cleavage activities were observed for its cleavage sites (∼88% in the green and ∼87% in the red channel) (Fig. 7D; Supporting Table 2). We then fit the percentage activity data to a kinetic sigmoidal model, as done in the previous section (Fig. 6D), to determine the t_1/2_ and rate of cleavage for each of the M^pro^ and PL^pro^ cleavage sites. This analysis revealed similar t_1/2_ (13.3 ± 0.6, 11.6 ± 3.5, 11.7 ± 3, 11.1 ± 1.7, 12.4 ± 2.3, 11.1 ± 2.3, 15 ± 1.9, 9.6 ± 1.6, 8.2 ± 3.4, 14 ± 3.9, and 8.2 ± 2.9 h for NSP4-5, NSP5-6, NSP6-7, NSP7-8, NSP9-10, NSP10-11-12, NSP12-13, NSP13-14, NSP14-15, and NSP15-16, respectively; one-way ANOVA *p* = 0.0700) for the NSP4-5 (green channel) across all M^pro^ site containing DuProSense biosensor constructs (Fig. 7E; Supporting Table 2; Supporting Figure 10B). However, the cleavage rates (0.18 ± 0.01, 0.22 ± 0.03, 0.25 ± 0.03, 0.2 ± 0.04, 0.23 ± 0.06, 0.24 ± 0.01, 0.19 ± 0.02, 0.22 ± 0.04, 0.14 ± 0.03, 0.22 ± 0.01, and 0.14 ± 0.03 h^-1^ for NSP4-5, NSP5-6, NSP6-7, NSP7-8, NSP9-10, NSP10-11-12, NSP12-13, NSP13-14, NSP14-15, and NSP15-16, respectively; one-way ANOVA *p* = 0.0038) of the NSP4-5 site was found to be significantly different (Fig. 7F; Supporting Table 2; Supporting Figure 10C). The differences in cleavage rate of NSP4-5 in all M^pro^ DuProSense constructs could result from the differences in the expression of either DuProSense biosensor constructs or the M^pro^. Another possibility of this observed differential cleavage rate of the same site (NSP4-5) is the presence of the second cleavage site in the DuProSense biosensor constructs. On the other hand, the t_1/2_ (8 ± 0.8 and 7.1 ± 0.2 h for NSP1-2 and NSP3-4, respectively; *p* = 0.1053) and cleavage rates (0.48 ± 0.08 and 0.52 ± 0.11 h^-1^ for NSP1-2 and NSP3-4, respectively; *p* = 0.5371) of NSP2-3 were found to be similar in PL^pro^ cleavage site containing DuProSense biosensor constructs (Fig. 7E,F; Supporting Table 2; Supporting Figure 10B,C).

Comparison of t_1/2_ values of M^pro^ cleavage sites revealed significantly higher t_1/2_ values for NSP5-6 (11.6 ± 3.5 vs. 19 ± 3 h; *p* = 0.0487), NSP6-7 (11.7 ± 3 vs. 25.3 ± 3.4 h; *p* = 0.0067), NSP7-8 (11.1 ± 1.7 vs. 18.3 ± 2.4 h; *p* = 0.0124), NSP10-11-12 (15 ± 1.9 vs. 23.9 ± 1.5 h; *p* = 0.0031), NSP12-13 (9.6 ± 1.6 vs. 18.8 ± 2.3 h; *p* = 0.0049), NSP13-14 (8.2 ± 3.4 vs. 20.4 ± 2.4 h; *p* = 0.0074), and NSP15-16 (8.2 ± 2.9 vs. 17.2 ± 4.2 h; *p* = 0.0391) sites, as compared to NSP4-5 site, in their respective DuProSense biosensors (Fig. 7E; Supporting Table 2; Supporting Figure 10D). However, comparison of cleavage rates revealed significantly slower rates for NSP6-7 (0.25 ± 0.03 vs. 0.13 ± 0.02 h^-1^; *p* = 0.0082), NSP8-9 (0.23 ± 0.06 vs. 0.12 ± 0.01 h^-1^; *p* = 0.0365), and NSP14-15 (0.22 ± 0.01 vs. 0.19 ± 0.01 h^-1^; *p* = 0.045), as compared to the NSP4-5 site (Fig. 7F; Supporting Table 2; Supporting Figure 10E). In contrast to the M^pro^ cleavage sites, no significant difference in the t1/2 (Fig. 7E; Supporting Table 2; Supporting Figure 10D) and cleavage rates (Fig. 7F; Supporting Table 2; Supporting Figure 10E) was observed for the PL^pro^ cleavage sites. While we found similar t_1/2_ and cleavage rates for all the PL^pro^ cleavage sites, previous reports on PL^pro^ enzyme kinetics determined through in vitro assay suggested a slower cleavage rate for the NSP1-3 and NSP3-4 site.^13^ One key difference between our live cell assay results and the previously reported in vitro assays is the use of the full-length PL^pro^ (NSP3) protein, which contains additional domains that can regulate the activity of the protease domain, in the live cells while the in vitro assays were performed with the protease domain alone. Additionally, PL^pro^(NSP3) is a transmembrane protein and localizes to endoplasmic reticulum membrane of host cells and contributes towards the biogenesis of double membrane vesicles (DVMs).^70^ While it is unclear at this stage how ER localization of PL^pro^ impacts its proteolytic activity, it is possible that such a restricted localization restricts substrate availability, and in turn, any difference in the cleavage rates of the three sites.

We then compared the two key parameters obtained from the kinetic sigmoidal fitting of the protease cleavage activity data, i.e. t_1/2_ and cleavage rate, for all the DuProSense biosensor constructs (Fig. 7G). This revealed a generally ∼2× faster rate for the PL^pro^ cleavage sites as compared to the M^pro^ cleavage sites in these experiments performed with cells transfected the proteases individually (Fig. 7G). Such an enhanced rate of cleavage was not observed in experiments with the DuProSense biosensor containing N-terminal cleavage sites of M^pro^ (NSP4-5) and PL^pro^ (NSP2-3) wherein both M^pro^ and PL^pro^ were expressed simultaneously in living cells (Fig. 6F). This could be due to a difference in the expression levels of the two proteases when they are either co-expressed or expressed together. Additionally, it is possible that the presence of two cleavage sites in the same DuProSense biosensor, such as in the biosensor containing NSP1-2 and 2-3 cleavage sites, increases the rate of their individual cleavage suggestive of an element of processivity in the mechanism of PL^pro^ activity, which is not possible in the case of the DuProSense biosensor containing one M^pro^ and on PL^pro^ cleavage site. The observed lag in NSP6-7 M^pro^ activity (Fig. 7E,F,G) might be the result of its high binding affinity to M^pro^ in intact as well as in cleaved product form.^71^ Interestingly, NSP13-14 demonstrated a high cleavage rate and a high relative t_1/2_ (Fig. 7E,F,G). However, NSP13-14 has been reported to have a medium cleavage rate and binding affinity in previous studies.^72^ Additionally, NSP10-11-12, a fast-cleaving site in our study, has been reported to be a slow cleavage site based on catalytic efficiency and substrate turnover number.^11,72^

Having observed a differential cleavage activity of some of the M^pro^ cleavage site containing DuProSense biosensors in living cells, we wondered whether these differences are also observed in vitro. We particularly focused our attention on the M^pro^ cleavage site containing DuProSense biosensors and not the PL^pro^ cleavage site containing biosensor since the later did not show significant differences in either their t_1/2_ or the rate of cleavage values. For this, we performed in vitro cleavage assays with a recombinantly purified M^pro^ protease and lysates prepared from cells expression all M^pro^ cleavage site containing DuProSense biosensors individually (Supporting Figure 11A). Incubation of various DuProSense containing lysates with M^pro^ resulted in similar, exponential rates of decreases in the green channel BRET, which represents the cleavage of the NSP4-5 site, whereas some differences were apparent in the rate of exponential decreases in the red channel BRET, which represents the cleavage of various M^pro^ cleavage sites other than NSP4-5, amongst the sites (Supporting Figure 11A). We fit the data to an exponential decay model to determine their t_1/2_ and rate of cleavage (Supporting Figure 11B,C) and determined their relative values against the NSP4-5 site (Supporting Figure 11D,E). This analysis revealed significantly longer t_1/2_ values for NSP5-6 (356.6 ± 52.1 vs. 756.9 ± 72.3 s; *p* = 0.0015), NSP6-7 (546.7 ± 71.2 vs. 7253.7 ± 733.7 s; *p* < 0.0001), NSP8-9 (559.2 ± 4.3 vs. 2120.3 ± 258.4 s; *p* = 0.0005), NSP9-10 (486.0 ± 108.8 vs. 785.4 ± 80.1 s; *p* < 0.0185), NSP10-11-12 (609.9 ± 74.7 vs. 1549.7 ± 214.5 s; *p* = 0.0020), and NSP13-14 (482.3 ± 60.8 vs. 815.4 ± 142.2 s; *p* < 0.0203) as compared to the t_1/2_ values of the NSP4-5 site in their respective DuProSense biosensors (Supporting Figure 11B,E). Similarly, the cleavage rates were found to be significantly slower for NSP5-6 (0.00197 ± 0.00032 vs. 0.00092 ± 0.00009 s; *p* = 0.005), NSP6-7 (0.00128 ± 0.00016 vs. 0.00010 ± 0.00001 s; *p* = 0.0002), NSP8-9 (0.00124 ± 0.00001 vs. 0.00033 ± 0.00004 s; *p* < 0.0001), NSP10-11-12 (0.00115 ± 0.00014 vs. 0.00045 ± 0.00007 s; *p* = 0.0016), NSP13-14 (0.00145 ± 0.00017 vs. 0.00087 ± 0.00014 s; *p* = 0.0105), NSP14-15 (0.00152 ± 0.00020 vs. 0.00080 ± 0.00022 s; *p* = 0.0141), and NSP15-16 (0.00134 ± 0.00022 vs. 0.00071 ± 0.00019 s; *p* = 0.0207), as compared to that of the NSP4-5 site in their respective DuProSense biosensors (Supporting Figure 11C,E). These results are in general agreement with previous in vitro studies.^11,13^

We then compared the relative t_1/2_ and cleavage rates of all M^pro^ cleavage sites determined from in vitro and live cell assays (relative to the t_1/2_ and cleavage rate of the NSP4-5 in their respective DuProSense biosensor), and determined the correlation (Fig. 7H,I). This revealed no significant correlation between the relative t_1/2_ values (*r* = 0.34; *p* = 0.3109) (Fig. 7H) but strong positive correlation between the cleavage rates (*r* = 0.70; *p* = 0.0159) (Fig. 7I) obtained from in vitro and live cell assays. The differences in the t_1/2_ and cleavage rates observed between the live cells and in vitro assays performed using DuProSense biosensors might have resulted from the differences in the DuProSense biosensor (substrate) and protease (enzyme) concentrations. In live cell assays, transfection of plasmid DNA expressing the DuProSense biosensors, and the protease will result in a continuous expression of the proteins over the period of the experiment (48 h). However, there is a finite concentration of DuProSense biosensor and protease in the in vitro assays. Additionally, the optimized conditions in the in vitro assays can change the structural and functional dynamics of proteins as compared to those in living cells. For example, the use of polyethylene glycol (PEG)^38^ and kosmotropic salts like sodium citrate^73^ increase the proteolytic activity of M^pro^ in in vitro assay most likely by stabilizing the M^pro^ dimer.^38^ Additionally, the intracellular interaction of M^pro^ and PL^pro^ with host cell proteins^74–76^ and the presence of putative host substrates^16,77^ can affect the enzyme kinetics in live cells.

We then took advantage of the M^pro^ site containing DuProSense biosensors to determine the proteolytic activity of SARS-CoV-1 M^pro^ and compare it with SARS-CoV-2 M^pro^ activity. The proteins share high sequence similarity with only 12, out of a total 306, residues being different. For this, we performed in vitro assays using a recombinantly purified SARS-CoV-1 M^pro^ and lysates prepared from cells expressing various M^pro^ site containing DuProSense biosensors (Supporting Figure 12) and compared the results with those obtained with SARS-CoV-2 M^pro^ in living cells (Supporting Figure 13A,C) and in vitro (Supporting Figure 13B,D) assays. The t_1/2_ values for various M^pro^ cleavage sites obtained with SARS-CoV-1 M^pro^ showed no significant correlation with t_1/2_ values obtained with SARS-CoV-2 M^pro^ in living cells (*r* = 0.44; *p* = 0.1778) and in vitro assay (*r* = 0.12; *p* = 0.7191) (Supporting Figure 13A,B). Additionally, a comparison of cleavage rates also revealed no significant correlation between SARS-CoV-1 M^pro^ in vitro and SARS-CoV-2 M^pro^ live cell assays (*r* = 0.0001; *p* = 0.9971) (Supporting Figure 12C). In addition to the general differences in the cleavage activity of SARS-CoV-1 and SARS-CoV-2 M^pro^, a key observation we made here is that the NSP6-7 was cleaved at a much higher rate by SARS-CoV-1 M^pro^ than SARS-CoV-2 M^pro^ (Supporting Figure 11C,12C). This could potentially arise from the residue-residue interactions and structural dynamic changes due to the difference of 12 residues, although the catalytic site residues are conserved, between SARS-CoV-1 and SAR-CoV-2 protein.^78^ Of particular interest is the amino acid residue at position 46, which is a S in SARS-CoV-2 M^pro^ (amino acid residue S46), which has been suggested to interact with NSP6-7^78,79^, but an A in SARS-CoV-1 M^pro^ (amino acid residue A46).

### Cleavage site-specific nirmatrelvir potency against M^pro^

Building on our findings related to differential cleavage kinetics of various M^pro^ cleavage sites using the DuProSense biosensors, we next investigated the potency of nirmatrelvir against M^pro^ in proteolytically processing various M^pro^ cleavage sites. Nirmatrelvir is an orally active SARS-CoV-2 M^pro^ inhibitor and is being administered as a COVID-19 antiviral drug along with ritonavir (Paxlovid, Pfizer).^80–82^ It is a catalytic site inhibitor and interacts with key binding site residues of M^pro^, namely M49, G143, M165, E166, G143, and Q189.^83,84^ Importantly, the SARS-CoV-2 M^pro^ cleavage site peptides have been reported to show a differential interaction with M^pro^ through the involvement of distinct sets of residues in the catalytic site of the protease.^11^ We therefore posit that binding of nirmatrelvir to M^pro^ could differentially impact the binding of the various M^pro^ cleavage site peptides and vice versa. For this, we first performed an assay with cells transfected with the M^pro^ expressing plasmid along with the different M^pro^ site containing DuProSense biosensor expressing plasmids individually, treated the cells with a single dose of nirmatrelvir (4 µM) and monitored BRET in the green (NSP4-5 site) and red (different M^pro^ cleavage sites) channel. We then determined the percentage inhibition of the NSP4-5 and the second M^pro^ cleavage site in the DuProSense biosensors individually (Supporting Figure 14A,B,C). This analysis revealed a significantly greater percentage M^pro^ inhibition in the case of DuProSense biosensor containing the NSP5-6 (15.5 ± 2.5 vs. 26 ± 1.1 %; *p* = 0.0027), NSP6-7 (15.3 ± 1.6 vs. 35.1 ± 3.4 %; *p* = 0.0008), NSP8-9 (16.0 ± 1.5 vs. 30.7 ± 8.2 %; *p* = 0.0375), NSP10-11-12 (16.3 ± 2.5 vs. 29 ± 3.2 %; *p* = 0.0057), NSP12-13 (14.0 ± 0.4 vs. 29.3 ± 0.9 %; *p* < 0.0001), and NSP13-14 (16.7 ± 2.1 vs. 34.3 ± 4.1 %; *p* = 0.0035), as compared to the inhibition observed for the NSP4-5 site in the same biosensor. Importantly, no significant differences were observed in the inhibition of M^pro^ activity on the NSP4-5 site sites (one-way ANOVA *p* =0.8842), while significant differences were observed for other sites (one-way ANOVA *p* < 0.0001) (Supporting Figure 14B), with a number of sites showing almost twice the level of inhibition (Supporting Figure 14C). Given that we had observed differences in the rate of cleavage of various M^pro^ cleavage sites containing DuProSense, we wondered if there is a relationship between the cleavage rate and the percentage inhibition of M^pro^. For this, we compared the cleavage rate and percentage M^pro^ inhibition observed with DuProSense biosensors containing M^pro^ cleavage sites and found no significant correlation either for NSP4-5 (*r* = 0.4; *p* = 0.2237) or other M^pro^ cleavage sites (*r* = −0.6; *p* = 0.070) (Supporting Figure 14D).

These results prompted us to investigate whether there are any differences in the potency (IC_50_ values) of nirmatrelvir inhibition of M^pro^ determined with the various M^pro^ cleavage site containing DuProSense biosensors and if they correlated with the cleavage kinetics of the sites. For this, we performed nirmatrelvir dose-response experiments with live cells expressing M^pro^ and various M^pro^ cleavage site containing DuProSense biosensors and monitored BRET in the green and the red channel after 22 h of treatment. We then determined the percentage M^pro^ cleavage activity with the NSP4-5 (green channel) and all other M^pro^ cleavage sites (red channel) (Fig. 8). Treatment of cells with a range of nirmatrelvir concentrations did not affect the expression of DuProSense biosensors as evident from the total bioluminescence measurements of various M^pro^ site containing DuProSense biosensors (Fig. 8A). Further, treatment of cells with nirmatrelvir resulted in a concentration-dependent decrease in M^pro^ activity (Fig. 8B), and as seen in the live cell cleavage kinetics experiments, no difference was observed in the inhibition of M^pro^ activity in the cleavage of the DuProSense biosensor containing NSP4-5 site in both the green and red channel biosensor modules (Fig. 8B,C). Importantly, in comparison to the nirmatrelvir IC_50_ for the NSP4-5 site, cells expressing DuProSense biosensors containing the NSP5-6 (13.4 ± 5.5 vs. 3.2 ± 0.7 µM; *p* = 0.0107), NSP6-7 (12.7 ± 4.1 vs. 1.3 ± 0.5 µM; *p* = 0.0017), NSP7-8 (17.3 ± 7.9 vs. 5.1 ± 0.4 µM; *p* = 0.0187), NSP8-9 (16.5 ± 7.2 vs. 3.1 ± 0.3 µM; *p* = 0.0061), NSP10-11-12 (13 ± 1.9 vs. 3.3 ± 1.9 µM; *p* = 0.0106), NSP12-13 (10.7 ± 1.6 vs. 3.8 ± 0.9 µM; *p* = 0.0024, and NSP13-14 (17.1 ± 5.3 vs. 6.1 ± 3.7 µM; *p* = 0.0495) sites showed a significantly lower IC_50_ value in their respective biosensors (Fig. 8B,C; Supporting Table 2), suggesting a greater potency of nirmatrelvir in inhibiting M^pro^ against these cleavage sites. In general, comparison of relative IC_50_ obtained as a ratio of the IC_50_ values obtained for various M^pro^cleavage sites and the NSP4-5 site in each of the DuProSense biosensors revealed significant difference across the various M^pro^ cleavage sites (one-way ANOVA, *p* < 0.0001) (Fig. 8D), with the NSP6-7 site showing lowest IC_50_ value (highest nirmatrelvir inhibition potency) and the NSP14-15 site showing the highest IC_50_ value (lowest nirmatrelvir inhibition potency) (Fig. 8D). These observations raised the possibility of a correlation between the nirmatrelvir IC_50_ values and the rate of cleavage of various M^pro^ cleavage sites. We, therefore, compared the IC_50_ values with the rate of cleavage obtained from live cell experiments (Fig. 8E). This analysis, however, revealed no significant correlation between the two parameters (r = 0.21; *p* = 0.5369), suggesting a more complex interplay between nirmatrelvir inhibition and cleavage rates in living cells (Fig. 8E). Nirmatrelvir is a competitive inhibitor of M^pro^ and therefore, differences in the affinity of various M^pro^ cleavage site peptides will lead to a differential competition between the inhibitor and the cleavage peptides. Previously, it has been reported that differences in binding kinetics of the substrates of a protein affect the inhibitory potency of the protein inhibitors.^85,86^ On the other hand, cleavage rates determined from live cell experiments are a composite of multiple factors including the cleavage site peptide affinity as well as catalytic turnover.

## Discussion

Since its initial use as a tool to measure protein-protein interaction, BRET has become a method of choice for a wide range of applications. However, it has typically been limited to detect a single molecular event. Results presented in this manuscript establish the possibility of simultaneously monitoring two molecular events in the sample. The presence of two BRET acceptor in a single protein construct does result in complexities arising from the occurrence of BRET from the donor (NLuc) to the acceptors (mNG and mSca) but also FRET from one acceptor (mNG) to the other acceptor (mSca). Importantly, we are able to resolve these complications through well controlled experiments using various biosensor constructs and, more importantly, innovative data analysis considering any FRET between the two acceptors in the biosensor constructs (Fig. 2,3,4). Thus, the design and validation of the DuProSense biosensor reported here opens the possibility of simultaneous characterization to cellular signaling events in a single assay. For one, it allowed us to compare the cleavage kinetics of SARS-CoV-2 M^pro^ and PL^pro^ proteases and determine the relative rates of cleavage of all their cleavage sites in living cells (Fig. 6,7). Based on our cleavage kinetics data, we propose the following model of SARS-CoV-2 pp1a and pp1ab polyprotein processing in cells (Fig. 9). SARS-CoV-2 positive-strand genomic RNA is translated into pp1a (containing NSP1 to 11) and pp1ab (containing NSP1 to 16) polyproteins from two overlapping open reading frames (ORFs), ORF1a and ORF1b, respectively.^87,88^ The ORF1b translation is mediated by a −1 programmed ribosomal frameshifting (−1 PRF) allowing the translation to continue beyond the stop codon of ORF1a.^88,89^ However, the frameshift efficiency of ORF1b is ∼60% which is expected to result in ∼40% lower expression of pp1ab, as compared to pp1a.^90^ This will, in turn, result in 1.6× expression of NSP1 to 11 and 0.6× expression of NSP12 to 16. Based on the cleavage rates obtained from our experiments, we classified the M^pro^ and PL^pro^ cleavage sites into very fast (relative rate ≥ 0.95), fast (0.85 ≤ relative rate ≤ 0.95), medium (0.75 ≤ relative rate ≤ 0.84), and slow (relative rate ≤ 0.74) cleaving. Thus, NSP1, NSP2, NPS3, NSP4, NSP5 and NSP11 proteins are released first. These are followed by the release of NSP12 (RdRp), NSP13 (helicase), NSP14, NSP15 and NSP16 from the intermediate polyprotein containing NSP12 to 16. Subsequently, NSP10 will be released from the intermediate polyprotein containing NSP8 to 9. Finally, NSP6, NSP7, NSP8 and NSP9 will be released from the intermediate polyproteins containing NSP6 and 7 and NSP8 and 9, respectively. Therefore, although the expression level of NSP12 to 16 is lower, their release from the polyprotein and thus, activation into functional forms is relatively faster (Fig. 9). These NSPs include RNA dependent RNA polymerase (NSP12), RNA helicase (NSP13), N7-methyltransferase/exoribonuclease (NSP14), endoribonuclease (NSP15), and NSP16 (2’-O-methyltransferase) and play an indispensable role in the viral genome replication and host immune suppression. This suggests that differential cleavage M^pro^ cleavage kinetics ensures a balance of critical proteins required for the viral replication and thus, SARS-CoV-2 infection. We note that while some of these critical NSPs are proteolytically processed at a faster rate, their cleavage by M^pro^ is inhibited by nirmatrelvir with a higher potency (Fig. 8), and thus, providing a mechanistic basis for the effectiveness of nirmatrelvir in COVID-19 treatment.

## Conclusion

To conclude, we have developed a biosensor platform of a genetically encoded, BRET-based, two-color, dual protease biosensors, DuProSense, that allows for simultaneous monitoring of the proteolytic activity of two proteases. We have extensively characterized the design using SARS-CoV-2 M^pro^ and PL^pro^ in terms of specificity of signal upon protease-mediated cleavage as well as pharmacological inhibition of the proteases. Live cell kinetic assays revealed similar cleavage activity of M^pro^ and PL^pro^. Further, while PL^pro^ showed similar cleavage kinetics for its sites, M^pro^ showed significantly different cleavage kinetics for its sites. While in vitro assays with recombinantly purified M^pro^ revealed differences in the rate of cleavage of M^pro^ cleavage sites, they were not correlated with the live cell cleavage rates. Importantly, live cell experiments with various M^pro^ cleavage site containing DuProSense biosensors revealed differences in the potency of nirmatrelvir against M^pro^ proteolytic activity. We believe that the DuProSense biosensor platform reported here will find direct utility in monitoring SARS-CoV-2 infection of cells in culture, drug discovery and functional genomics application. In this way, the DuProSense biosensor will be useful in our fight against the current as well as future threats from SARS-CoV-2 (and similar beta-coronaviruses) and their ever-evolving variants. More importantly, the DuProSense biosensor platform can be extended to any pair of proteases, including human host proteases. In general, the dual color BRET assay developed here can be advantageous applied to simultaneously monitor two signaling events or pathways in diseases such as pathogenic infections or cancer.

## Materials and Methods

### BRET-based dual protease biosensor (DuProSense biosensor) plasmid construct generation

We generated four DuProSense biosensor constructs containing mNG at the N-terminal and mBeRFP, LSS-mKate2, CyOFP1, and mSca RFPs at the C-terminal of NLuc luciferase protein. The DuProSense biosensor constructs were generated utilizing previously reported M^pro^ (amino acid sequence: AVLQSGFR; nucleotide sequence 5’ GCA GTG CTC CAA AGC GGA TTT CGC 3’) and PL^pro^ (amino acid sequence: LKGGAPTK; nucleotide sequence 5’ CTG AAA GGC GGC GCG CCG ACC AAA 3’) N-terminal autocleavage peptide sequences.^10,13,91^ First, a gene fragment containing a part of the mNG coding sequence, M^pro^ N-terminal autocleavage site, NLuc, PL^pro^ N-terminal autocleavage sequence and mBeRFP (mNG-M^pro^-NLuc-PL^pro^-mBeRFP) was synthesized with *Bst*XI and *Xba*I restriction enzyme sites at the 5’ and 3’ termini, respectively (GenScript, Singapore) and subcloned into the pUC57 plasmid to generate the pUC57-mNG-M^pro^-NLuc-PL^pro^-mBeRFP plasmid construct. The DNA fragment, mNG-M^pro^-NLuc-PL^pro^-mBeRFP was excised from the pUC57-mNG-M^pro^-NLuc-PL^pro^-mBeRFP plasmid using the restriction enzymes *Bst*XI and *Xba*I and ligated to the similarly digested mNeonGreen-DEVD-nanoLuc plasmid ((a gift from Maarten Merkx; Addgene plasmid # 98287; http://n2t.net/addgene:98287; RRID:Addgene_98287)^92^ to generate the pmNG-M^pro^-NLuc-PL^pro^-mBeRFP (DuProSense biosensor with mBeRFP as the RFP) plasmid. Subsequently, LSS-mKate2, CyOFP1 and mSca gene fragments were PCR amplified from pLSSmKate2-C1 (a gift from Vladislav Verkhusha; Addgene plasmid # 31869; http://n2t.net/addgene: 31869; RRID:Addgene_31869)^93^, pcDNA3-Antares2 c-myc (a gift from Huiwang Ai; Addgene plasmid # 100027; http://n2t.net/addgene: 100027; RRID:Addgene_100027)^94^ and pmScarlet_C1 (a gift from Dorus Gadella; Addgene plasmid # 85042; http://n2t.net/addgene: 85042; RRID:Addgene_85042)^95^ plasmids, respectively, using forward primers containing an *EcoR*I restriction enzyme site and reverse primers containing *Kpn*I restriction enzyme (Supporting Table 1). The PCR products and the pmNG-M^pro^-NLuc-PL^pro^-mBeRFP plasmid were digested using *EcoR*I and *Kpn*I restriction enzymes and ligated using T4 DNA ligase (Thermofisher Scientific). The clones pmNG-M^pro^-NLuc-PL^pro^-LSS-mKate2, pmNG-M^pro^-NLuc-PL^pro^-CYOFP1, and pmNG-M^pro^-NLuc-PL^pro^-mSca were confirmed by restriction-digestion. All plasmid sequences were further confirmed by Sanger sequencing (Macrogen, South Korea) using a forward primer complementary to the cytomegalovirus (CMV) promoter nucleotide sequence and respective reverse primers (Supporting Table 1). All four DuProSense biosensor constructs contained a His6-tag and a 3×FLAG-tag at the N- and C-termini, respectively.

The M^pro^ cleavage site mutant of the mSca-containing DuProSense biosensor (pmNG-M^pro^-mut-NLuc-PL^pro^-mSca) was generated by substituting the amino acid residue Q to A at the P1 site in the autocleavage peptide sequence of M^pro^ (AVL**A**SGFR; nucleotide sequence 5’ GCA GTG CTC **GCC** AGC GGA TTT CGC 3’). The PL^pro^ cleavage site mutant of the mSca-containing DuProSense biosensor (pmNG-M^pro^-NLuc-PL^pro^-mut-mSca) was generated by replacing the amino acid residue G to I in the PL^pro^ cleavage site (LKG**I**APTK; nucleotide sequence 5’ CTG AAA GGC **ATT** GCG CCG ACC AAA)^10^. The M^pro^ cleavage site and PL^pro^ cleavage site double mutant (pmNG-M^pro^-mut-NLuc-PL^pro^-mut-mSca) was generated by substituting the amino acid residue G to I in the PL^pro^ cleavage site (LKG**I**APTK; nucleotide sequence 5’ CTG AAA GGC **ATT** GCG CCG ACC AAA)^10^ of the M^pro^ cleavage site mutant of the mSca-containing DuProSense biosensor plasmid (pmNG-M^pro^-mut-NLuc-PL^pro^-mSca) (GenScript Biotech (Singapore) Pte. Ltd., Singapore).

The DuProSense biosensors containing various M^pro^ sites were generated by replacing the PL^pro^ N-terminal autocleavage site peptide sequence (LKGGAPTK; nucleotide sequence 5’ CTG AAA GGC GGC GCG CCG ACC AAA 3’) of mSca-containing DuProSense with M^pro^ cleavage sites (Supporting Table 3) (GenScript Biotech (Singapore) Pte. Ltd., Singapore). The DuProSense biosensors containing various PL^pro^ sites were generated by replacing the M^pro^ N-terminal autocleavage site peptide sequence (AVLQSGFR; nucleotide sequence 5’ GCA GTG CTC CAA AGC GGA TTT CGC 3’) of mSca-containing DuProSense with PL^pro^ cleavage sites (Supporting Table 3) (GenScript Biotech (Singapore) Pte. Ltd., Singapore).

### Cell Culture and transfection

HEK293T cells were used to perform both live cell assays and for preparing DuProSense biosensor containing cell lysates for in vitro assays. Cells were cultured in Dulbecco’s Modified Eagle Medium (DMEM) supplemented with 10% fetal bovine serum, and 1% penicillin-streptomycin at 37 °C in 5% CO_2_. For live cell BRET assays, HEK293T in 96-well white plates were transfected using polyethyleneimine (PEI) lipid (Sigma-Aldrich; 408727-100mL) after 24 h of seeding. Briefly, the plasmid DNA and Opti-MEM (Invitrogen; 31985088) were mixed using pipetting and brief vortex after the addition of 1.25 μg/well of PEI lipid. The transfection mix was incubated at room temperature for 30 min before adding to wells drop by drop. For the optimization of dual color BRET calculations, HEK293T cells were transfected with pNLuc, pmNG-NLuc, pNLuc-mSca, and pmNG-NLuc-mSca individually (Fig. 2). For spectral characterization of DuProSense with various RFPs, HEK293T cells were transfected individually with pmNG-M^pro^-NLuc-PL^pro^-mBeRFP, pmNG-M^pro^-NLuc-PL^pro^-LSS-mKate2, pmNG-M^pro^-NLuc-PL^pro^-CyOFP1, and pmNG-M^pro^-NLuc-PL^pro^-mScarlet (Fig. 3). To monitor the cleavage activity the M^pro^ and PL^pro^ individually, HEK293T cells were co-transfected with mSca-containing DuProSense biosensor plasmid along with either pLVX-EF1alpha-SARS-CoV-2-nsp5-2×Strep-IRES-Puro (M^pro^ WT) (a gift from Nevan Krogan; Addgene plasmid # 141370; http://n2t.net/addgene:141370; RRID:Addgene_14137)^96^ or Nsp3 -EGFP (PL^pro^ WT) (a gift from Bruno Antonny; Addgene #165108; http://n2t.net/addgene:165108; RRID:Addgene_165108)^97^ plasmid at a ratio of 1:5 of biosensor-to-protease plasmid DNA (Fig. 4-6). For simultaneous monitoring of the M^pro^ and PL^pro^ cleavage activity, HEK293T cells were co-transfected with mSca-containing DuProSense biosensor plasmid along with pLVX-EF1alpha-SARS-CoV-2-nsp5-2×Strep-IRES-Puro (M^pro^ WT) (a gift from Nevan Krogan; Addgene #141370; http://n2t.net/addgene:141370; RRID:Addgene_14137)^96^ and Nsp3 -EGFP (PL^pro^ WT) (a gift from Bruno Antonny; Addgene #165108; http://n2t.net/addgene:165108; RRID:Addgene_165108)^97^ plasmid at a ratio of 1:5:5 of biosensor-to-M^pro^ and PL^pro^ plasmid DNA (Fig. 4,6). For determining the temporal cleavage dynamics of all M^pro^ and site-specific M^pro^ inhibitory potency in live cells, HEK293T cells were co-transfected with DuProSense biosensor containing M^pro^ cleavage sites and pLVX-EF1alpha-SARS-CoV-2-nsp5-2×Strep-IRES-Puro (M^pro^ WT) (a gift from Nevan Krogan; Addgene #141370; http://n2t.net/addgene:141370; RRID:Addgene_14137)^96^ (Fig. 7,8). For determining the temporal cleavage dynamics of all PL^pro^ cleavage sites, HEK293T cells were co-transfected with DuProSense biosensor constructs containing PL^pro^ cleavage sites and Nsp3 -EGFP (PL^pro^ WT) (a gift from Bruno Antonny; Addgene #165108; http://n2t.net/addgene:165108; RRID:Addgene_165108)^97^ plasmid (at a 1:5 ratio of DuProSense biosensor plasmid DNA to protease) (Fig. 7). HEK293T cells were seeded onto 10 cm dishes and were transfected after 24 h for collecting cell lysates used in in vitro assays and western blots. To check the expression and cleavage of DuProSense biosensors upon M^pro^ and PL^pro^ expression, HEK293T cells were transfected with pmNG-M^pro^-NLuc-PL^pro^-mBeRFP, pmNG-M^pro^-NLuc-PL^pro^-LSS-mKate2, pmNG-M^pro^-NLuc-PL^pro^-CyOFP1, and pmNG-M^pro^-NLuc-PL^pro^-mScarlet either alone or co-transfected with either pLVX-EF1alpha-SARS-CoV-2-nsp5-2×Strep-IRES-Puro (M^pro^ WT) (a gift from Nevan Krogan; Addgene #141370; http://n2t.net/addgene:141370; RRID:Addgene_14137)^96^ or Nsp3 -EGFP (PL^pro^ WT) (a gift from Bruno Antonny; Addgene #165108; http://n2t.net/addgene:165108; RRID:Addgene_165108)^97^ plasmid or both at a ratio of 1:5 of biosensor-to-protease plasmid DNA (Fig. 3). For in vitro M^pro^ cleavage kinetics assays and FRET measurements, the HEK293T cells were transfected with DuProSense biosensor constructs containing various M^pro^ cleavage sites (Supporting Figure 6,11,12).

### Cell lysate preparation

For cell lysate preparation, HEK293T cells were seeded onto 10 cm dishes and were transfected with relevant plasmid DNA. After 48 h of transfection, dishes containing cells were transferred on ice and cell culture media was aspirated out from the dishes. As done previously^27,31,43,52,98–103^, cells were then washed with ice-cold Dulbecco’s Phosphate-Buffered Saline (DPBS). After DPBS wash, 400 µL of chilled mammalian cells lysis buffer (50 mM HEPES (pH 7.5), 50 mM NaCl, 0.1% Triton-X 100, 1 mM Dithiothreitol (DTT) & 1 mM ethylenediamine tetraacetic acid (EDTA)) was added onto the cells^38^. Cells were then collected into 1.5 mL Eppendorf tubes and centrifuged at 14,000 rotations per minute (rpm) for 1 h. After centrifugation, supernatants were collected and stored at −80 °C until further usage.

### Bioluminescence and fluorescence measurements

Bioluminescence and fluorescence measurements were performed using the Tecan SPARK® multimode microplate reader. Bioluminescence spectral scans were acquired according to the previously described protocol.^38,104,105^ For in vitro fluorescent spectral analysis, an equal volume of cell lysates prepared from cells expressing the four DuProSense biosensor constructs (mBeRFP, LSS-mKate2, CyOFP1, and mSca) were excited at 440 nm, and fluorescence spectra were measured. For determining FRET between mNG and mSca in various M^pro^ and PL^pro^ cleavage sites containing DuProSense biosensors, cell lysates expressing these DuProSense biosensors were excited with 440 nm light and emissions were measured at 520 nm and 590 nm. FRET efficiency was calculated as a ratio of emission at 590 and 520 nm:

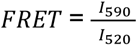

### Bioluminescence spectra measurement and optimization of dual BRET quantification

The bioluminescent spectra of live HEK293T cells transfected with relevant plasmids were acquired 48 h post-transfection by adding furimazine (Promega, Wisconsin, USA) at a dilution of 1:200. Three independent experiments were performed in triplicates. The optimization of dual BRET calculation was performed using the bioluminescence spectra of the cells expressing NLuc, mNG-NLuc, NLuc-mSca, and mNG-NLuc-mSca. The equations for the conventional green channel or M^pro^ biosensor BRET and conventional red channel or PL^pro^ biosensor BRET are shown below:

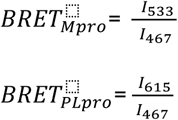

The equations for optimized green channel or M^pro^ biosensor BRET and optimized red channel or PL^pro^ biosensor BRET calculations are shown below:

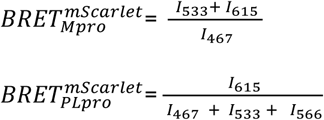

The equations for optimized green channel or M^pro^ biosensor BRET and optimized red channel or PL^pro^ biosensor BRET calculations for DuProSense biosensor with mBeRFP, LSSmKate2, and CyOFP1 are shown below:

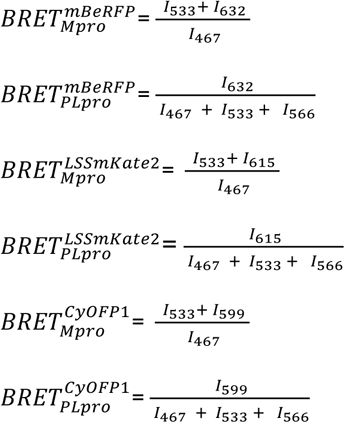

where *I*_533_ (mNG peak emission wavelength)*, I*_615_ (mScarlet peak emission wavelength)*, I*_632_ (mBeRFP peak emission wavelength)*, I*_599_ (LSSmKate2 peak emission wavelength)*, and I*_467_ (NLuc peak emission wavelength) are relative intensities obtained from bioluminescence spectra of the indicated DuProSense biosensor constructs (Fig. 4).

### Plasmid concentration-dependent protease activity live cell assay

HEK293T cells were transfected with mSca-containing DuProSense biosensor plasmid (25 ng/well) along with either pLVX-EF1alpha-SARS-CoV-2-nsp5-2×Strep-IRES-Puro (M^pro^) (a gift from Nevan Krogan; Addgene #141370; http://n2t.net/addgene:141370; RRID:Addgene_14137)^96^ or Nsp3 -EGFP (PL^pro^) (a gift from Bruno Antonny; Addgene #165108; http://n2t.net/addgene:165108; RRID:Addgene_165108)^97^ plasmid in a range of biosensor to protease ratio of 1:0.0005 to 1:5 (protease plasmid DNA concentration 0.0125, 0.125, 1.25, 12.5, 125 ng). For the combinatorial protease dose-response experiment, equal amounts of both pLVX-EF1alpha-SARS-CoV-2-nsp5-2×Strep-IRES-Puro (M^pro^) (a gift from Nevan Krogan; Addgene #141370; http://n2t.net/addgene:141370; RRID:Addgene_14137)^96^ and Nsp3 -EGFP (PL^pro^) (a gift from Bruno Antonny; Addgene #165108; http://n2t.net/addgene:165108; RRID:Addgene_165108)^97^ were co-transfected with the mSca-containing DuProSense biosensor plasmid DNA. Bioluminescence spectra were monitored 48 h post-transfection (Supporting Figure 2).

### Live cell protease activity assay

HEK293T cells were transfected with relevant plasmid DNA in 96-well white flat bottom plates. For spectral characterization and protease activity monitoring, BRET measurements were performed by the addition of furimazine (Promega, Wisconsin, USA) at a dilution of 1:200 after 48 h transfection (Fig 2-4). For live cells temporal cleavage dynamics assays, BRET measurements were performed by the addition of furimazine (Promega, Wisconsin, USA) at a dilution of 1:200 at 2h, 4h, 6h, 8h, 10h, 12h, 14, 16, 18, 20h, 22 h, 24h, 28 h, 36 h, and 48 h post-transfection (Fig 5,6). Three independent experiments were performed in triplicates. The total bioluminescence was calculated by adding the luminescence values obtained for each wavelength. The BRET measurements were calculated as previously described. The % protease activity was calculated from BRET as given below:

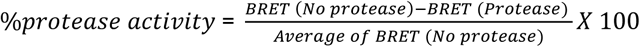

Kinetic sigmoidal nonlinear fit was performed using the equation:

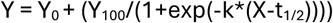

### Live cell inhibition of M^pro^ and PL^pro^ by GC376 and GRL-0617

HEK293T cells were co-transfected with mSca-containing DuProSense biosensor plasmid DNA and either pLVX-EF1alpha-SARS-CoV-2-nsp5-2×Strep-IRES-Puro (M^pro^) (a gift from Nevan Krogan; Addgene #141370; http://n2t.net/addgene:141370; RRID:Addgene_14137)^96^ or Nsp3-EGFP (PL^pro^) (a gift from Bruno Antonny; Addgene #165108; http://n2t.net/addgene:165108; RRID:Addgene_165108)^97^ plasmid DNA in 96-well white flat bottom plates. The ratio of biosensor-to-protease plasmid DNA was maintained at 1:5. To check the utility of DuProSense biosensor as an inhibitory assay reporter, transfected cells were simultaneously treated with a range of concentrations of corresponding protease inhibitor (GC376 against M^pro^, GRL-0617 against PL^pro^; both inhibitor stock solutions in 50% DMSO). After 22 h of incubation with the inhibitor, BRET measurements were taken by adding furimazine (Promega, Wisconsin, USA) at a dilution of 1:200. The percentage protease activity was calculated by normalizing the BRET values with negative control (in the absence of protease) and positive control (with the treatment of compound having no inhibitory effect on protease; GRL-0617 on M^pro^, GC376 on PL^pro^).

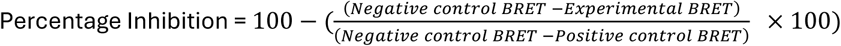

### Live cell inhibition of M^pro^ by nirmatrelvir

HEK293T cells were co-transfected with mSca-containing DuProSense biosensor plasmid DNA and pLVX-EF1alpha-SARS-CoV-2-nsp5-2×Strep-IRES-Puro (M^pro^) (a gift from Nevan Krogan; Addgene #141370; http://n2t.net/addgene:141370; RRID:Addgene_14137)^96^ plasmid DNA in 96-well white flat bottom plates. Simultaneously, the cells were treated with a range of doses (100, 33.3, 10, 3.3, 1, 0.3, 0.1, 0.03 µM) of nirmatrelvir (Sigma; 4 mM stock solution in DMSO) and BRET measurements were taken after 22 h by adding furimazine (Promega, Wisconsin, USA) at a dilution of 1:200.

### Western blot analysis

HEK293T cells co-transfected with the relevant plasmid DNA were lysed in 200 μL of 2× Laemmli sample buffer (50 mM Tris-Cl pH 6.8, 1.6% SDS, 8% glycerol, 4% β-mercaptoethanol (β-ME) and 0.04% bromophenol blue) (heated to 95 °C and sonicated prior to addition). Western blot was carried out as previously described^38,106^. Briefly, equal volumes of the cell lysates (30 μL) were separated by 10% SDS-PAGE followed by transferring proteins onto PVDF (Polyvinylidene fluoride) membranes. Membranes were blocked in Tris-buffered saline containing 0.1% Tween-20 (TBS-T) with bovine serum albumin (BSA – 5%) for 1 h at room temperature. Blots were incubated with anti-FLAG-tag antibody (DYKDDDDK Tag Mouse Monoclonal Antibody (FG4R); ThermoFisher Scientific-MA1-91878; 1:500) overnight at 4 °C in dilution buffer (TBS-T containing 5% BSA). Secondary anti-mouse IgG HRP (Anti-Mouse IgG:HRP Donkey pAb; ECM biosciences-MS3001; 1:10000 diluted in TBS-T) was used to detect intact and cleaved DuProSense biosensor proteins.

### Expression and purification of SARS-CoV-2 M^pro^

The chemically transformed BL21-CodonPlus (DE3) E. coli strain with the SARS-CoV-2 M^pro^ bacterial expression plasmid DNA, pETM33_NSP5_M^pro^ (a gift from Ylva Ivarsson; Addgene plasmid # 156475; http://n2t.net/addgene:156475; RRID:Addgene_156475)^107^ was grown in Luria Broth (LB) media containing 50 μg/mL kanamycin and chloramphenicol overnight at 180 rpm and 37 °C. Then, the inoculum was transferred to the fresh LB media and incubated for 2 h at 37 °C and 220 rpm, followed by induction of protein expression using IPTG at the concentration of 1 mM for 2.30 h at 37 °C and 220 rpm. The cells were pelleted at 4,000×g at 4 °C for 10 minutes and resuspended in 10 mL bacterial cells lysis buffer (50 mM Tris (pH 8), 300 mM NaCl, 10 mM β-ME, 1 mM PMSF, 10% (v/v) glycerol), followed by 15-30 minutes sonication on ice and centrifugation at 4,000× g at 4 °C for 10 minutes. The supernatant was collected and centrifuged at 18,000×g at 4 °C for 1 h, followed by incubation with the GSH beads for 2 h. The GSH beads were washed with buffer containing 50 mM Tris (pH 7), 150 mM NaCl, 10 mM β-ME, 1 mM EDTA, 10% (v/v) glycerol, and 0.01% TritonX-100. The GSH beads bound with M^pro^ were incubated with PreScission Protease (GE Healthcare # 27-0843-01) in cleavage buffer (50 mM Tris (pH 7), 150 mM NaCl, 1 mM EDTA, 1 mM DTT, 10% (v/v) glycerol and 0.01% TritonX-100) for 16 h at 4 °C. The supernatant containing the SARS-CoV-2 M^pro^ was obtained after centrifugation at 4 °C and 500× g was aliquoted and stored at −80 °C until further usage.^38,103^

### In vitro DuProSense biosensor BRET assay

Lysates from HEK293T cells transfected with relevant plasmid DNA were collected as described previously.^27,31,43,52,98,100–102,106^ An equal amount of cell lysates normalized using mNG fluorescence were incubated with 5 μM of recombinantly purified SARS-CoV-2 M^pro^ in CoV-2 M^pro^ assay buffer (TBS containing 1 M sodium citrate, 2 mM DTT & 1 mM EDTA) or 500 nM of SARS-CoV-1 M^pro^ (SARS coronavirus, 3CL Protease, Recombinant from E. coli; NR-700; BEI Resources, NIAID, NIH; stock solution of the protein was prepared by dissolving the lyophilized protein in TBS containing 10% glycerol) at a concentration of 50 μM) in mammalian cells lysis buffer and 25% PEG 8000. BRET was monitored through bioluminescence scans. BRET spectral scan measurements were performed at 37 °C using a Tecan SPARK® multimode microplate reader after the addition of furimazine (Promega, Wisconsin, USA) at a dilution of 1:200 as previously described.^38,104,105^

### Data analysis and figure preparation

GraphPad Prism (version 9 for macOS, GraphPad Software, La Jolla California USA; www.graphpad.com), in combination with Microsoft Excel, was used for data analysis and graph preparation. Figures were assembled using Adobe Illustrator.

## Data availability

All the relevant data of this study are available within this paper and the Supporting Information file.

## Acknowledgments

This work is supported by an Industrial Innovation Fund from the Office of Innovation & Industrial Relation, Hamad Bin Khalifa University (HBKU) (HBKU-INT-IC-IIF-05-05), an Academic Research Grant (ARG) from the Qatar Research and Development Innovation Council (QRDI) (ARG01-0517-230211) and internal funding from the College of Health & Life Sciences (CHLS), HBKU, a member of the Qatar Foundation. A.F. and S.M.N.U. are supported by student scholarships and A.M.G. is supported by a postdoctoral fellowship from CHLS, HBKU, a member of the Qatar Foundation.

## Author Contributions

K.H.B. conceived experiments. A.F., A.M.G, S.M.N.U and K.H.B. performed experiments, analyzed data, prepared figures, and wrote, reviewed, and approved the manuscript.

## Competing interests

A US utility patent (18/763,140) application with K.H.B, A.F. & A.M.G. as inventors has been filed.

## Supporting Information

Document DuProSense S1. Supporting Table 1-3, Supporting Text, and Supporting Figures 1-14

