## Supplementary material for "Differential Kinetics of SARS-CoV-2 Proteases Revealed by a Dual-Color, BRET-based Protease Biosensor, DuProSense": . Supporting Table 1-3, Supporting Text, and Supporting Figures 1-14

Affiliations:
[1]Division of Biological and Biomedical Sciences, College of Health & Life Sciences, Hamad Bin Khalifa University, Education City, Qatar Foundation, Doha – 34110, Qatar

[2]Equal contribution

ORCID:
Kabir H Biswas: 0000-0001-9194-4127
Anupriya M Geethakumari: 0000-0002-8932-9114
Asma Fatima: 0009-0004-7191-5109
S M Nasir Uddin: 0009-0006-4264-2302

### Supporting Table

**Supporting Table 1. List of primers for RFP gene amplification.**

| | Primer | Primer name | Sequence |
|---|---|---|---|
| 1. | Forward Primer | mSca-EcoRI-F | GCGCGGAATTCATGGTGAGCAAGGGCGAGGC |
| 2. | Reverse Primer | mSca-KpnI-R | GCGCCGGTACCCTTGTACAGCTCGTCCATGCCGCCGG |
| 3. | Forward Primer | CyOFP1-EcoRI-F | GGCGCGAATTCATGGTGAGCAAGGGCGAGGAGCTG |
| 4. | Reverse Primer | CyOFP1-KpnI-R | GCAGCGGTACCAACGAAATCTTCGAGTGTGAAGCCTCCGCCC |
| 5. | Forward Primer | LSS-mKate2-EcoRI F | GCGCGGAATTCATGGTGTCTAAGGGCGAAGAGC |
| 6. | Reverse Primer | LSS-mKate2-KpnI R | GCGCGGGTACCATTAAGCTTGTGCCCCAGTTTGC |

**Supporting Table 2. Table showing various parameters obtained from experiments performed with all SARS-CoV-2 protease ($M^{pro}$ and $PL^{pro}$) cleavage site biosensors.**

| | | | $BRET_{max}$ | | $t_{1/2}$ | | | Rate | | | Nirmatrelvir $IC_{50}$ | | |
|---|---|---|---|---|---|---|---|---|---|---|---|---|---|
| # | Protease | DuProSense Biosensor | Green biosensor (%) (mean ± SD; *N* = 3) | Red biosensor (%) (mean ± SD; *N* = 3) | Green biosensor (h) (mean ± SD; *N* = 3) | Red biosensor (h) (mean ± SD; *N* = 3) | *p* (Student's t-test) | Green biosensor ($h^{-1}$) (mean ± SD; *N* = 3) | Red biosensor ($h^{-1}$) (mean ± SD; *N* = 3) | *p* (Student's t-test) | Green biosensor (μM) (mean ± SD; *N* = 3) | Red biosensor (μM) (mean ± SD; *N* = 3) | *p* (Student's t-test) |
| 1 | | **NSP4-5/4-5** | 88 ± 0.5 | 84 ± 0.9 | 13.3 ± 0.6 | 12 ± 5 | 0.6729 | 0.18 ± 0.01 | 0.16 ± 0.01 | 0.2099 | 37.9 ± 6.9 | 29.5 ± 13.9 | 0.3237 |
| 2 | | **NSP4-5/5-6** | 89 ± 2.5 | 84 ± 2.8 | 11.6 ± 3.5 | 19 ± 3 | 0.0487 | 0.22 ± 0.03 | 0.22 ± 0.04 | 0.8656 | 13.4 ± 5.5 | 3.2 ± 0.7 | 0.0107 |
| 3 | | **NSP4-5/6-7** | 87 ± 1.9 | 58 ± 3.2 | 11.7 ± 3 | 25.3 ± 3.4 | 0.0067 | 0.25 ± 0.03 | 0.13 ± 0.02 | 0.0082 | 12.7 ± 4.1 | 1.3 ± 0.5 | 0.0017 |
| 4 | | **NSP4-5/7-8** | 87 ± 3.8 | 85 ± 3.6 | 11.1 ± 1.7 | 18.3 ± 2.4 | 0.0124 | 0.2 ± 0.04 | 0.19 ± 0.04 | 0.7283 | 17.3 ± 7.9 | 5.1 ± 0.4 | 0.0187 |
| 5 | | **NSP4-5/8-9** | 85 ± 6.1 | 77 ± 6.1 | 12.4 ± 2.3 | 20.6 ± 4.2 | 0.0416 | 0.23 ± 0.06 | 0.12 ± 0.01 | 0.0365 | 16.5 ± 7.2 | 3.1 ± 0.3 | 0.0062 |
| 6 | $M^{pro}$ | **NSP4-5/9-10** | 87 ± 3.7 | 85 ± 2.7 | 11.1 ± 2.3 | 15.6 ± 2.7 | 0.0938 | 0.24 ± 0.01 | 0.19 ± 0.04 | 0.116 | 21.9 ± 10.8 | 9.9 ± 2.8 | 0.1516 |
| 7 | | **NSP4-5/10-11-12** | 88 ± 1.6 | 80 ± 3.1 | 15 ± 1.9 | 23.9 ± 1.5 | 0.0031 | 0.19 ± 0.02 | 0.19 ± 0.01 | 0.8146 | 13 ± 1.9 | 3.3 ± 1.9 | 0.0106 |
| 8 | | **NSP4-5/12-13** | 90 ± 1.4 | 84 ± 2 | 9.6 ± 1.6 | 18.8 ± 2.3 | 0.0049 | 0.22 ± 0.04 | 0.2 ± 0.02 | 0.3727 | 10.7 ± 1.6 | 3.8 ± 0.9 | 0.0024 |
| 9 | | **NSP4-5/13-14** | 89 ± 2.1 | 84 ± 3 | 8.2 ± 3.4 | 20.4 ± 2.4 | 0.0074 | 0.14 ± 0.03 | 0.16 ± 0.01 | 0.4191 | 17.1 ± 5.3 | 6.1 ± 3.7 | 0.0495 |
| 10 | | **NSP4-5/14-15** | 89 ± 2.9 | 83 ± 7.1 | 14 ± 3.9 | 20.6 ± 1.4 | 0.0519 | 0.22 ± 0.01 | 0.19 ± 0.01 | 0.045 | 14.9 ± 5.6 | 20 ± 6.1 | 0.3075 |
| 11 | | **NSP4-5/15-16** | 88 ± 4 | 84 ± 5.6 | 8.2 ± 2.9 | 17.2 ± 4.2 | 0.0391 | 0.14 ± 0.03 | 0.12 ± 0.01 | 0.579 | 22.5 ± 17.3 | 7.4 ± 2 | 0.1022 |
| 12 | $PL^{pro}$ | **NSP1-2/2-3** | 88 ± 0.6 | 87 ± 0.6 | 8 ± 0.8 | 9.5 ± 0.9 | 0.0877 | 0.48 ± 0.08 | 0.4 ± 0.09 | 0.3017 | | | |
| 13 | | **NSP3-4/2-3** | 89 ± 0.8 | 87 ± 0.9 | 7.1 ± 0.2 | 8.3 ± 0.5 | 0.019 | 0.52 ± 0.11 | 0.36 ± 0.08 | 0.1152 | | | |

**Supporting Table 3. Cleavage site sequences are present in DuProSense biosensors containing SARS-CoV-2 protease ($M^{pro}$ and $PL^{pro}$) cleavage sites.**

| Sl. No. | Cleavage sites | DuProSense biosensor | Green biosensor; cleavage site sequence | Red biosensor; cleavage site sequence |
|---|---|---|---|---|
| 1 | | **NSP4-5/4-5** | AVLQSGFR | AVLQSGFR |
| 2 | | **NSP4-5/5-6** | AVLQSGFR | VTFQSAVK |
| 3 | | **NSP4-5/6-7** | AVLQSGFR | ATVQSKMS |
| 4 | | **NSP4-5/7-8** | AVLQSGFR | ATLQAIAS |
| 5 | | **NSP4-5/8-9** | AVLQSGFR | VKLQNNEL |
| 6 | $M^{pro}$ | **NSP4-5/9-10** | AVLQSGFR | VRLQAGNA |
| 7 | | **NSP4-5/10-11-12** | AVLQSGFR | PMLQSADA |
| 8 | | **NSP4-5/12-13** | AVLQSGFR | TVLQAVGA |
| 9 | | **NSP4-5/13-14** | AVLQSGFR | ATLQAENV |
| 10 | | **NSP4-5/14-15** | AVLQSGFR | TRLQSLEN |
| 11 | | **NSP4-5/15-16** | AVLQSGFR | PKLQSSQA |
| 12 | $PL^{pro}$ | **NSP1-2/2-3** | LNGGAYTR | LKGGAPTK |
| 13 | | **NSP3-4/2-3** | LKGGKIVN | LKGGAPTK |

### Supporting Text

#### Amino acid sequence of BRET-based SARS-CoV-2 DuProSense biosensor constructs:

**mNG-M$^{pro}$-Nter-auto-NLuc-PLpro-Nter-auto-mBeRFP**:

MGSS**HHHHHH**SSGLVPRGSHMVSKGEEDNMASLPATHELHIFGSINGVDFDMVGQGTGNPNDGYEELNLKSTKGDLQFSPWILVPHIGYGFHQYLPY
PDGMSPFQAAMVDGSGYQVHRTMQFEDGASLTVNYRYTYEGSHIKGEAQVKGTGFPADGPVMTNSLTAADWCRSKKTYPNDKTIISTFKWSYTTGNG
KRYRSTARTTYTFAKPMAANYLKNQPMYVFRKTELKHSKTELNFKEWQKAFTDVMGMDELYKAAATENLY**AVLQSGFR**GSGSAMVFTLEDFVGDWRQ
TAGYNLDQVLEQGGVSSLFQNLGVSVTPIQRIVLSGENGLKIDIHVIIPYEGLSGDQMGQIEKIFKVVYPVDDHHFKVILHYGTLVIDGVTPNMIDY
FGRPYEGIAVFDGKKITVTGTLWNGNKIIDERLINPDGSLLFRVTINGVTGWRLCERILAGPRSGS**LKGGAPTK**ASQEFMVSKGEELIKENMHMKLY
MEGTVNNHHFKCTSEGEGKPYEGTQTMRIKVVEGGPLPFAFDILATSFMYGSKTFINHTQGIPDFFKQSFPEGFTWERSTTYEDGGVLTATQDTSLQ
DGCLIYNVKIRGVNFPSNGPVMQKKTLGWEASTEMLYPADGGLEGRDYMALKLVGGGHLICNAKTTYRSKKPAKNLKMPGVYYVDRRLERIKEADKE
TSVEQHEVAVARYCDLPSKLGHKGT**DYKDHDGDYKDHDIDYKDDDDKD**I

*Highlights*:
Bold: His-tag
Green: mNG
Yellow: M$^{pro}$-N-ter-auto
Blue: NLuc
Grey: PL$^{pro}$-N-ter-auto
Red: mBeRFP
Bold: 3×FLAG-tag
Theoretical pI: 6.26
Theoretical MW: 81.3 kDa

**mNG-M$^{pro}$-Nter-auto-NLuc-PLpro-Nter-auto-LSS-mKate2:**

MGSS**HHHHHH**SSGLVPRGSHMVSKGEEDNMASLPATHELHIFGSINGVDFDMVGQGTGNPNDGYEELNLKSTKGDLQFSPWILVPHIGYGFHQYLPY
PDGMSPFQAAMVDGSGYQVHRTMQFEDGASLTVNYRYTYEGSHIKGEAQVKGTGFPADGPVMTNSLTAADWCRSKKTYPNDKTIISTFKWSYTTGNG
KRYRSTARTTYTFAKPMAANYLKNQPMYVFRKTELKHSKTELNFKEWQKAFTDVMGMDELYKAAATENLYAVLQSGFRGSGSAMVFTLEDFVGDWRQ
TAGYNLDQVLEQGGVSSLFQNLGVSVTPIQRIVLSGENGLKIDIHVIIPYEGLSGDQMGQIEKIFKVVYPVDDHHFKVILHYGTLVIDGVTPNMIDY
FGRPYEGIAVFDGKKITVTGTLWNGNKIIDERLINPDGSLLFRVTINGVTGWRLCERILAGPRSGSLKGGAPTKASQEFMVSKGEELIKENMHMKLY
MEGTVNNHHFKCTSEGEGKPYEGTQTMRIKVVEGGPLPFAFDILATSFMYGSYTFINHTQGIPDFFKQSFPEGFTWERVTTYEDGGVLTATQDTSLQ
DGCLIYNVKIRGVNFTSNGPVMQKKTLGWEAGTEMLYPADGGLEGRSDDALKLVGGGHLICNLKSTYRSKKPAKNLKVPGVYYVDRRLERIKEADKE
TYVEQHEVAVARYCDLPSKLGHKLNGT**DYKDHDGDYKDHDIDYKDDDDKD**I

*Highlights*:
Bold: His-tag
Green: mNG
Yellow: M$^{pro}$-N-ter-auto
Blue: NLuc
Grey: PL$^{pro}$-N-ter-auto
Red: LSS-mKate2
Bold: 3×FLAG-tag
Theoretical pI: 6.15
Theoretical MW: 81.5 kDa

**mNG-M$^{pro}$-Nter-auto-NLuc-PLpro-Nter-auto-LSS-CyOFP1:**

MGSS**HHHHHH**SSGLVPRGSHMVSKGEEDNMASLPATHELHIFGSINGVDFDMVGQGTGNPNDGYEELNLKSTKGDLQFSPWILVPHIGYGFHQYLPY
PDGMSPFQAAMVDGSGYQVHRTMQFEDGASLTVNYRYTYEGSHIKGEAQVKGTGFPADGPVMTNSLTAADWCRSKKTYPNDKTIISTFKWSYTTGNG
KRYRSTARTTYTFAKPMAANYLKNQPMYVFRKTELKHSKTELNFKEWQKAFTDVMGMDELYKAAATENLYAVLQSGFRGSGSAMVFTLEDFVGDWRQ
TAGYNLDQVLEQGGVSSLFQNLGVSVTPIQRIVLSGENGLKIDIHVIIPYEGLSGDQMGQIEKIFKVVYPVDDHHFKVILHYGTLVIDGVTPNMIDY
FGRPYEGIAVFDGKKITVTGTLWNGNKIIDERLINPDGSLLFRVTINGVTGWRLCERILAGPRSGSLKGGAPTKASQEFMVSKGEELIKENMRSKLY
LEGSVNGHQFKCTHEGEGKPYEGKQTNRIKVVEGGPLPFAFDILATHFMYGSKVFIKYPADLPDYFKQSFPEGFTWERVMVFEDGGVLTATQDTSLQ

DGELIYNVKVRGVNFPANGPVMQKKTLGWEPSTETMYPADGGLEGRCDKALKLVGGGHLHVNFKTTYKSKKPVKMPGVHYVDRRLERIKEADNETYV
EQYEHAVARYSNLGGGFTLEDFVGT**DYKDHDGDYKDHDIDYKDDDDKD**I

*Highlights*:
Bold: His-tag
Green: mNG
Yellow: $M^{pro}$-N-ter-auto
Blue: NLuc
Grey: $PL^{pro}$-N-ter-auto
Red: CyOFP1
Bold: 3×FLAG-tag
Theoretical pI: 6.09
Theoretical Mw: 81.4 kDa

**mNG-$M^{pro}$-Nter-auto-NLuc-PLpro-Nter-auto-mSca:**

MGSS**HHHHHH**SSGLVPRGSHMVSKGEEDNMASLPATHELHIFGSINGVDFDMVGQGTGNPNDGYEELNLKSTKGDLQFSPWILVPHIGYGFHQYLPY
PDGMSPFQAAMVDGSGYQVHRTMQFEDGASLTVNYRYTYEGSHIKGEAQVKGTGFPADGPVMTNSLTAADWCRSKKTYPNDKTIISTFKWSYTTGNG
KRYRSTARTTYTFAKPMAANYLKNQPMYVFRKTELKHSKTELNFKEWQKAFTDVMGMDELYKAAATENLYAVLQSGFRGSGSAMVFTLEDFVGDWRQ
TAGYNLDQVLEQGGVSSLFQNLGVSVTPIQRIVLSGENGLKIDIHVIIPYEGLSGDQMGQIEKIFKVVYPVDDHHFKVILHYGTLVIDGVTPNMIDY
FGRPYEGIAVFDGKKITVTGTLWNGNKIIDERLINPDGSLLFRVTINGVTGWRLCERILAGPRSGSLKGGAPTKASQEFMVSKGEAVIKEFMRFKVH
MEGSMNGHEFEIEGEGEGRPYEGTQTAKLKVTKGGPLPFSWDILSPQFMYGSRAFTKHPADIPDYYKQSFPEGFKWERVMNFEDGGAVTVTQDTSLE
DGTLIYKVKLRGTNFPPDGPVMQKKTMGWEASTERLYPEDGVLKGDIKMALRLKDGGRYLADFKTTYKAKKPVQMPGAYNVDRKLDITSHNEDYTVV
EQYERSEGRHSTGGMDELYKGT**DYKDHDGDYKDHDIDYKDDDDKD**I

*Highlights*:
Bold: His-tag
Green: mNG
Yellow: $M^{pro}$-N-ter-auto
Blue: NLuc
Grey: $PL^{pro}$-N-ter-auto
Red: mSca
Bold: 3×FLAG-tag
Theoretical pI: 5.99
Theoretical Mw: 81.2 kDa

**mNG-NSP4-5-Nter-auto-NLuc-NSP5-6-mSca:**

MGSS**HHHHHH**SSGLVPRGSHMVSKGEEDNMASLPATHELHIFGSINGVDFDMVGQGTGNPNDGYEELNLKSTKGDLQFSPWILVPHIGYGFHQYLPY
PDGMSPFQAAMVDGSGYQVHRTMQFEDGASLTVNYRYTYEGSHIKGEAQVKGTGFPADGPVMTNSLTAADWCRSKKTYPNDKTIISTFKWSYTTGNG
KRYRSTARTTYTFAKPMAANYLKNQPMYVFRKTELKHSKTELNFKEWQKAFTDVMGMDELYKAAATENLYAVLQSGFRGSGSAMVFTLEDFVGDWRQ
TAGYNLDQVLEQGGVSSLFQNLGVSVTPIQRIVLSGENGLKIDIHVIIPYEGLSGDQMGQIEKIFKVVYPVDDHHFKVILHYGTLVIDGVTPNMIDY
FGRPYEGIAVFDGKKITVTGTLWNGNKIIDERLINPDGSLLFRVTINGVTGWRLCERILAGPRSGSVTFQSAVKASQEFMVSKGEAVIKEFMRFKVH
MEGSMNGHEFEIEGEGEGRPYEGTQTAKLKVTKGGPLPFSWDILSPQFMYGSRAFTKHPADIPDYYKQSFPEGFKWERVMNFEDGGAVTVTQDTSLE
DGTLIYKVKLRGTNFPPDGPVMQKKTMGWEASTERLYPEDGVLKGDIKMALRLKDGGRYLADFKTTYKAKKPVQMPGAYNVDRKLDITSHNEDYTVV
EQYERSEGRHSTGGMDELYKGT**DYKDHDGDYKDHDIDYKDDDDKD**I

*Highlights*:
Bold: His-tag
Green: mNG
Yellow: NSP4-5 cleavage site
Blue: NLuc
Grey: NSP5-6 cleavage site
Red: mSca
Bold: 3×FLAG-tag
Theoretical pI: 5.94

Theoretical Mw: 81.3 kDa

**mNG-NSP4-5-Nter-auto-NLuc-NSP6-7- mSca:**

MGSS**HHHHHH**SSGLVPRGSHMVSKGEEDNMASLPATHELHIFGSINGVDFDMVGQGTGNPNDGYEELNLKSTKGDLQFSPWILVPHIGYGFHQYLPY
PDGMSPFQAAMVDGSGYQVHRTMQFEDGASLTVNYRYTYEGSHIKGEAQVKGTGFPADGPVMTNSLTAADWCRSKKTYPNDKTIISTFKWSYTTGNG
KRYRSTARTTYTFAKPMAANYLKNQPMYVFRKTELKHSKTELNFKEWQKAFTDVMGMDELYKAAATENLYAVLQSGFRGSGSAMVFTLEDFVGDWRQ
TAGYNLDQVLEQGGVSSLFQNLGVSVTPIQRIVLSGENGLKIDIHVIIPYEGLSGDQMGQIEKIFKVVYPVDDHHFKVILHYGTLVIDGVTPNMIDY
FGRPYEGIAVFDGKKITVTGTLWNGNKIIDERLINPDGSLLFRVTINGVTGWRLCERILAGPRSGSATVQSKMSASQEFMVSKGEAVIKEFMRFKVH
MEGSMNGHEFEIEGEGEGRPYEGTQTAKLKVTKGGPLPFSWDILSPQFMYGSRAFTKHPADIPDYYKQSFPEGFKWERVMNFEDGGAVTVTQDTSLE
DGTLIYKVKLRGTNFPPDGPVMQKKTMGWEASTERLYPEDGVLKGDIKMALRLKDGGRYLADFKTTYKAKKPVQMPGAYNVDRKLDITSHNEDYTVV
EQYERSEGRHSTGGMDELYKGT**DYKDHDGDYKDHDIDYKDDDDKD**I

*Highlights*:
Bold: His-tag
Green: mNG
Yellow: NSP4-5 cleavage site
Blue: NLuc
Grey: NSP6-7 cleavage site
Red: mSca
Bold: 3×FLAG-tag
Theoretical pI: 5.94
Theoretical Mw: 81.3 kDa

**mNG-NSP4-5-Nter-auto-NLuc-NSP7-8- mSca:**

MGSS**HHHHHH**SSGLVPRGSHMVSKGEEDNMASLPATHELHIFGSINGVDFDMVGQGTGNPNDGYEELNLKSTKGDLQFSPWILVPHIGYGFHQYLPY
PDGMSPFQAAMVDGSGYQVHRTMQFEDGASLTVNYRYTYEGSHIKGEAQVKGTGFPADGPVMTNSLTAADWCRSKKTYPNDKTIISTFKWSYTTGNG
KRYRSTARTTYTFAKPMAANYLKNQPMYVFRKTELKHSKTELNFKEWQKAFTDVMGMDELYKAAATENLYAVLQSGFRGSGSAMVFTLEDFVGDWRQ
TAGYNLDQVLEQGGVSSLFQNLGVSVTPIQRIVLSGENGLKIDIHVIIPYEGLSGDQMGQIEKIFKVVYPVDDHHFKVILHYGTLVIDGVTPNMIDY
FGRPYEGIAVFDGKKITVTGTLWNGNKIIDERLINPDGSLLFRVTINGVTGWRLCERILAGPRSGSATLQAIASASQEFMVSKGEAVIKEFMRFKVH
MEGSMNGHEFEIEGEGEGRPYEGTQTAKLKVTKGGPLPFSWDILSPQFMYGSRAFTKHPADIPDYYKQSFPEGFKWERVMNFEDGGAVTVTQDTSLE
DGTLIYKVKLRGTNFPPDGPVMQKKTMGWEASTERLYPEDGVLKGDIKMALRLKDGGRYLADFKTTYKAKKPVQMPGAYNVDRKLDITSHNEDYTVV
EQYERSEGRHSTGGMDELYKGT**DYKDHDGDYKDHDIDYKDDDDKD**I

*Highlights*:
Bold: His-tag
Green: mNG
Yellow: NSP4-5 cleavage site
Blue: NLuc
Grey: NSP7-8 cleavage site
Red: mSca
Bold: 3×FLAG-tag
Theoretical pI: 5.89
Theoretical Mw: 81.2 kDa

**mNG-NSP4-5-Nter-auto-NLuc-NSP8-9- mSca:**

MGSS**HHHHHH**SSGLVPRGSHMVSKGEEDNMASLPATHELHIFGSINGVDFDMVGQGTGNPNDGYEELNLKSTKGDLQFSPWILVPHIGYGFHQYLPY
PDGMSPFQAAMVDGSGYQVHRTMQFEDGASLTVNYRYTYEGSHIKGEAQVKGTGFPADGPVMTNSLTAADWCRSKKTYPNDKTIISTFKWSYTTGNG
KRYRSTARTTYTFAKPMAANYLKNQPMYVFRKTELKHSKTELNFKEWQKAFTDVMGMDELYKAAATENLYAVLQSGFRGSGSAMVFTLEDFVGDWRQ
TAGYNLDQVLEQGGVSSLFQNLGVSVTPIQRIVLSGENGLKIDIHVIIPYEGLSGDQMGQIEKIFKVVYPVDDHHFKVILHYGTLVIDGVTPNMIDY
FGRPYEGIAVFDGKKITVTGTLWNGNKIIDERLINPDGSLLFRVTINGVTGWRLCERILAGPRSGSVKLQNNELASQEFMVSKGEAVIKEFMRFKVH
MEGSMNGHEFEIEGEGEGRPYEGTQTAKLKVTKGGPLPFSWDILSPQFMYGSRAFTKHPADIPDYYKQSFPEGFKWERVMNFEDGGAVTVTQDTSLE
DGTLIYKVKLRGTNFPPDGPVMQKKTMGWEASTERLYPEDGVLKGDIKMALRLKDGGRYLADFKTTYKAKKPVQMPGAYNVDRKLDITSHNEDYTVV
EQYERSEGRHSTGGMDELYKGT**DYKDHDGDYKDHDIDYKDDDDKD**I

*Highlights*:
Bold: His-tag
Green: mNG
Yellow: NSP4-5 cleavage site
Blue: NLuc
Grey: NSP8-9 cleavage site
Red: mSca
Bold: 3×FLAG-tag
Theoretical pI: 5.89
Theoretical Mw: 81.4 kDa

**mNG-NSP4-5-Nter-auto-NLuc-NSP9-10-mSca:**

MGSS**HHHHHH**SSGLVPRGSHMVSKGEEDNMASLPATHELHIFGSINGVDFDMVGQGTGNPNDGYEELNLKSTKGDLQFSPWILVPHIGYGFHQYLPY
PDGMSPFQAAMVDGSGYQVHRTMQFEDGASLTVNYRYTYEGSHIKGEAQVKGTGFPADGPVMTNSLTAADWCRSKKTYPNDKTIISTFKWSYTTGNG
KRYRSTARTTYTFAKPMAANYLKNQPMYVFRKTELKHSKTELNFKEWQKAFTDVMGMDELYKAAATENLYAVLQSGFRGSGSAMVFTLEDFVGDWRQ
TAGYNLDQVLEQGGVSSLFQNLGVSVTPIQRIVLSGENGLKIDIHVIIPYEGLSGDQMGQIEKIFKVVYPVDDHHFKVILHYGTLVIDGVTPNMIDY
FGRPYEGIAVFDGKKITVTGTLWNGNKIIDERLINPDGSLLFRVTINGVTGWRLCERILAGPRSGSVRLQAGNAASQEFMVSKGEAVIKEFMRFKVH
MEGSMNGHEFEIEGEGEGRPYEGTQTAKLKVTKGGPLPFSWDILSPQFMYGSRAFTKHPADIPDYYKQSFPEGFKWERVMNFEDGGAVTVTQDTSLE
DGTLIYKVKLRGTNFPPDGPVMQKKTMGWEASTERLYPEDGVLKGDIKMALRLKDGGRYLADFKTTYKAKKPVQMPGAYNVDRKLDITSHNEDYTVV
EQYERSEGRHSTGGMDELYKGT**DYKDHDGDYKDHDIDYKDDDDKD**I

*Highlights*:
Bold: His-tag
Green: mNG
Yellow: NSP4-5 cleavage site
Blue: NLuc
Grey: NSP9-10 cleavage site
Red: mSca
Bold: 3×FLAG-tag
Theoretical pI: 5.94
Theoretical Mw: 81.3 kDa

**mNG-NSP4-5-Nter-auto-NLuc-NSP10-11-12- mSca:**

MGSS**HHHHHH**SSGLVPRGSHMVSKGEEDNMASLPATHELHIFGSINGVDFDMVGQGTGNPNDGYEELNLKSTKGDLQFSPWILVPHIGYGFHQYLPY
PDGMSPFQAAMVDGSGYQVHRTMQFEDGASLTVNYRYTYEGSHIKGEAQVKGTGFPADGPVMTNSLTAADWCRSKKTYPNDKTIISTFKWSYTTGNG
KRYRSTARTTYTFAKPMAANYLKNQPMYVFRKTELKHSKTELNFKEWQKAFTDVMGMDELYKAAATENLYAVLQSGFRGSGSAMVFTLEDFVGDWRQ
TAGYNLDQVLEQGGVSSLFQNLGVSVTPIQRIVLSGENGLKIDIHVIIPYEGLSGDQMGQIEKIFKVVYPVDDHHFKVILHYGTLVIDGVTPNMIDY
FGRPYEGIAVFDGKKITVTGTLWNGNKIIDERLINPDGSLLFRVTINGVTGWRLCERILAGPRSGSPMLQSADAASQEFMVSKGEAVIKEFMRFKVH
MEGSMNGHEFEIEGEGEGRPYEGTQTAKLKVTKGGPLPFSWDILSPQFMYGSRAFTKHPADIPDYYKQSFPEGFKWERVMNFEDGGAVTVTQDTSLE
DGTLIYKVKLRGTNFPPDGPVMQKKTMGWEASTERLYPEDGVLKGDIKMALRLKDGGRYLADFKTTYKAKKPVQMPGAYNVDRKLDITSHNEDYTVV
EQYERSEGRHSTGGMDELYKGT**DYKDHDGDYKDHDIDYKDDDDKD**I

*Highlights*:
Bold: His-tag
Green: mNG
Yellow: NSP4-5 cleavage site
Blue: NLuc
Grey: NSP10-11-12 cleavage site
Red: mSca
Bold: 3×FLAG-tag
Theoretical pI: 5.84
Theoretical Mw: 81.3 kDa

**mNG-NSP4-5-Nter-auto-NLuc-NSP12-13- mSca:**

MGSS**HHHHHH**SSGLVPRGSHMVSKGEEDNMASLPATHELHIFGSINGVDFDMVGQGTGNPNDGYEELNLKSTKGDLQFSPWILVPHIGYGFHQYLPY
PDGMSPFQAAMVDGSGYQVHRTMQFEDGASLTVNYRYTYEGSHIKGEAQVKGTGFPADGPVMTNSLTAADWCRSKKTYPNDKTIISTFKWSYTTGNG
KRYRSTARTTYTFAKPMAANYLKNQPMYVFRKTELKHSKTELNFKEWQKAFTDVMGMDELYKAAATENLYAVLQSGFRGSGSAMVFTLEDFVGDWRQ
TAGYNLDQVLEQGGVSSLFQNLGVSVTPIQRIVLSGENGLKIDIHVIIPYEGLSGDQMGQIEKIFKVVYPVDDHHFKVILHYGTLVIDGVTPNMIDY
FGRPYEGIAVFDGKKITVTGTLWNGNKIIDERLINPDGSLLFRVTINGVTGWRLCERILAGPRSGSTVLQAVGAASQEFMVSKGEAVIKEFMRFKVH
MEGSMNGHEFEIEGEGEGRPYEGTQTAKLKVTKGGPLPFSWDILSPQFMYGSRAFTKHPADIPDYYKQSFPEGFKWERVMNFEDGGAVTVTQDTSLE
DGTLIYKVKLRGTNFPPDGPVMQKKTMGWEASTERLYPEDGVLKGDIKMALRLKDGGRYLADFKTTYKAKKPVQMPGAYNVDRKLDITSHNEDYTVV
EQYERSEGRHSTGGMDELYKGT**DYKDHDGDYKDHDIDYKDDDDKD**I

*Highlights*:
Bold: His-tag
Green: mNG
Yellow: NSP4-5 cleavage site
Blue: NLuc
Grey: NSP12-13 cleavage site
Red: mSca
Bold: 3×FLAG-tag
Theoretical pI: 5.89
Theoretical Mw: 81.2 kDa

**mNG-NSP4-5-Nter-auto-NLuc-NSP13-14- mSca:**

MGSS**HHHHHH**SSGLVPRGSHMVSKGEEDNMASLPATHELHIFGSINGVDFDMVGQGTGNPNDGYEELNLKSTKGDLQFSPWILVPHIGYGFHQYLPY
PDGMSPFQAAMVDGSGYQVHRTMQFEDGASLTVNYRYTYEGSHIKGEAQVKGTGFPADGPVMTNSLTAADWCRSKKTYPNDKTIISTFKWSYTTGNG
KRYRSTARTTYTFAKPMAANYLKNQPMYVFRKTELKHSKTELNFKEWQKAFTDVMGMDELYKAAATENLYAVLQSGFRGSGSAMVFTLEDFVGDWRQ
TAGYNLDQVLEQGGVSSLFQNLGVSVTPIQRIVLSGENGLKIDIHVIIPYEGLSGDQMGQIEKIFKVVYPVDDHHFKVILHYGTLVIDGVTPNMIDY
FGRPYEGIAVFDGKKITVTGTLWNGNKIIDERLINPDGSLLFRVTINGVTGWRLCERILAGPRSGSATLQAENVASQEFMVSKGEAVIKEFMRFKVH
MEGSMNGHEFEIEGEGEGRPYEGTQTAKLKVTKGGPLPFSWDILSPQFMYGSRAFTKHPADIPDYYKQSFPEGFKWERVMNFEDGGAVTVTQDTSLE
DGTLIYKVKLRGTNFPPDGPVMQKKTMGWEASTERLYPEDGVLKGDIKMALRLKDGGRYLADFKTTYKAKKPVQMPGAYNVDRKLDITSHNEDYTVV
EQYERSEGRHSTGGMDELYKGT**DYKDHDGDYKDHDIDYKDDDDKD**I

*Highlights*:
Bold: His-tag
Green: mNG
Yellow: NSP4-5 cleavage site
Blue: NLuc
Grey: NSP13-14 cleavage site
Red: mSca
Bold: 3×FLAG-tag
Theoretical pI: 5.84
Theoretical Mw: 81.3 kDa

**mNG-NSP4-5-Nter-auto-NLuc-NSP14-15- mSca:**

MGSS**HHHHHH**SSGLVPRGSHMVSKGEEDNMASLPATHELHIFGSINGVDFDMVGQGTGNPNDGYEELNLKSTKGDLQFSPWILVPHIGYGFHQYLPY
PDGMSPFQAAMVDGSGYQVHRTMQFEDGASLTVNYRYTYEGSHIKGEAQVKGTGFPADGPVMTNSLTAADWCRSKKTYPNDKTIISTFKWSYTTGNG
KRYRSTARTTYTFAKPMAANYLKNQPMYVFRKTELKHSKTELNFKEWQKAFTDVMGMDELYKAAATENLYAVLQSGFRGSGSAMVFTLEDFVGDWRQ
TAGYNLDQVLEQGGVSSLFQNLGVSVTPIQRIVLSGENGLKIDIHVIIPYEGLSGDQMGQIEKIFKVVYPVDDHHFKVILHYGTLVIDGVTPNMIDY
FGRPYEGIAVFDGKKITVTGTLWNGNKIIDERLINPDGSLLFRVTINGVTGWRLCERILAGPRSGSTRLQSLENASQEFMVSKGEAVIKEFMRFKVH
MEGSMNGHEFEIEGEGEGRPYEGTQTAKLKVTKGGPLPFSWDILSPQFMYGSRAFTKHPADIPDYYKQSFPEGFKWERVMNFEDGGAVTVTQDTSLE
DGTLIYKVKLRGTNFPPDGPVMQKKTMGWEASTERLYPEDGVLKGDIKMALRLKDGGRYLADFKTTYKAKKPVQMPGAYNVDRKLDITSHNEDYTVV
EQYERSEGRHSTGGMDELYKGT**DYKDHDGDYKDHDIDYKDDDDKD**I

*Highlights*:
Bold: His-tag
Green: mNG
Yellow: NSP4-5 cleavage site
Blue: NLuc
Grey: NSP14-15 cleavage site
Red: mSca
Bold: 3×FLAG-tag
Theoretical pI: 5.89
Theoretical Mw: 81.4 kDa

**mNG-NSP4-5-Nter-auto-NLuc-NSP15-16-mSca:**

MGSS**HHHHHH**SSGLVPRGSHMVSKGEEDNMASLPATHELHIFGSINGVDFDMVGQGTGNPNDGYEELNLKSTKGDLQFSPWILVPHIGYGFHQYLPY
PDGMSPFQAAMVDGSGYQVHRTMQFEDGASLTVNYRYTYEGSHIKGEAQVKGTGFPADGPVMTNSLTAADWCRSKKTYPNDKTIISTFKWSYTTGNG
KRYRSTARTTYTFAKPMAANYLKNQPMYVFRKTELKHSKTELNFKEWQKAFTDVMGMDELYKAAATENLYAVLQSGFRGSGSAMVFTLEDFVGDWRQ
TAGYNLDQVLEQGGVSSLFQNLGVSVTPIQRIVLSGENGLKIDIHVIIPYEGLSGDQMGQIEKIFKVVYPVDDHHFKVILHYGTLVIDGVTPNMIDY
FGRPYEGIAVFDGKKITVTGTLWNGNKIIDERLINPDGSLLFRVTINGVTGWRLCERILAGPRSGSPKLQSSQAASQEFMVSKGEAVIKEFMRFKVH
MEGSMNGHEFEIEGEGEGRPYEGTQTAKLKVTKGGPLPFSWDILSPQFMYGSRAFTKHPADIPDYYKQSFPEGFKWERVMNFEDGGAVTVTQDTSLE
DGTLIYKVKLRGTNFPPDGPVMQKKTMGWEASTERLYPEDGVLKGDIKMALRLKDGGRYLADFKTTYKAKKPVQMPGAYNVDRKLDITSHNEDYTVV
EQYERSEGRHSTGGMDELYKGT**DYKDHDGDYKDHDIDYKDDDDKD**I

*Highlights*:
Bold: His-tag
Green: mNG
Yellow: NSP4-5 cleavage site
Blue: NLuc
Grey: NSP15-16 cleavage site
Red: mSca
Bold: 3×FLAG-tag
Theoretical pI: 5.94
Theoretical Mw: 81.3 kDa

**mNG-NSP1-2-Nter-auto-NLuc-NSP2-3- mSca:**

MGSS**HHHHHH**SSGLVPRGSHMVSKGEEDNMASLPATHELHIFGSINGVDFDMVGQGTGNPNDGYEELNLKSTKGDLQFSPWILVPHIGYGFHQYLPY
PDGMSPFQAAMVDGSGYQVHRTMQFEDGASLTVNYRYTYEGSHIKGEAQVKGTGFPADGPVMTNSLTAADWCRSKKTYPNDKTIISTFKWSYTTGNG
KRYRSTARTTYTFAKPMAANYLKNQPMYVFRKTELKHSKTELNFKEWQKAFTDVMGMDELYKAAATENLYLNGGAYTRGSGSAMVFTLEDFVGDWRQ
TAGYNLDQVLEQGGVSSLFQNLGVSVTPIQRIVLSGENGLKIDIHVIIPYEGLSGDQMGQIEKIFKVVYPVDDHHFKVILHYGTLVIDGVTPNMIDY
FGRPYEGIAVFDGKKITVTGTLWNGNKIIDERLINPDGSLLFRVTINGVTGWRLCERILAGPRSGSLKGGAPTKASQEFMVSKGEAVIKEFMRFKVH
MEGSMNGHEFEIEGEGEGRPYEGTQTAKLKVTKGGPLPFSWDILSPQFMYGSRAFTKHPADIPDYYKQSFPEGFKWERVMNFEDGGAVTVTQDTSLE
DGTLIYKVKLRGTNFPPDGPVMQKKTMGWEASTERLYPEDGVLKGDIKMALRLKDGGRYLADFKTTYKAKKPVQMPGAYNVDRKLDITSHNEDYTVV
EQYERSEGRHSTGGMDELYKGT**DYKDHDGDYKDHDIDYKDDDDKD**I

*Highlights*:
Bold: His-tag
Green: mNG
Yellow: NSP1-2 cleavage site
Blue: NLuc
Grey: NSP2-3 cleavage site
Red: mSca
Bold: 3×FLAG-tag
Theoretical pI: 5.99
Theoretical Mw: 81.2 kDa

**mNG-NSP3-4-Nter-auto-NLuc-NSP2-3-mSca:**

MGSS**HHHHHH**SSGLVPRGSHMVSKGEEDNMASLPATHELHIFGSINGVDFDMVGQGTGNPNDGYEELNLKSTKGDLQFSPWILVPHIGYGFHQYLPY
PDGMSPFQAAMVDGSGYQVHRTMQFEDGASLTVNYRYTYEGSHIKGEAQVKGTGFPADGPVMTNSLTAADWCRSKKTYPNDKTIISTFKWSYTTGNG
KRYRSTARTTYTFAKPMAANYLKNQPMYVFRKTELKHSKTELNFKEWQKAFTDVMGMDELYKAAATENLYLKGGKIVNGSGSAMVFTLEDFVGDWRQ
TAGYNLDQVLEQGGVSSLFQNLGVSVTPIQRIVLSGENGLKIDIHVIIPYEGLSGDQMGQIEKIFKVVYPVDDHHFKVILHYGTLVIDGVTPNMIDY
FGRPYEGIAVFDGKKITVTGTLWNGNKIIDERLINPDGSLLFRVTINGVTGWRLCERILAGPRSGSLKGGAPTKASQEFMVSKGEAVIKEFMRFKVH
MEGSMNGHEFEIEGEGEGRPYEGTQTAKLKVTKGGPLPFSWDILSPQFMYGSRAFTKHPADIPDYYKQSFPEGFKWERVMNFEDGGAVTVTQDTSLE
DGTLIYKVKLRGTNFPPDGPVMQKKTMGWEASTERLYPEDGVLKGDIKMALRLKDGGRYLADFKTTYKAKKPVQMPGAYNVDRKLDITSHNEDYTVV
EQYERSEGRHSTGGMDELYKGT**DYKDHDGDYKDHDIDYKDDDDKD**I

*Highlights*:
Bold: His-tag
Green: mNG
Yellow: NSP3-4 cleavage site
Blue: NLuc
Grey: NSP2-3 cleavage site
Red: mSca
Bold: 3×FLAG-tag
Theoretical pI: 6.05
Theoretical Mw: 81.2 kDa

### Supporting Figures

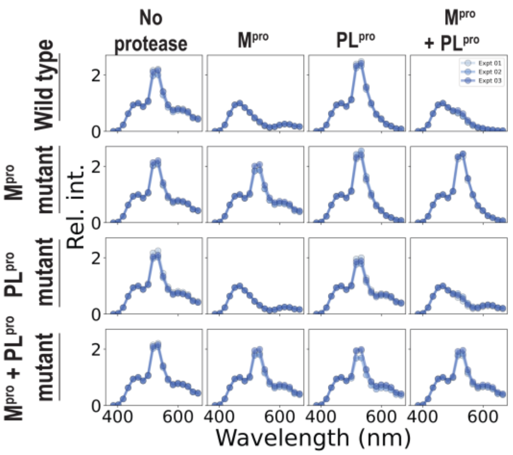


**Supporting Figure 1. Mutations in SARS-CoV-2 M$^{pro}$ and PL$^{pro}$ cleavage sites abrogate DuProSense biosensor BRET response.**
Graphs showing bioluminescence spectra of cells expressing DuProSense biosensor wild type and the mutants in the absence and presence of the proteases, M$^{pro}$ and PL$^{pro}$. Note the cleavage of the sensors in wild type as evidenced by the reduction in mNG (533 nm) and mSca (615 nm) peaks in the presence of M$^{pro}$ and PL$^{pro}$, respectively whereas no cleavage was observed in the mutants in the presence of respective proteases. Data shown are mean from three independent experiments, with each experiment performed in triplicates.

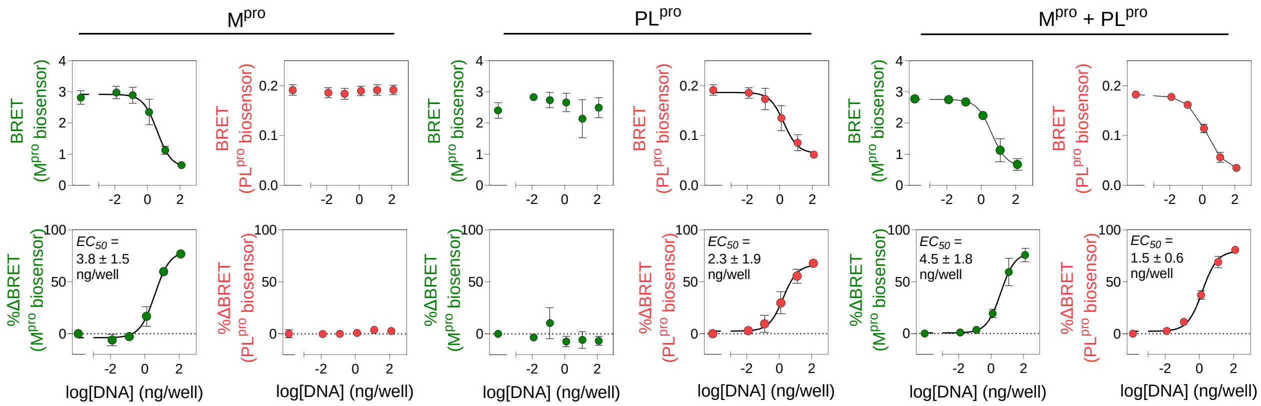


*Supporting Figure 2. Individual and simultaneous M$^{pro}$ and PL$^{pro}$ plasmid DNA concentration-dependent cleavage of the mSca-containing DuProSense biosensor.*

Graphs showing BRET ratio (upper panel) and percentage change in BRET (lower panel) in the green and the red channel BRET measured from HEK293T cells transfected with DuProSense biosensor either individually or simultaneously with M$^{pro}$ and PL$^{pro}$ plasmid DNA. Data shown are mean ± S.D. from two independent experiments, with each experiment performed in triplicates.

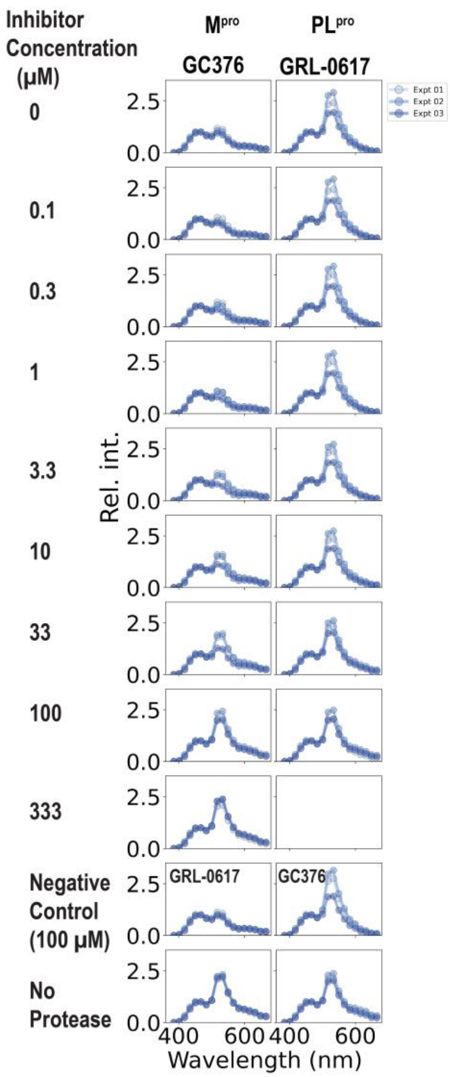


**Supporting Figure 3. Bioluminescence spectra of mSca-containing DuProSense biosensor showing highly specific pharmacological inhibition of $M^{pro}$ and $PL^{pro}$ in living cells.**

Graphs showing the bioluminescence spectra measured from living HEK293T cells expressing mSca-containing DuProSense biosensor in the presence of $M^{pro}$ and GC376 (left panel) and $PL^{pro}$ and GRL-0617 (right panel). Note the dose-dependent inhibition of proteases which is evident from the increase in mNG peak intensity (533 nm; left panel) and mSca peak intensity (615 nm; right panel). Data shown are mean from three independent experiments, with each experiment performed in triplicates.

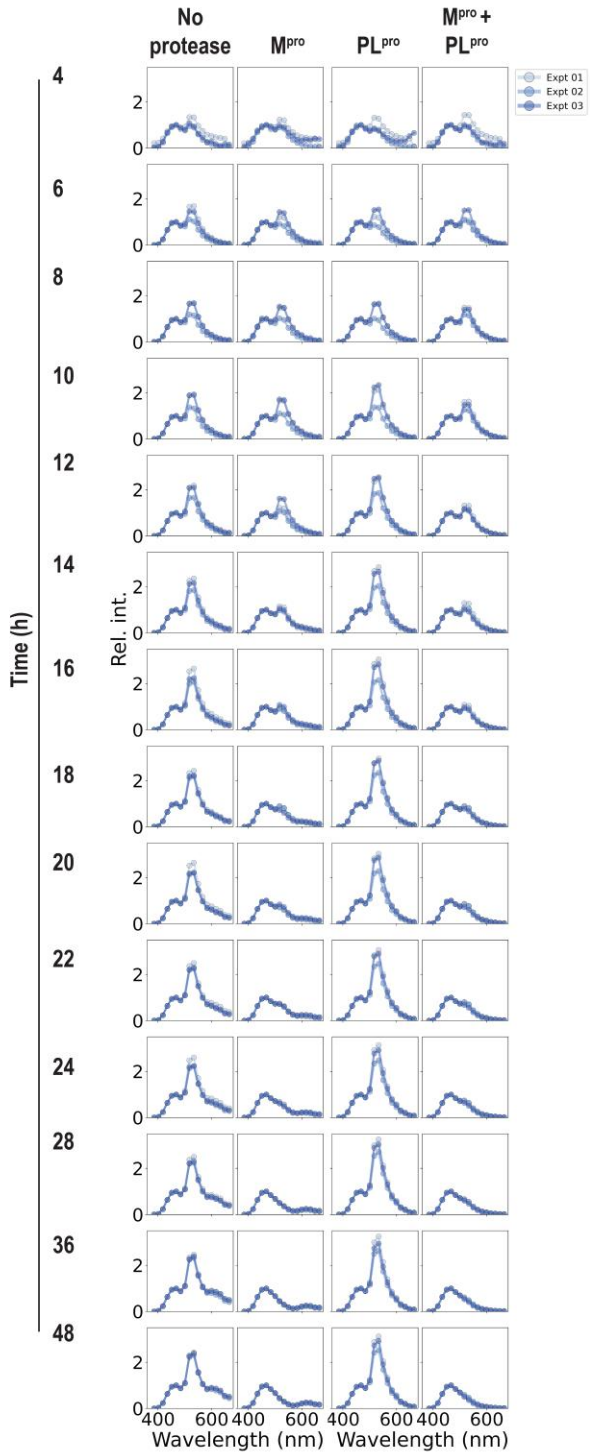


**Supporting Figure 4. Bioluminescence spectra – mSca-containing DuProSense biosensor enables simultaneous monitoring of $M^{pro}$ and $PL^{pro}$ cleavage activity in living cells.**

Graphs showing the bioluminescence spectra measured from living HEK293T cells expressing DuProSense biosensor cleavage in the absence or presence of either $M^{pro}$ or $PL^{pro}$ or both. Data shown are mean from three independent experiments, with each experiment performed in triplicates.

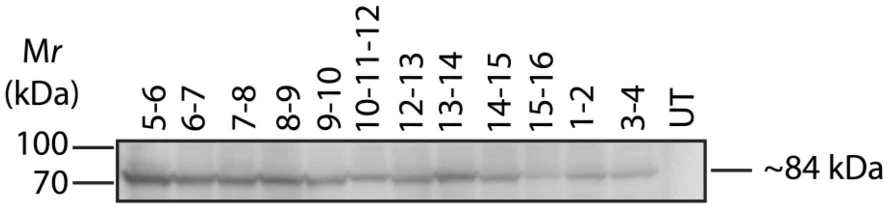


**Supporting Figure 5. Expression of various DuProSense biosensors.**

Western blot performed using anti-FLAG tag antibody showing the expression of various DuProSense biosensors in the lysate from HEK293T cells. UT, untransfected control.

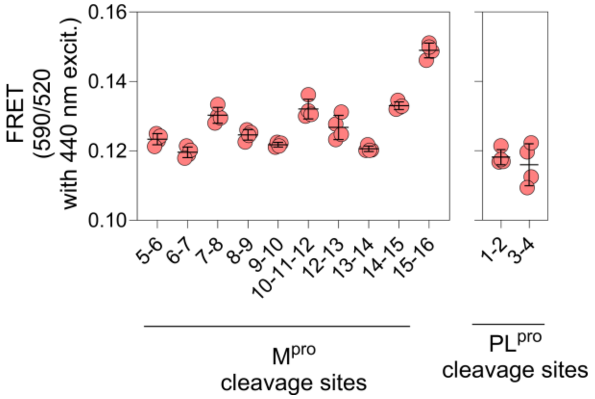


**Supporting Figure 6. FRET ratio of various $M^{pro}$ and $PL^{pro}$ DuProSense biosensor constructs.**

Graph showing the FRET ratio (590/520) measured from cell lysates of HEK293T cells expressing various Mpro and PLpro DuProSense biosensor constructs when excited at 440 nm. Left panel, DuProSense biosensor containing $M^{pro}$ cleavage sites; right panel, DuProSense biosensor containing $PL^{pro}$ cleavage sites. Data shown are mean ± S.D. from four independent experiments.

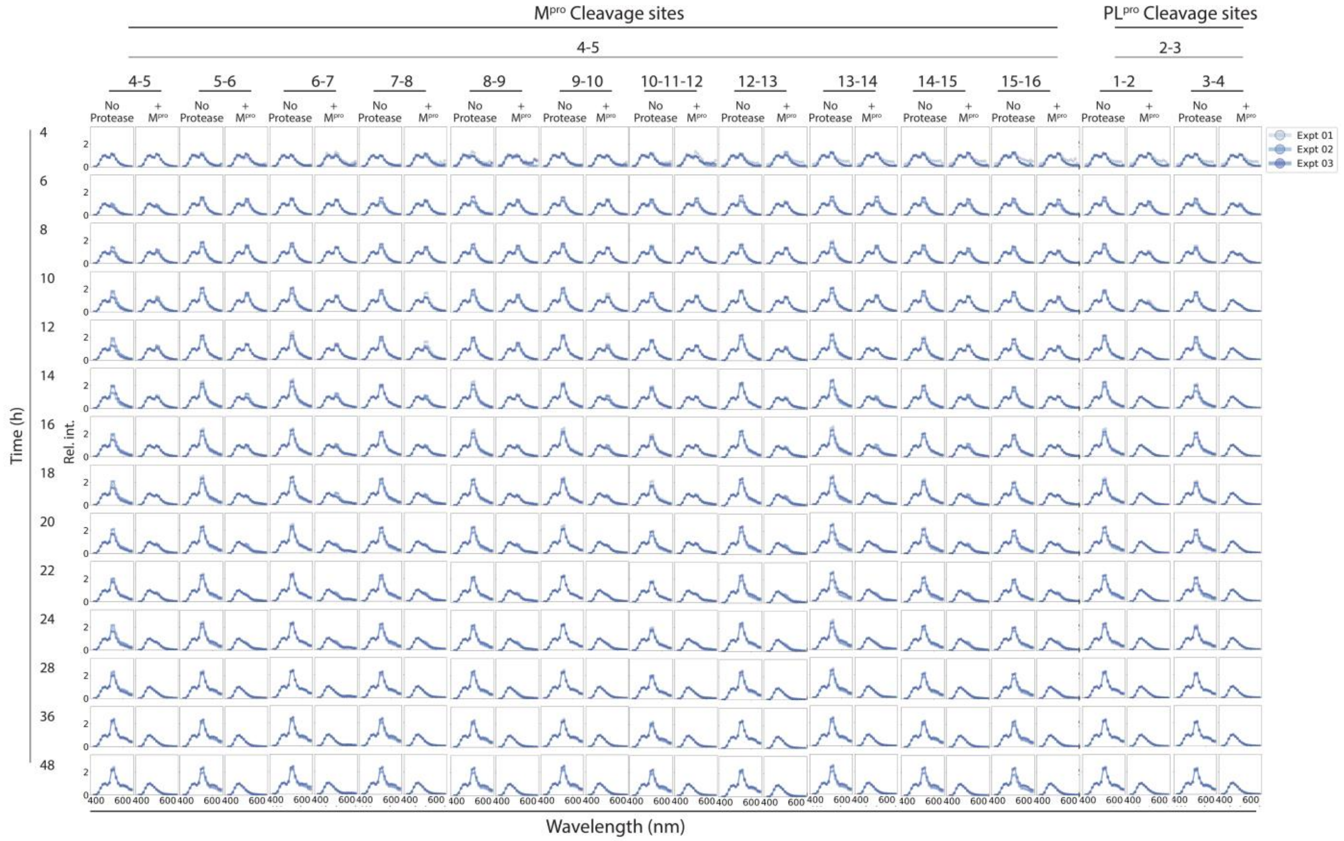


**Supporting Figure 7. Bioluminescence spectra – DuProSense biosensor platform reveals differential cleavage kinetics of M$^{pro}$ cleavage sites in living cells.**

Graphs showing the bioluminescence spectra measured from living HEK293T cells expressing DuProSense biosensors containing M$^{pro}$ and PL$^{pro}$ cleavage sites at indicated time points. Note the time-dependent cleavage of corresponding M$^{pro}$ or PL$^{pro}$ DuProSense biosensors in the presence of M$^{pro}$ or PL$^{pro}$, respectively as reflected by the reduction in the mNG (533 nm) and mSca (615 nm) peaks in the bioluminescence spectra. Data shown are mean from three independent experiments, with each experiment performed in triplicates.

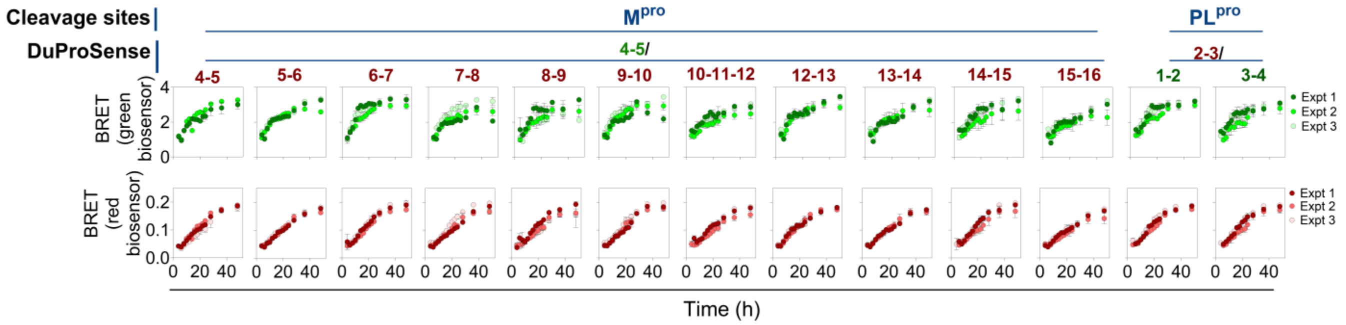


**Supporting Figure 8. BRET kinetics of DuProSense biosensor in the absence of proteases in live cells.**

Graphs showing the BRET in green (upper panel) and red (lower panel) channels measured from living HEK293T cells expressing DuProSense biosensors containing M$^{pro}$ and PL$^{pro}$ cleavage sites at indicated time points in the absence of $M^{pro}$ and $PL^{pro}$. Data shown are mean ± S.D. from three independent experiments, with each experiment performed in triplicates.

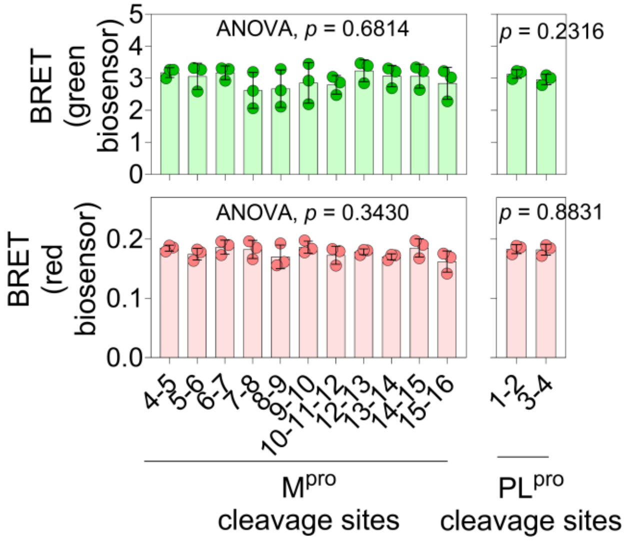


***Supporting Figure 9. BRET efficiency of* various $M^{pro}$ and $PL^{pro}$ DuProSense biosensor constructs**. Graphs showing the BRET ratio of each DuProSense biosensor in green and red channels measured from living HEK293T cells expressing DuProSense biosensors containing $M^{pro}$ and $PL^{pro}$ cleavage sites in the absence of proteases at 48 h post-transfection. Data shown are mean ± S.D. from three independent experiments, with each experiment performed in triplicates.

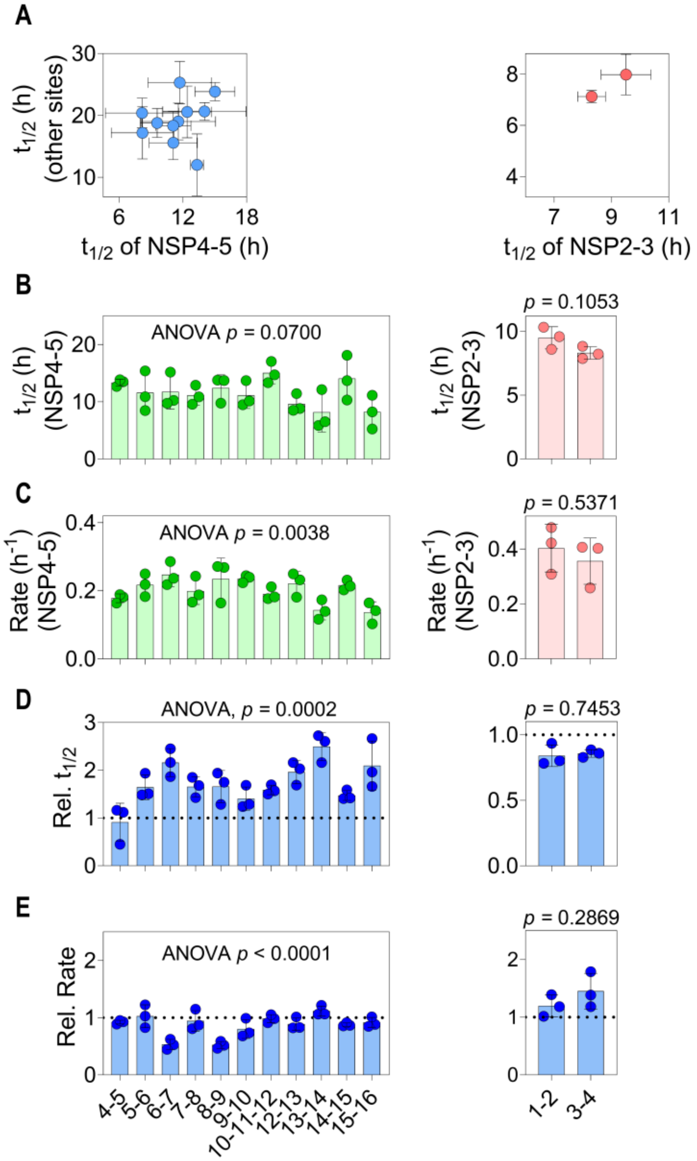


**Supporting Figure 10. Comparison of the cleavage kinetics of $M^{pro}$ cleavage sites obtained from live cell assay.**

(A) Graph showing the $t_{1/2}$ of all $M^{pro}$ and $PL^{pro}$ cleavage sites compared to NSP4-5 (right panel) and NSP2-3 (left panel), respectively obtained from the kinetic sigmoidal model fitted the data obtained from living HEK293T cells expressing DuProSense biosensors containing $M^{pro}$ and $PL^{pro}$ cleavage sites in the presence of $M^{pro}$ and $PL^{pro}$.

(B, C) Graphs showing $t_{1/2}$ (B) and cleavage rate (C) of $M^{pro}$ cleavage site NSP4-5 and $PL^{pro}$ cleavage site NSP2-3 obtained from the kinetic sigmoidal model fitted the data obtained from living HEK293T cells expressing DuProSense biosensors containing $M^{pro}$ and $PL^{pro}$ cleavage sites in the presence of $M^{pro}$ and $PL^{pro}$.

(D,E) Graphs showing the relative $t_{1/2}$ (D) and rate (E) of all $M^{pro}$ (left panel) and $PL^{pro}$ (right panel) cleavage sites relative to NSP4-5 and NSP2-3 cleavage sites, respectively, obtained from the kinetic sigmoidal model fitted the data obtained from living HEK293T cells expressing DuProSense biosensors containing $M^{pro}$ and $PL^{pro}$ cleavage sites in the presence of $M^{pro}$ and $PL^{pro}$.

Data shown are mean ± S.D. from three independent experiments, with each experiment performed in triplicates.

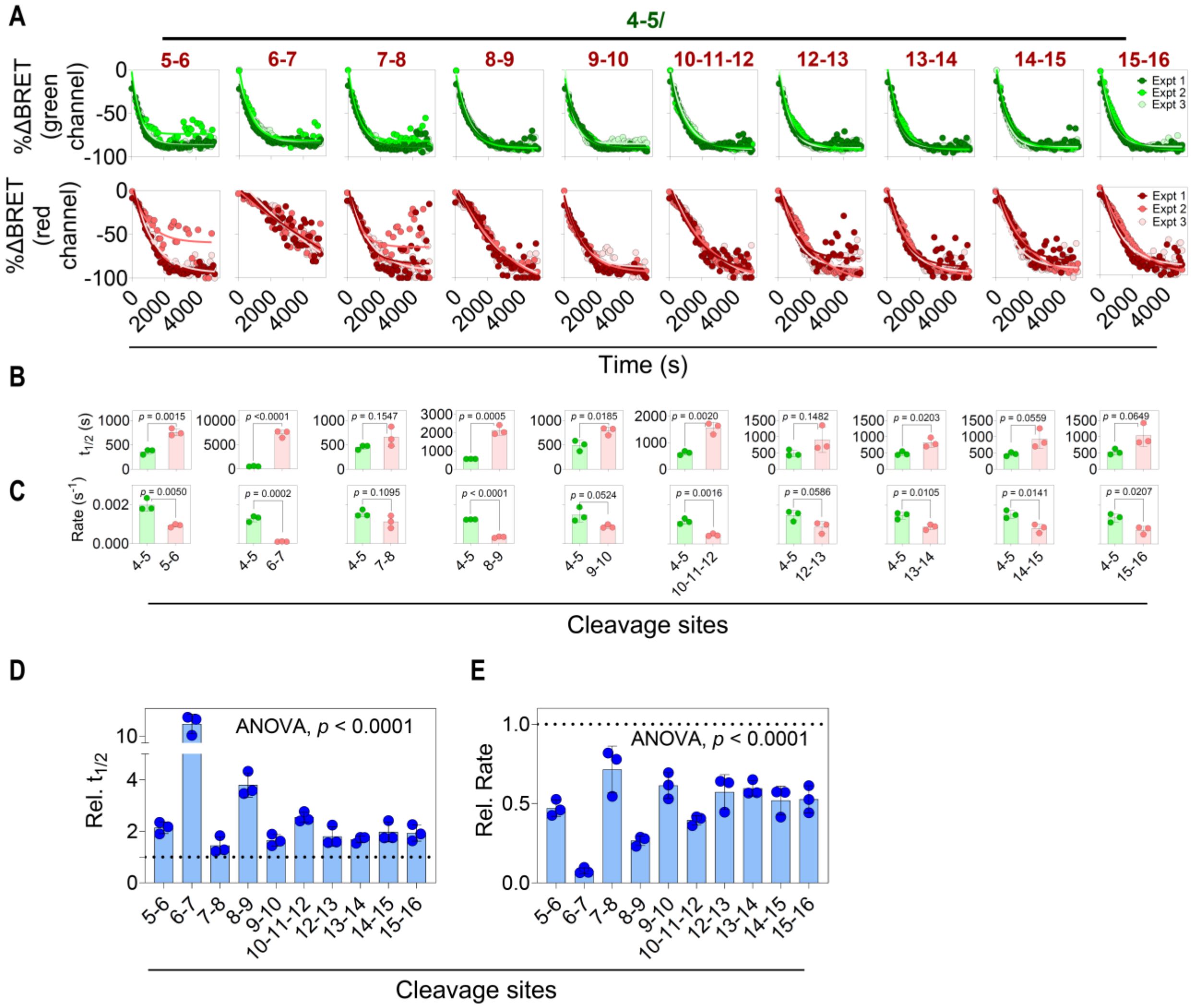


*Supporting Figure 11. In vitro cleavage kinetics of various DuProSense biosensors with SARS-CoV-2 $M^{pro}$.*

(A) Graphs showing the percentage change in BRET of DuProSense biosensor containing $M^{pro}$ cleavage sites with $M^{pro}$ measured from cell lysates of HEK293T cells expressing DuProSense biosensors containing $M^{pro}$ cleavage sites incubated with purified $M^{pro}$.
(B, C) Graphs showing the half-time ($t_{1/2}$) (B) and cleavage rates (C) of all $M^{pro}$ cleavage sites obtained from the kinetic sigmoidal model fitted the data obtained from in vitro assay using lysate of cells expressing DuProSense biosensors containing $M^{pro}$ and purified $M^{pro}$.
(D, E) Graphs showing the relative $t_{1/2}$ (D) and relative rate (E) of all cleavage sites of $M^{pro}$.
Data shown are mean ± S.D. from three independent experiments.

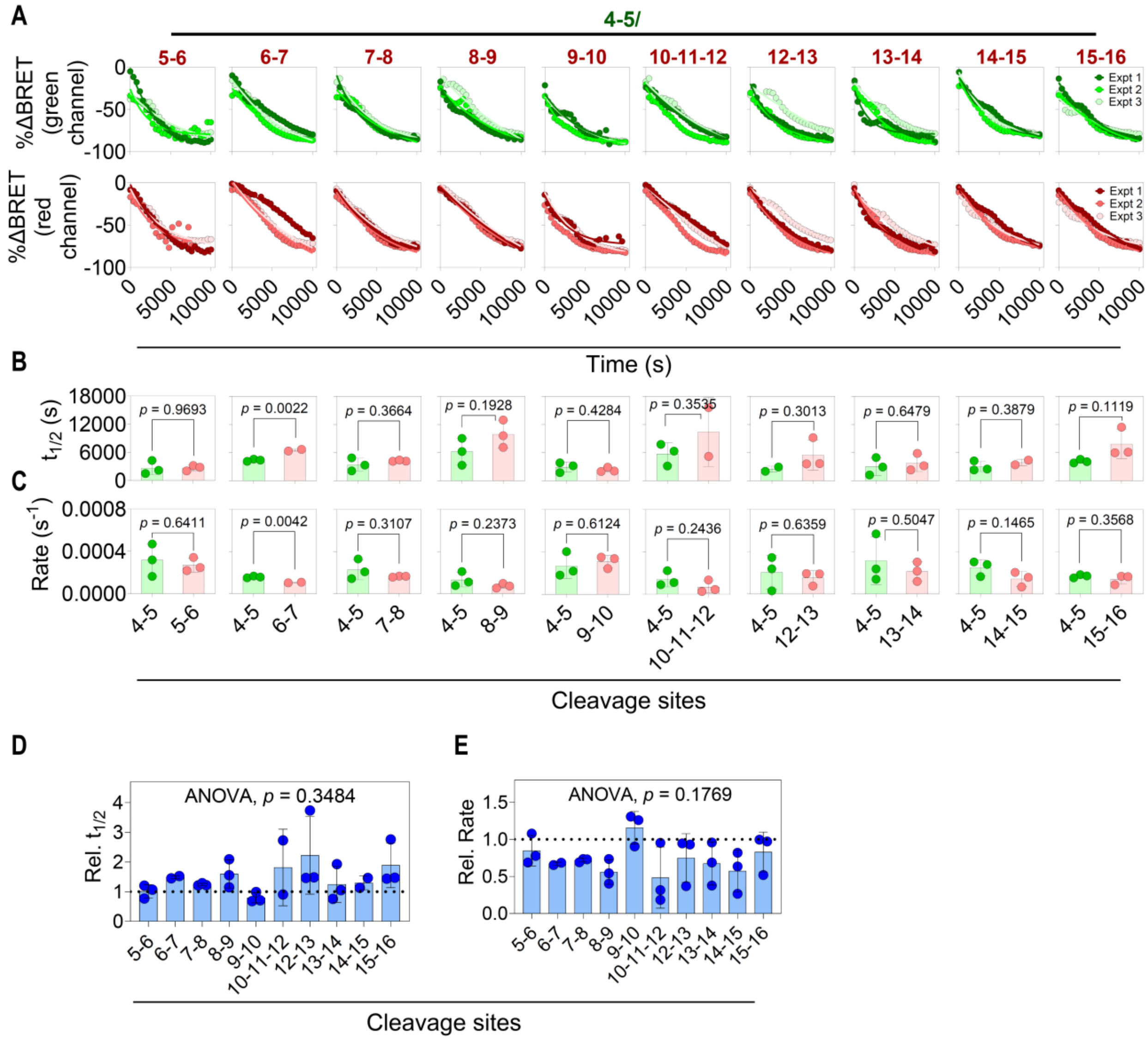


**Supporting Figure 12. In vitro cleavage kinetics of various DuProSense biosensors with SARS-CoV-1 $M^{pro}$.**

(A) Graphs showing the percentage change in BRET of DuProSense biosensor containing $M^{pro}$ cleavage sites with $M^{pro}$ measured from cell lysates of HEK293T cells expressing DuProSense biosensors containing $M^{pro}$ cleavage sites incubated with purified SARS-CoV-1 $M^{pro}$.

The green channel represents the $M^{pro}$ NSP4-5 sites and the red channel represents rest of the $M^{pro}$ cleavage sites.

(B, C) Graphs showing the half-time ($t_{1/2}$) (B) and cleavage rate (C) of all cleavage sites of $M^{pro}$ obtained from the kinetic sigmoidal model fitted the data obtained from in vitro assay using lysate of cells expressing DuProSense biosensors containing $M^{pro}$ and purified SARS-CoV-1 $M^{pro}$.

(D, E) Graphs showing the relative $t_{1/2}$ (D) and relative rate (E) of all substrate sites of $M^{pro}$ in comparison with the N-terminal auto-cleavage site.

Data shown are mean ± S.D. from three independent experiments.

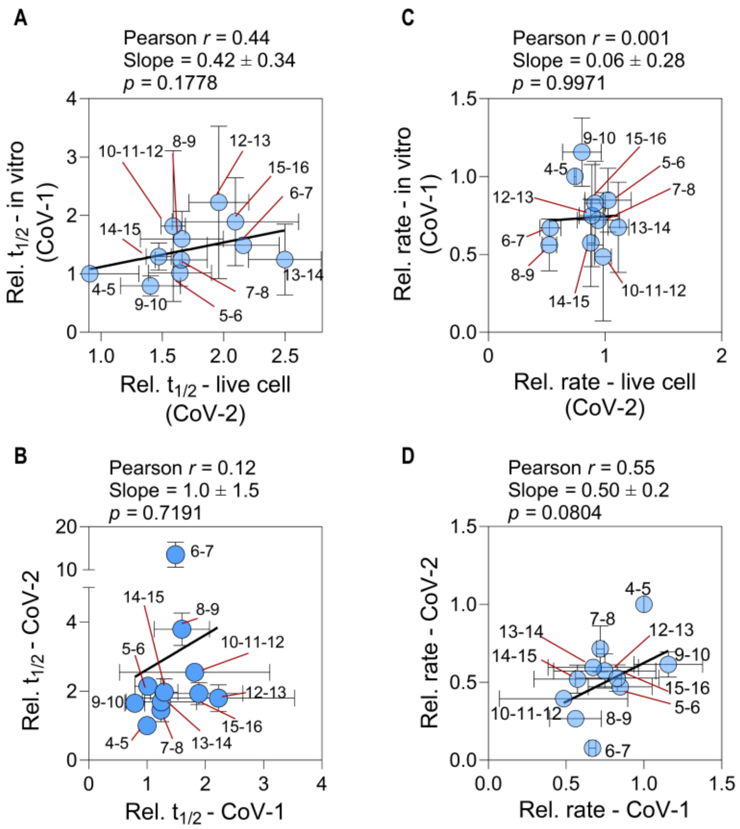


**Supporting Figure 13. Comparison of the cleavage kinetics of all cleavage sites in the presence of SARS-CoV-2 and SARS-CoV-1 $M^{pro}$.**

(A, C) Graphs showing the correlation between relative $t_{1/2}$ (A) and relative rates (B) obtained from the live cell assay in the presence of SARS-CoV-2 $M^{pro}$ is compared to in vitro assay in the presence of SAR-CoV-1 $M^{pro}$.
(B, D) Graphs showing the correlation between relative $t_{1/2}$ (B) and relative rate (D) obtained from in vitro assays with CoV-1 $M^{pro}$ and CoV-2 $M^{pro}$.
Data shown are mean ± S.D. from three independent experiments.

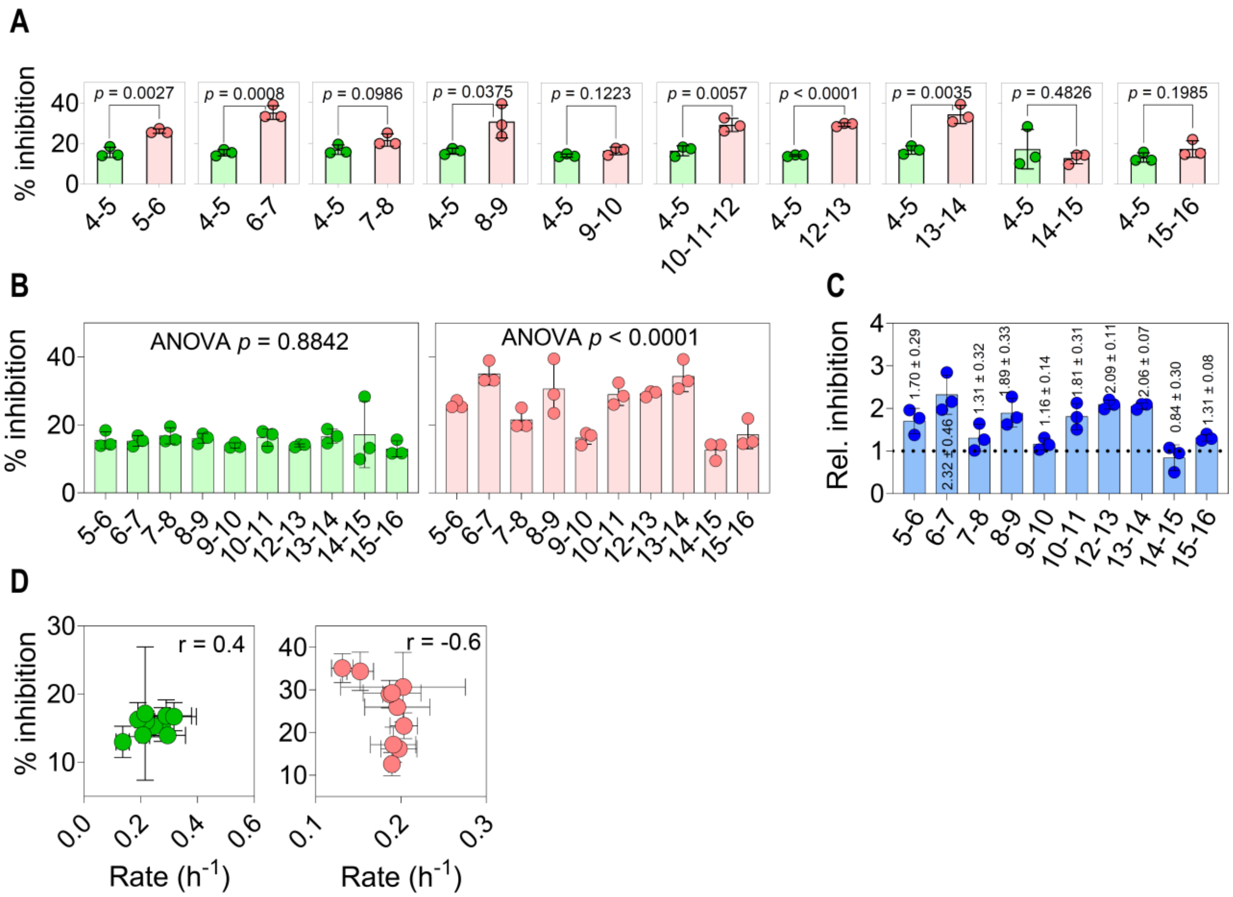


**Supporting Figure 14. DuProSense biosensor platform reveals cleavage site-specific M$^{pro}$ inhibitory potency of a single dose (4 μM) of nirmatrelvir in living cells.**

(A) Graphs showing % nirmatrelvir mediated M$^{pro}$ inhibition determined by the DuProSense biosensor containing M$^{pro}$ various cleavage sites expressed in living HEK293T cells in the presence of 4 μM nirmatrelvir measured after 22 h transfection.
(B) Graph showing the comparison of % M$^{pro}$ inhibition determined by the DuProSense biosensor containing M$^{pro}$ various cleavage sites in green (left panel) and red channel (right panel) of DuProSense biosensor expressed in living HEK293T cells in the presence of 4 μM nirmatrelvir measured after 22 h transfection.
(C) Graph showing the relative M$^{pro}$ inhibition determined by the DuProSense biosensors containing multiple sites relative to the respective NSP4-5.
(D) Graph showing the correlation between % inhibition and cleavage rates of the NSP4-5 of M$^{pro}$ (left panel) and that of other substrate sites (right panel).
Data shown are mean ± S.D. from three independent experiments, with each experiment performed in triplicates.
